# Childhood Trauma and Cortical Thickness in Bipolar Disorder with and without Psychosis: Effects by Diagnostic Group and Trauma Subtype

**DOI:** 10.64898/2026.10.13.758917

**Authors:** Diksha Chapagai, Grace Hodges, Abdulla Mutraji, Jessica A. Turner

## Abstract

**Background:** Bipolar disorder affects about 2-3% of the population and is associated with cortical abnormalities, with bipolar disorder with psychosis (BDP) reported to show more pronounced changes. It remains unclear whether childhood trauma is associated with cortical thickness differences across bipolar disorder subgroups, particularly in BDP.

**Methods:** Using the publicly available PARDIP (Psychosis and Affective Research Domains and Intermediate Phenotypes) data set, we examined whether childhood trauma was associated with cortical thickness differences among 120 individuals: 33 with bipolar disorder without psychosis (BD; 36% male), 38 with bipolar disorder with psychosis (BDP; 47% male), and 49 healthy controls (HC; 61% male). Participants completed the Childhood Trauma Questionnaire-Short Form (CTQ-SF), and T1-weighted MRI scans were processed with FreeSurfer to extract cortical thickness across the 68 Desikan-Killiany regions.

Linear mixed effects models tested associations between CTQ subtype scores and cortical thickness and trauma-by-diagnosis interactions, with sensitivity analyses for medication class and illness duration following the ENIGMA guidelines.

**Results:** No cortical thickness difference survived correction for multiple testing. At the nominal level, BD showed reduced thickness relative to HC in cingulate, frontal and occipital regions, whereas BDP showed greater superior temporal thickness relative to BD. Sexual abuse was nominally associated with reduced temporal thickness. Exploratory trauma-by-diagnosis interactions suggested region- and subtype-specific patterns that did not survive correction. Medication class and illness duration showed no significant effects after FDR correction.

**Conclusions:** These exploratory findings suggest nominal, region-specific associations between childhood trauma subtypes and cortical thickness in bipolar disorder. However, no findings survived correction for multiple comparisons, so they should be read as preliminary and need replication in larger samples.

## Introduction

Bipolar disorder affects approximately 2-3% of the population worldwide (1) and is associated with significant psychiatric disability and heterogeneity in illness course and severity. Structural magnetic resonance imaging (MRI) studies have repeatedly demonstrated cortical abnormalities in bipolar disorder, although the magnitude and regional specificity of these findings vary across studies (2,3). The ENIGMA Bipolar Disorder Working Group demonstrated abnormalities in bipolar disorder are region-specific rather than global, and consistently involve frontal, temporal, and insular cortices (2). Illness courses such as episode burden and manic severity have also been linked to progressive cortical changes, indicating that cortical thickness changes with illness progression (4). BDP has not consistently been distinguished from BD in structural imaging (5).

Environmental factors also play an important role in shaping heterogeneity within bipolar disorders. Childhood trauma is very prevalent among individuals with bipolar disorder and has been linked to earlier illness onset, increased suicidality, greater symptom severity, and an elevated chance of developing psychosis (6). In bipolar disorder, trauma exposure is associated with increased likelihood of developing psychotic symptoms, earlier emergence of psychosis, and more persistent illness (7). Emotional abuse and neglect show strong associations with psychosis risk (7). Neurodevelopmental models emphasize the importance of the type, timing, and the chronicity of trauma exposure, as early adversity may disrupt typical brain maturation processes (8).

Different forms of childhood adversity have been associated with distinct cortical thickness across brain regions (8,9). Emotional abuse and neglect show strong associations with medial prefrontal thinning (11). Evidence also indicates that trauma-related brain alterations overlap with regions implicated in psychosis, including in studies using broader measures of childhood adversity (8,11). Studies in individuals at ultra-high risk for psychosis have also shown that childhood trauma moderates cortical complexity and morphology in frontal and temporal regions (12,13).

Although much of this evidence comes from first-episode psychosis or ultra-high-risk samples rather than bipolar disorder, it raises the possibility that trauma-related cortical alterations are at least partly transdiagnostic. Significant gaps remain despite the growing evidence of linking trauma, brain structure, and psychosis. Few studies have tested trauma by-diagnosis interaction in bipolar disorder, and even fewer have examined trauma-related differences in brain structure. Addressing these gaps is essential for clarifying factors contributing to illness heterogeneity and identifying individuals at high risk for severe outcomes.

The present study examined cortical thickness across healthy controls (HC), BD, and BDP, to determine whether childhood trauma measures were associated with cortical thickness and assessed whether these associations differed by the presence of psychosis. Given the modest sample size and the number of regional comparisons, these analyses were considered exploratory. We hypothesized that BDP would show greater cortical thinning than BD. We further hypothesized that childhood trauma would have a stronger effect on cortical thickness in BDP compared to BD.

## Methods

### Data Source and Sample Characteristics

Data were obtained from the Psychosis and Affective Research Domains and Intermediate Phenotypes (PARDIP) dataset, accessed through the National Institute of Mental Health (NIMH) Data Archive (14). Participant recruitment and data collection are described in Bipolar Schizophrenia Network on Intermediate Phenotypes (BSNIP) studies (15). B-SNIP and PARDIP protocols were approved by local Institutional Review Boards across all participating study sites, and all participants provided written informed consent. The study was conducted in accordance with relevant institutional guidelines and regulations, including the Declaration of Helsinki. Diagnoses were established using the Structured Clinical Interview for DSM-IV Axis I Disorders, Patient Edition (SCID-I/P), based on consensus diagnostic conferences incorporating all available clinical information (16). Structural MRI data were collected across four different scanning sites using 3T scanners from Siemens, GE, and Philips platforms. The final analyzed sample comprised 120 adults; 33 BD, 38 BDP, and 49 HC. Of these, 69 (32 BD, 37 BDP) had complete medication data and 68 (31 BD, 37 BDP) illness duration data and were included in the corresponding sensitivity analyses.

## Measures

### Childhood Trauma

Childhood trauma was assessed using the Childhood Trauma Questionnaire-Short Form (CTQ-SF), which is a self-report measure used to evaluate exposure to adverse experiences in childhood and early adolescence (17). The CTQ assesses five trauma subtypes: emotional abuse, emotional neglect, physical abuse, physical neglect, and sexual abuse. The questionnaires are rated on a 5-point Likert scale and summed to generate each subscale score and a total trauma score for cumulative exposure.

Emotional abuse and emotional neglect were highly correlated in these data (r = .83); therefore, these subscales were averaged together, which is consistent with prior CTQ literature. CTQ scores were winsorized at the 1st and 99th percentiles to reduce the effect of extreme values.

### MRI Processing and Quality Control

All the T1-weighted structural MRI PARDIP data sets were processed using FreeSurfer (18), version 7.1, which generated cortical surface reconstruction, parcellation and vertex-wise cortical thickness estimates. Cortical thickness values were extracted for 68 regions defined by the Desikan-Killiany Atlas (18). The quality control procedures followed ENIGMA consortium standards as described previously (2). Scans with excessive movement or any segmentation errors (e.g., inaccurate cortical reconstruction or misclassification of white or grey matter), were excluded during quality control.

## Statistical Analysis Overview

Linear mixed effects models were used in each analysis. Age, sex, and intracranial volume (ICV) were included as fixed-effect covariates, and scanner site was included as a random effect to account for the multi-site data collection. All models applied false discovery rate (FDR) correction across the 68 cortical regions. Because no region survived FDR correction in any analysis, results are additionally reported at uncorrected threshold (p < .05 for diagnostic group and interaction comparisons and p < .01 for trauma subtype analyses) and are interpreted as exploratory only. Cortical thickness values were unavailable for a small number of participants in few regions after the quality control; the models were then fitted on complete cases. The analytic sample ranges from 113 to 120 participants across regions for diagnostic group comparisons for HC, from 112 to 119 for trauma and interaction models.

### Diagnostic Group Differences in Cortical Thickness

Diagnostic groups were compared in two ways. First, a three-group comparison (HC, BD, BDP) was used to examine cortical thickness differences. Second, BD and BDP were combined (BD + BDP) into a single bipolar disorder group and compared with HC to examine the overall diagnostic effect of bipolar disorder.

### Effects of Childhood Trauma Subtypes on Cortical Thickness

Models including each trauma subtype as a predictor were estimated across all 68 regions to determine whether trauma explained additional variance in cortical thickness beyond the diagnostic group effect. Scores for emotional abuse and neglect, physical abuse, physical neglect, and sexual abuse were included as predictors to assess the unique contribution of trauma subtype while controlling for diagnosis and covariates.

### Trauma-by-Diagnosis Interactions in Cortical Thickness

Linear mixed effects models were run with interaction terms between trauma subtype and diagnostic group while controlling for age, sex, ICV and scanner site (random effect).

### Sensitivity Analyses

Sensitivity analyses were conducted following the ENIGMA guidelines (2). Medication effects were tested within patients (n = 69; 32 BD, 37 BDP) using the linear mixed effects models with the medication classes (lithium, antiepileptics, antidepressants, and atypical antipsychotics) as binary predictors while also controlling for age, sex, ICV and scanner site. Illness duration was also examined using ENIGMA’s two-step hierarchical regression within patients with onset data (n = 68), with age, sex, ICV, and scanner entered in the first step and illness duration added in the second. Because illness duration was defined as current age minus age at onset, duration and age were strongly correlated in the subsample (r = 0.70, p < .001). All sensitivity analyses were FDR corrected.

## Results

### Sample Characteristics

Final sample characteristics are presented in Table 1. The diagnostic groups did not differ significantly in age (one-way ANOVA, p > .05) or sex distribution (χ² test, p > .05). Individuals with BD reported the highest level of childhood trauma across most of the CTQ subtypes whereas HC showed the lowest levels.

**Table 1.** Sample Characteristics by Diagnosis Group.

| Phenotypic Group | N | Age Range | % Male | MADRS | Emotional Abuse/Neglect | Physical Abuse | Physical Neglect | Sexual Abuse |
| --- | --- | --- | --- | --- | --- | --- | --- | --- |
| Healthy Controls (HC) | 49 | 19-60 | 61 | - | 7.79 (4.02) | 6.61 (3.32) | 6.00 (1.84) | 5.73 (2.26) |
| Bipolar without Psychosis (BD) | 33 | 19-59 | 36 | 12.69 (8.80) | 13.82 (5.58) | 10.64 (5.52) | 8.99 (3.52) | 9.88 (7.72) |
| Bipolar with Psychosis (BDP) | 38 | 19-59 | 47 | 15.17 (11.24) | 12.45 (5.18) | 8.86 (5.19) | 9.23 (3.50) | 8.14 (5.78) |
| Group Differences | | | $\chi^2(2) = 5.03$<br>$p = .081$ | | $F(2, 116) = 17.90$<br>$p < .001$ | $F(2, 116) = 7.73$<br>$p < .001$ | $F(2, 116) = 16.27$<br>$p < .001$ | $F(2, 116) = 6.09$<br>$p = .003$ |

### Diagnostic Differences in Cortical Thickness

No cortical thickness differences survived FDR correction for multiple comparisons. However, nominally significant results (P < .05, uncorrected) were identified across multiple regions shown in Table 2. Full cortical thickness results for all 68 regions for the comparisons of BD, BDP, and HC are reported in Appendix Table A1.

**Table 2.**
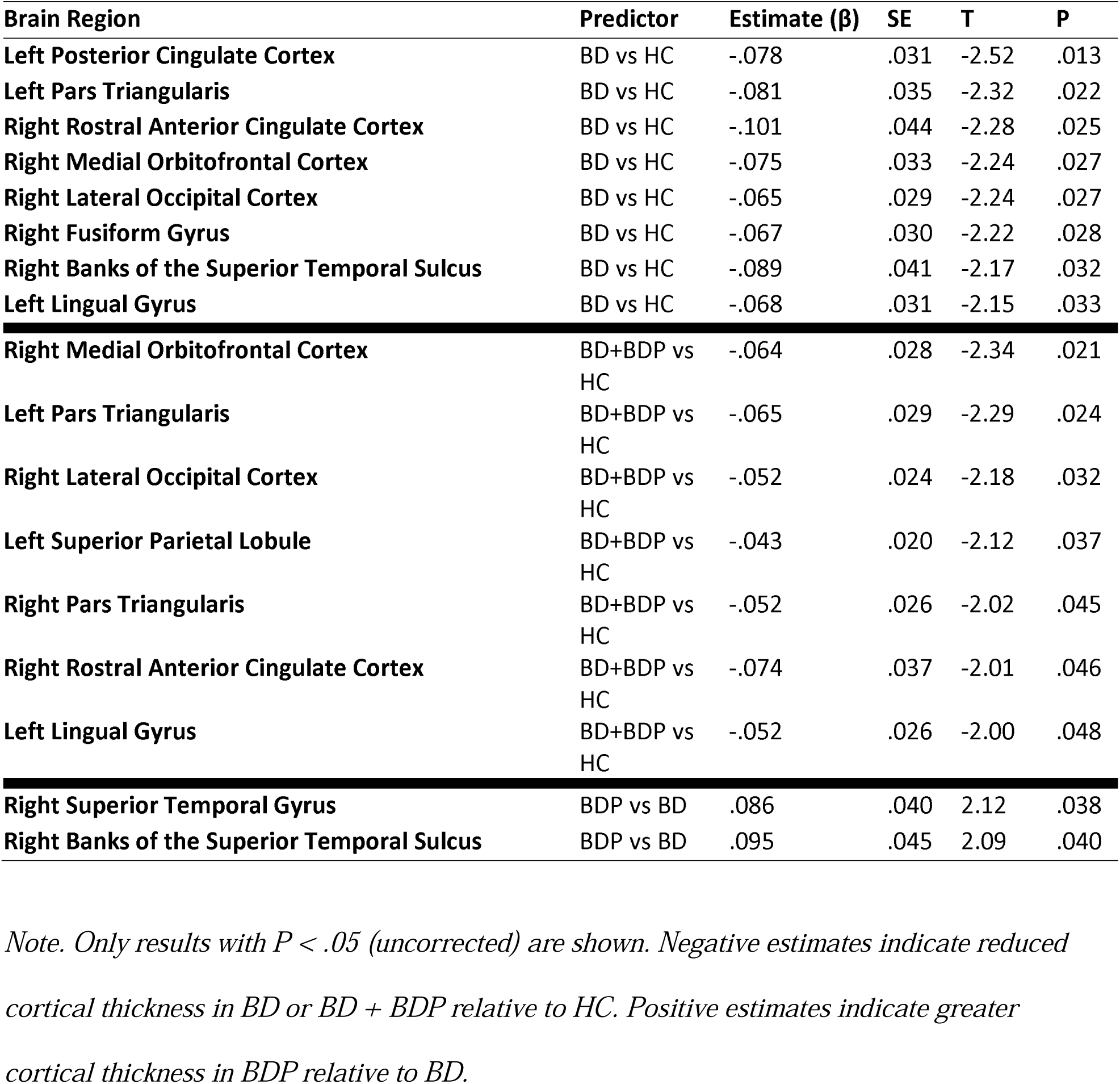
Cortical Thickness Differences Between Diagnostic Groups.

As seen in Figure 1, BDP generally had thicker cortices than BD in these data, but BD was thinner than HC. BDP did not show nominally significant cortical differences from HC, while BD + BDP combined showed the largest effect size of reduced cortical thickness relative to HC in the right medial orbitofrontal cortex, left pars triangularis, and right lateral occipital cortex. BDP showed nominally thicker cortex than BD in the right superior temporal gyrus and the banks of the superior temporal sulcus (STS).

**Figure 1.**
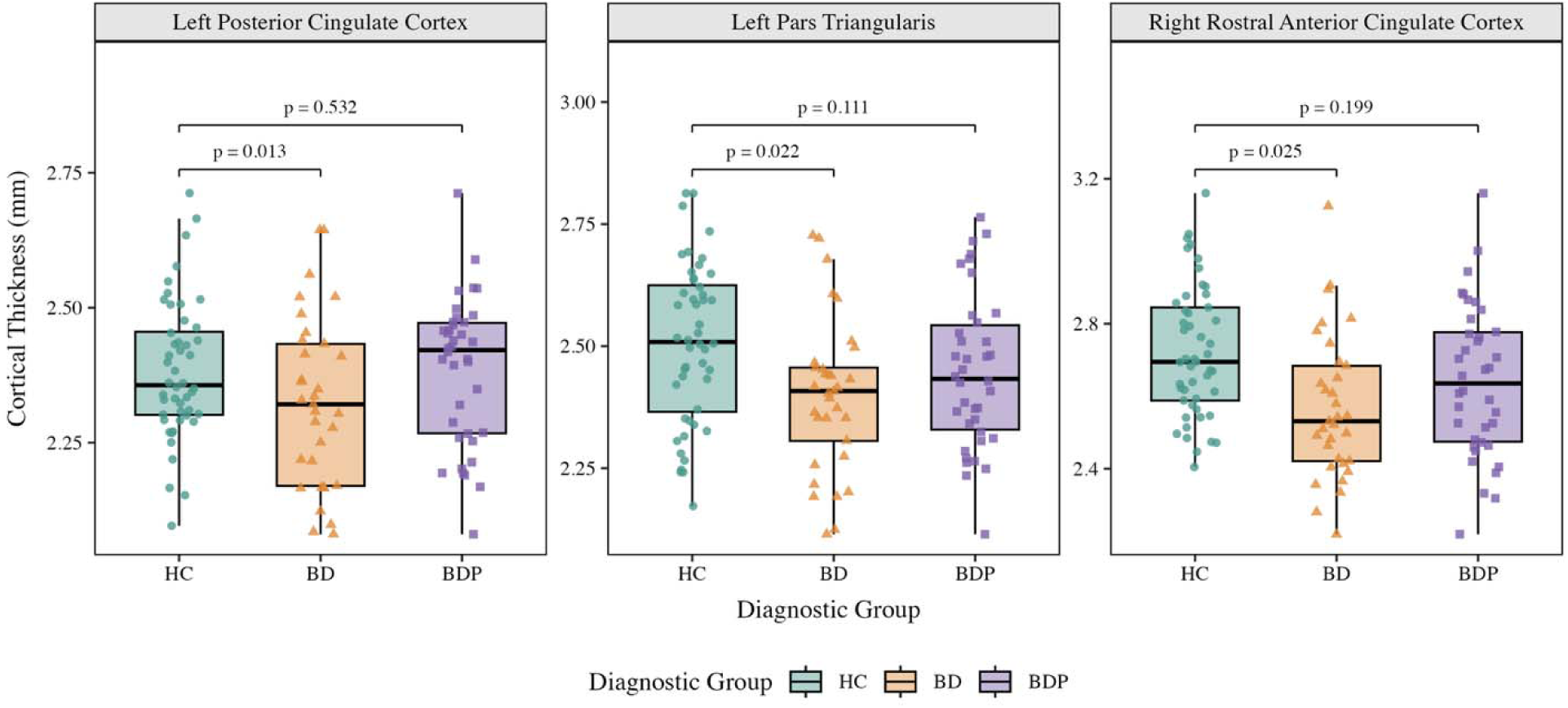
Diagnostic group differences in cortical thickness across BD, BDP, and HC. Cortical thickness values are shown for regions with nominally significant diagnostic group effects (p < .05, uncorrected).

### Effects of Childhood Trauma Subtypes on Cortical Thickness

Linear mixed effects models identified nominal (uncorrected) associations between childhood trauma exposure and cortical thickness across several regions after controlling diagnosis and covariates. Among trauma subtypes, sexual abuse showed the most associations, with reduced cortical thickness in the bilateral temporal poles, right superior temporal gyrus, bilateral postcentral gyri, bilateral supramarginal gyri, the left inferior parietal lobule, left precuneus and right medial orbitofrontal cortex. Physical abuse showed that the opposite pattern was nominally associated with greater cortical thickness in the left temporal pole, left entorhinal cortex, left superior temporal gyrus, and right transverse temporal gyrus.

Emotional abuse and neglect were nominally associated with reduced cortical thickness in the left temporal pole. Regions demonstrating a nominally significant association between childhood trauma and cortical thickness (p < .01) are presented in Table 3 and the three strongest associations are illustrated in Figure 2. Full results across all 68 regions are reported in Appendix Table A2.

**Figure 2.**
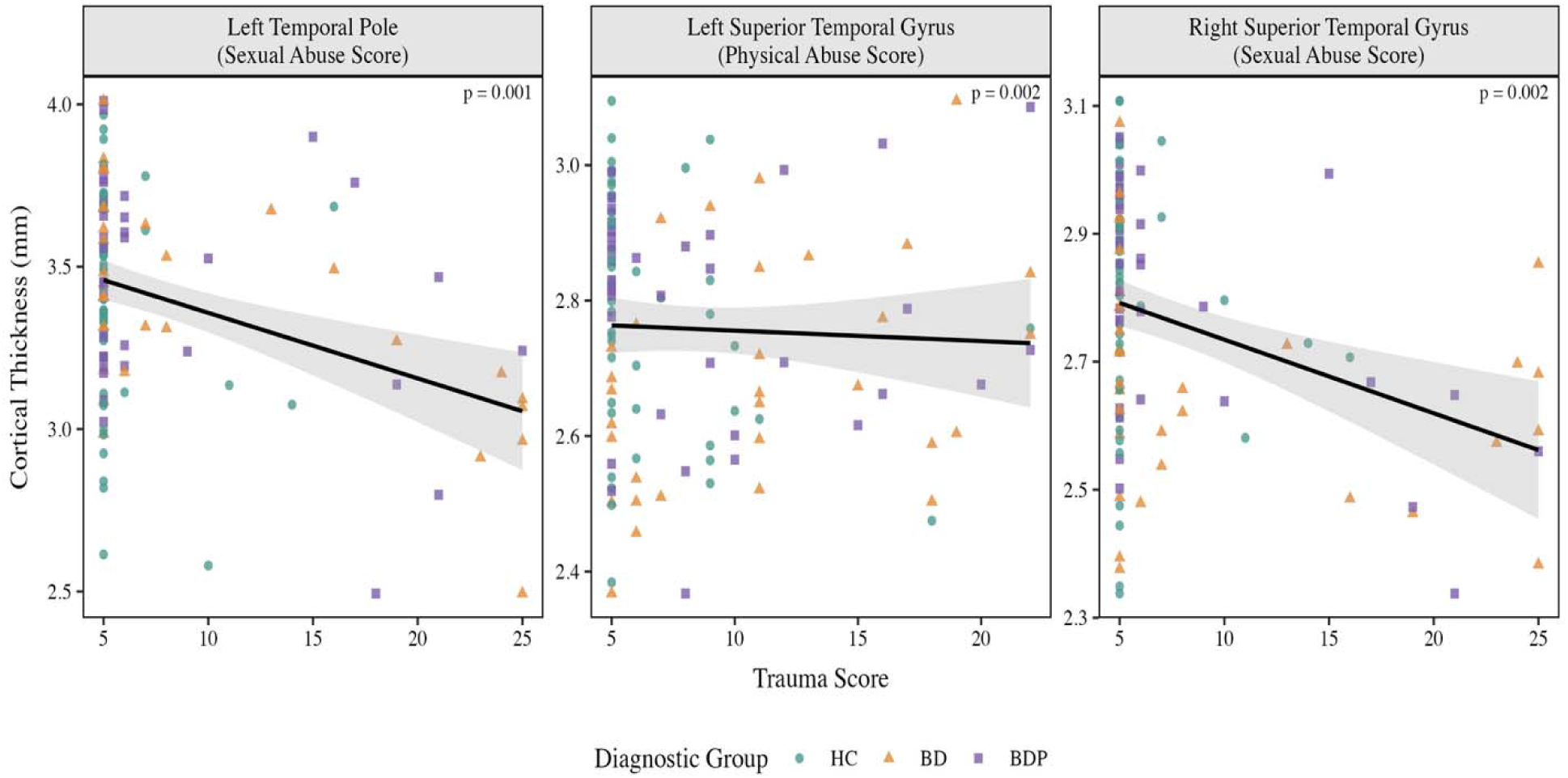
Main effects of childhood trauma on cortical thickness across representative brain regions.

**Table 3.** Childhood Trauma Subtypes and Cortical Thickness Across Brain Regions.

| Type of Abuse | Brain Region | Estimate ($\beta$) | P |
| --- | --- | --- | --- |
| Sexual Abuse | Left Temporal Pole | -.019 | .001 |
| Sexual Abuse | Right Superior Temporal Gyrus | -.010 | .002 |
| Sexual Abuse | Left Precuneus | -.007 | .002 |
| Sexual Abuse | Right Medial Orbitofrontal Cortex | -.009 | .003 |
| Sexual Abuse | Right Temporal Pole | -.019 | .003 |
| Sexual Abuse | Left Supramarginal Gyrus | -.008 | .003 |
| Sexual Abuse | Right Postcentral Gyrus | -.008 | .003 |
| Sexual Abuse | Left Inferior Parietal Lobule | -.007 | .003 |
| Sexual Abuse | Left Postcentral Gyrus | -.007 | .004 |
| Sexual Abuse | Right Supramarginal Gyrus | -.007 | .006 |
| Physical Abuse | Left Superior Temporal Gyrus | .013 | .002 |
| Physical Abuse | Right Transverse Temporal Gyrus | .018 | .004 |
| Physical Abuse | Left Temporal Pole | .022 | .007 |
| Physical Abuse | Left Entorhinal Cortex | .022 | .009 |
| Emotional Abuse and Neglect | Left Temporal Pole | -.024 | .005 |

*Note. Only associations with $P < .01$ (uncorrected) are shown. No regions survived FDR correction ( $P > .05$ ). Negative estimates indicate reduced cortical thickness, whereas positive estimates indicate increased thickness.*

### Trauma-by-Diagnosis Interactions in Cortical Thickness

When BD and BDP were separated, trauma effects differed across the diagnostic groups. Interaction effects between trauma subtype and diagnostic group are illustrated in Figures 3 – 6, and nominally significant interactions are presented in Table 4. Full interaction results across all 68 regions are reported in Appendix Table A3.

**Figure 3.**
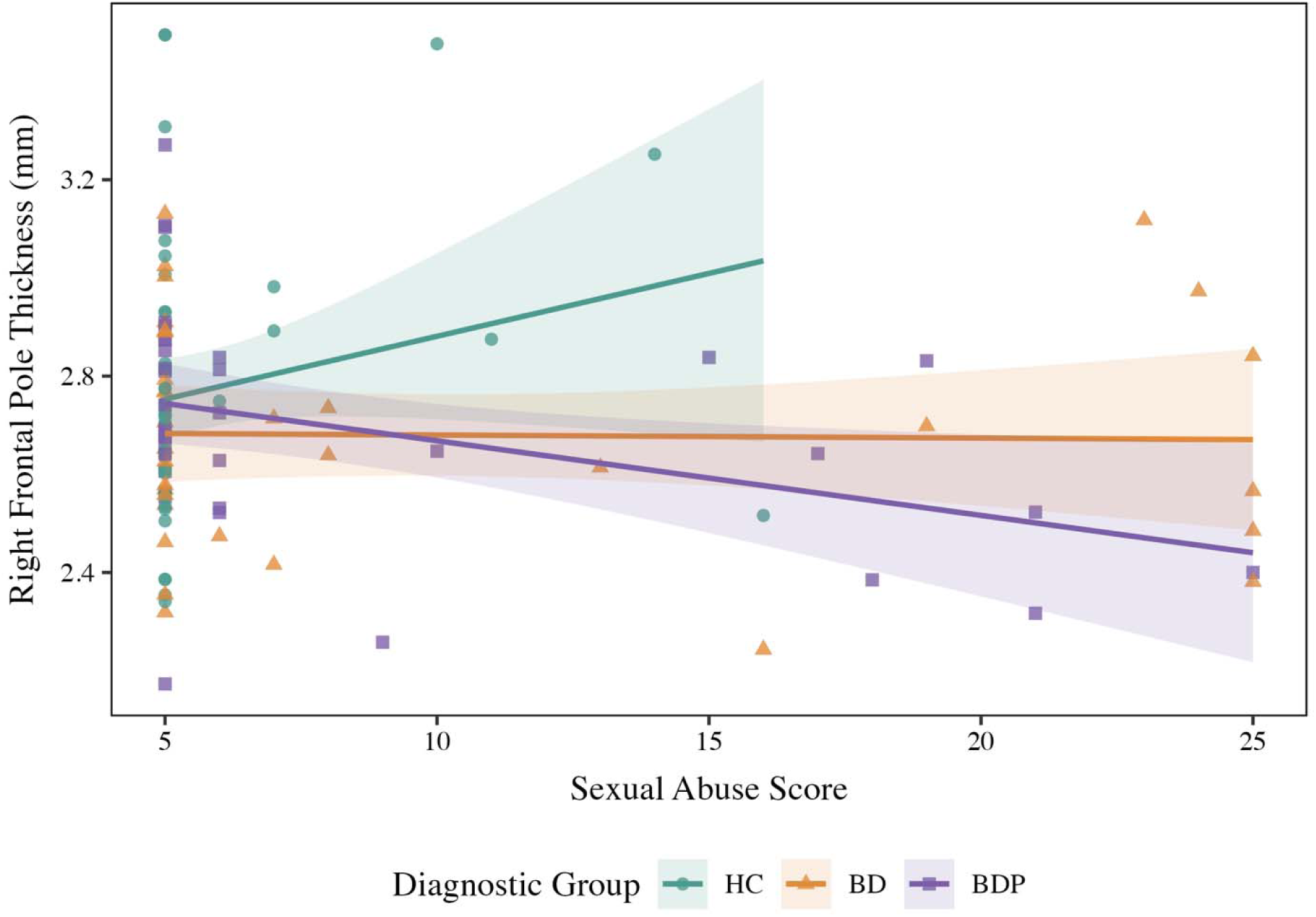
Association between sexual abuse scores and cortical thickness in the right frontal pole by diagnostic group. Lines represent group-specific regression slopes for HC, BD, and BDP.

**Figure 4.**
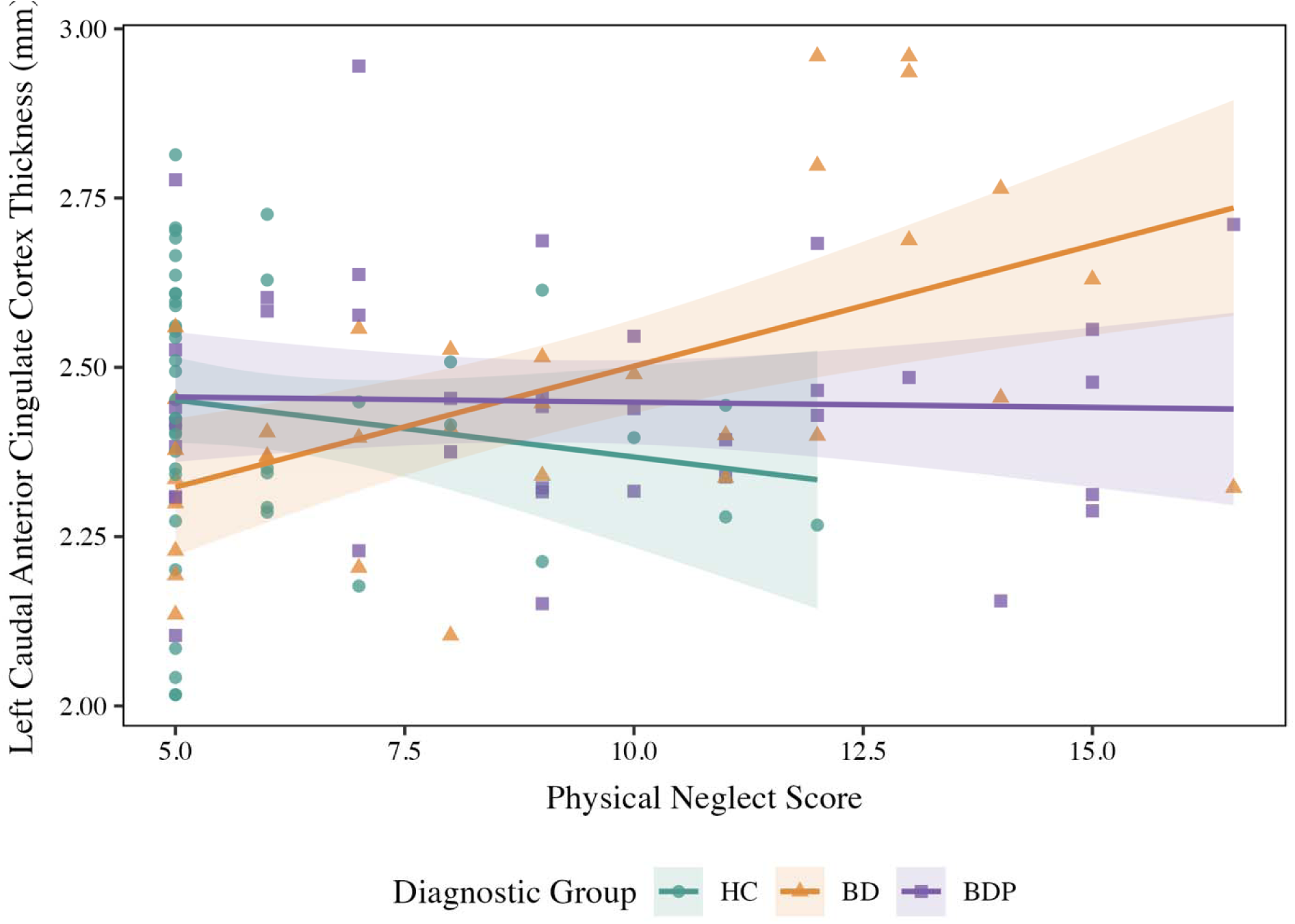
Association between physical neglect scores and cortical thickness in the left caudal anterior cingulate cortex by diagnostic group. Lines represent group-specific regression slopes for HC, BD, and BDP.

**Figure 5.**
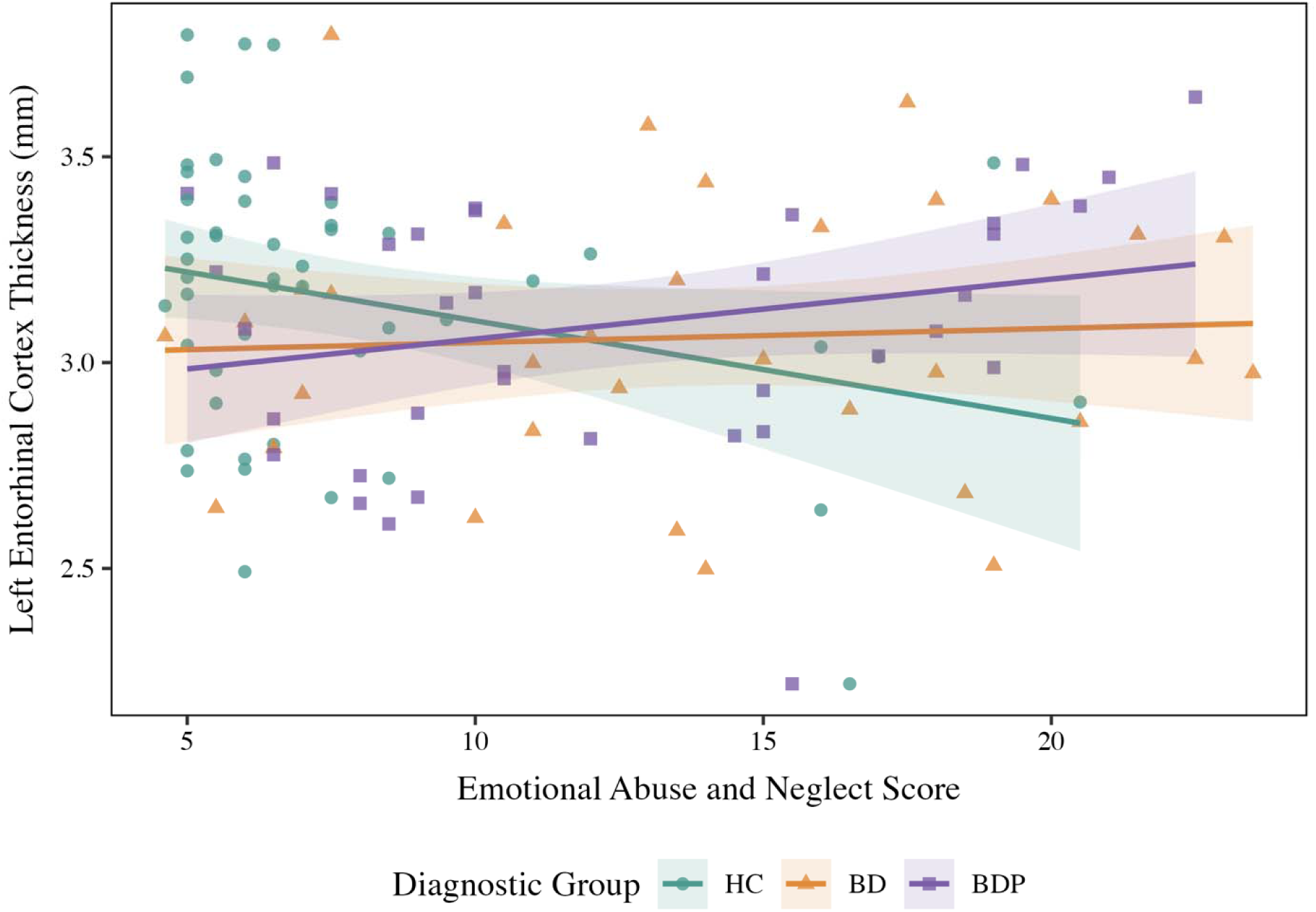
Association between emotional abuse and neglect scores and cortical thickness in the left entorhinal cortex by diagnostic group. Lines represent group-specific regression slopes for HC, BD, and BDP.

**Figure 6.**
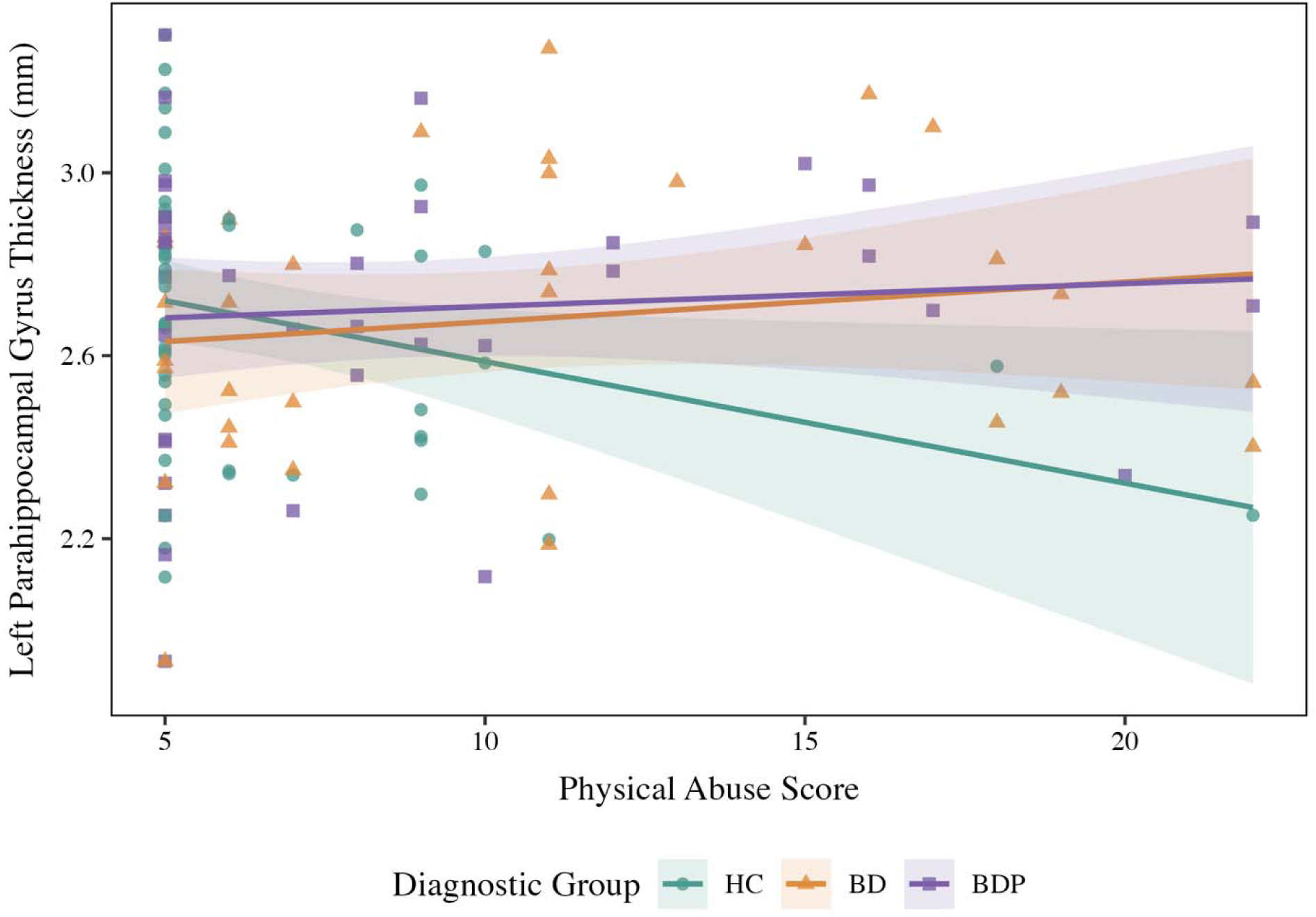
Association between physical abuse scores and cortical thickness in the left parahippocampal gyrus by diagnostic group. Lines represent group-specific regression slopes for HC, BD, and BDP.

**Table 4.** Trauma Subtype by Diagnostic Group Interaction Effects on Cortical Thickness.

| Brain Region | Trauma | Group | Estimate ($\beta$) | P |
| --- | --- | --- | --- | --- |
| Left Caudal Anterior Cingulate Cortex | Physical Neglect | BD | .078 | .001 |
| Right Isthmus of the Cingulate Cortex | Physical Neglect | BDP | -.043 | .010 |
| Left Entorhinal Cortex | Emotional Abuse and Neglect | BDP | .062 | .005 |
| Right Cuneus | Emotional Abuse and Neglect | BD | -.030 | .007 |
| Left Lateral Occipital Cortex | Emotional Abuse and Neglect | BD | -.023 | .010 |
| Left Parahippocampal Gyrus | Physical Abuse | BDP | .061 | .007 |

*Note. Interaction effects are shown for $P < .01$ (uncorrected). No regions survived FDR correction for multiple comparisons. Estimates reflect differences in trauma-thickness association for each group relative to HC.*

Physical neglect showed a positive association in left caudal anterior cingulate thickness in BD than HC (β = .078, p = .001), and a negative association with right isthmus cingulate thickness in BDP than in HC (β = −.043, p = .010).

Similarly, emotional abuse and neglect showed a positive association in the left entorhinal cortex in BDP relative to HC (β = .062, p = .005) and more negative association in BD relative to HC in the right cuneus (β = −.030, p = .007) and left lateral occipital cortex (β = −.023, p = .010). Physical abuse demonstrated group-specific patterns in the left parahippocampal gyrus in BDP relative to HC (β = .061, p = .007).

### Sensitivity Analyses

No medication class significantly predicted cortical thickness in any ROI after FDR correction. Additional models found no significant effect of lithium with BD or BDP separately, and no lithium by diagnosis interaction. Illness duration was also not significantly associated with cortical thickness in any ROI after FDR correction. The analyses were limited by subsample size (n = 69 for medication, n= 68 for illness duration) and a strong correlation between duration and age (r = .70); the null results do not exclude modest effects of medications or illness course.

## Discussion

This study examined how childhood trauma relates to the cortical thickness across individuals with HC, BD, and BDP, focusing on whether childhood trauma effects differed by diagnostic groups. The overall findings were that childhood trauma was nominally associated with region-specific differences in cortical thickness, although no effect survived correction for multiple comparisons. Contrary to the initial hypothesis, trauma effects were not consistently stronger in BDP relative to BD.

These findings suggest that childhood trauma is not merely a co-occurring factor of bipolar disorder but may be associated with variability in brain structure and illness severity. Consistent with prior research, nominal trauma-related effects were observed in regions such as the medial orbitofrontal cortex, temporal pole, and superior temporal gyrus (10). The same brain networks have been implicated in bipolar disorder and psychosis (2).

### Diagnostic Differences in Cortical Thickness

No cortical thickness differences survived FDR correction for multiple comparisons, but there were nominally significant findings indicating reduced cortical thickness in bipolar disorder, particularly in cingulate, frontal, and occipital regions such as the posterior cingulate cortex, rostral anterior cingulate cortex, medial orbitofrontal cortex, lateral occipital cortex, and pars triangularis. The findings are consistent with large neuroimaging studies from the ENIGMA consortium which have shown that cortical abnormalities in bipolar disorder are region-specific rather than global, particularly in the medial orbitofrontal cortex. (2)

Contrary to our hypothesis, BDP did not consistently show greater cortical thinning than BD relative to HC; instead, it showed minimal cortical thickness differences from HC. These analyses did not detect greater cortical thickness differences in BDP, which may be consistent with evidence that psychosis history in bipolar disorder is more strongly related to cortical surface area than to thickness (2).

### Effects of Childhood Trauma on Cortical Thickness

Childhood trauma was associated with cortical thickness across different brain regions while controlling diagnostic groups. Effects were region-specific with both increases and decreases in cortical thickness depending on the trauma subtype and brain region. Sexual abuse was associated with reduced cortical thickness in regions including the temporal pole and superior temporal gyrus, which are involved in emotional processing and social cognition (10). Physical abuse was associated with increased cortical thickness in regions such as superior temporal gyrus, transverse temporal gyrus, and entorhinal cortex. Emotional abuse and neglect were associated with reduced cortical thickness in the temporal pole. These effects may arise through stress-related neurobiological mechanisms such as prolonged HPA axis activation and synaptic loss (8)

Relatively more widespread effects were observed for sexual abuse, which are consistent with prior studies that show that these trauma types are particularly linked to the alteration in frontal and temporal regions, which are involved in emotional processing and regulation (10). When trauma-related effects were compared with diagnostic differences in cortical thickness, there was a partial overlap in affected brain regions, specifically in the right medial orbitofrontal cortex being thinner in bipolar disorder relative to controls and is associated with sexual abuse and the right superior temporal gyrus differing between BDP and BD and associated with sexual abuse.

### Trauma-by-Diagnosis Interactions

The interaction analyses indicate that nominal trauma-related associations with cortical thickness differed across diagnostic groups and varied by trauma subtype. Emotional abuse and neglect showed more widespread interaction effects in BD than BDP (7 vs 2 regions), as did physical abuse (2 vs 1 regions). In contrast, sexual abuse showed nominal interactions only in BDP (3 regions), and physical neglect showed more widespread effects in BDP than in BD (5 vs 2 regions). These findings indicate that trauma effects were not uniformly amplified in psychosis but appeared to depend on trauma subtypes. The lack of a consistently stronger effect in BDP also suggests that BDP may reflect a distinct neurobiological profile rather than simply a more severe form of bipolar disorder.

Crucially, the findings did not support the initial hypothesis that psychosis would amplify trauma effects on cortical thickness. In several regions, nominal trauma-related effects were weaker in BDP than in BD. This suggests that the presence of psychosis may not intensify bipolar-related cortical alterations but instead reflects a distinct neurobiological mechanism. Many of these regions, including the anterior cingulate cortex, parahippocampal gyrus, entorhinal cortex, and lateral occipital cortex are involved in emotional regulation, memory, and visual and social processing, suggesting that trauma-by-diagnosis effects may differ across these functional systems (10).

### Clinical and Theoretical Implications

Altogether, these findings highlight childhood trauma as a factor associated with heterogeneity in bipolar disorder. The regions implicated, such as the medial orbitofrontal cortex, temporal pole, and anterior cingulate cortex are involved in emotional regulation and cognitive control and may be associated with the earlier onset, more severe symptoms, and poorer functional outcomes seen in individuals with trauma exposure. The sensitivity analyses indicated that the nominal cortical thickness findings were not significantly associated with medication or illness duration after FDR correction. However, these analyses were underpowered, and illness duration was strongly collinear with age, so they should be read as failing to identify confounding rather than ruling it out.

### Limitations

Several additional limitations should be considered when interpreting these findings. First, the cross-sectional design precludes causal inferences regarding the temporal relationships among childhood trauma, psychosis, and cortical thickness. Second, childhood trauma was assessed with a retrospective self-report measure that may be vulnerable to recall bias and underreporting. Third, MRI data were collected across multiple sites using scanners from three vendors; although scanner sites were modeled as a random effect, this adjusts for mean differences, but residual variability may remain. Fourth, medication was modeled only as binary class indicators, without dose or duration, so residual confounding from treatment (association of lithium with greater cortical thickness has been reported in prior work although no medication effect was significant in the sample after FDR correction) cannot be excluded.

Fifth, the patient subgroups were modest in size, which limited statistical power, particularly for the trauma-by-diagnosis interaction analyses. Finally, group comparisons were interpreted at an uncorrected significance threshold, consistent with prior PARDIP and ENIGMA approaches, which increases sensitivity but also the risk of Type I error; accordingly, all nominal findings should be regarded as preliminary and require replication. Although the sensitivity analyses for medication and illness duration were null after FDR correction, the sensitivity subsample were small relative to ENIGMA (n = 69 for medication and n = 68 for illness duration, vs n = 1,837), which likely reduced power to detect such effects.

## Future Directions

Future research should use longitudinal neuroimaging designs to examine whether childhood trauma is associated with changes in cortical alterations over time, including whether trauma exposure relates to differences in the rate of cortical thinning and thickening. Incorporating stress-related biomarkers such as cortisol and other inflammatory markers may clarify biological mechanisms linking trauma to brain structure. In addition, examining the effects of psychotropic medication as well as distinguishing between specific trauma subtypes and timing of exposure may further clarify the biological mechanisms. Integrating functional and cognitive outcomes as well as evaluating trauma-informed interventions may help link structural findings to clinical trajectories in bipolar disorder.

## Conclusion

Overall, this study provides evidence that childhood trauma is associated with region-specific alterations in cortical thickness in bipolar disorder, and the effects differed across diagnostic groups and are not consistently stronger in BDP. The study also highlights the importance of trauma in understanding neurobiological heterogeneity and illness severity and underscores the importance of integrating environmental risk factors into bipolar disorder models.

## Acknowledgments

This work did not receive any specific grant from funding agencies in the public, commercial, or not-for-profit sectors. These analyses have not been previously presented or posted.

Data used in this study were obtained from the Bipolar-Schizophrenia Network on Intermediate Phenotypes (B-SNIP) and Psychosis and Affective Research Domains and Intermediate Phenotypes (PARDIP) consortia, in accordance with the terms of the data use agreement.

Data and/or research tools used in the preparation of this manuscript were obtained from the National Institute of Mental Health (NIMH) Data Archive (NDA). NDA is a collaborative informatics system created by the National Institutes of Health to provide a national resource to support and accelerate research in mental health. Dataset identifier(s): 2126. This manuscript reflects the views of the authors and may not reflect the opinions or views of the NIH or of the Submitters submitting original data to NDA.

We thank Jayden Morales, Cindy An, Urvi Wagh, and Sanjushree Karthikeyan for their assistance with quality control.

## Author Contributions (Credit)

Diksha Chapagai: Conceptualization, Methodology, Investigation, Formal Analysis, Writing original draft and editing.

Grace Hodges: Conceptualization, Formal Analysis, Review & Editing.

Abdulla Mutraji: Review, and editing.

Jessica Turner: Supervision, statistical discussions, results clarification, and writing support.

## Data Availability

Data used in the preparation of this manuscript were obtained from the National Institute of Mental Health (NIMH) Data Archive (NDA). NDA is a collaborative informatics system created by the National Institutes of Health to provide a national resource to support and accelerate research in mental health. Dataset identifier(s): ***2126.*** This manuscript reflects the views of the authors and may not reflect the opinions or views of the NIH or of the Submitters submitting original data to NDA.

## Declaration of Generative AI and AI-Assisted Technologies

During the preparation of this work, the author(s) used Claude (Anthropic) to improve language and clarity. After using this tool, the author(s) reviewed and edited the content as needed and takes full responsibility for the content of the publication.

## Appendix

**Appendix Table A1.**
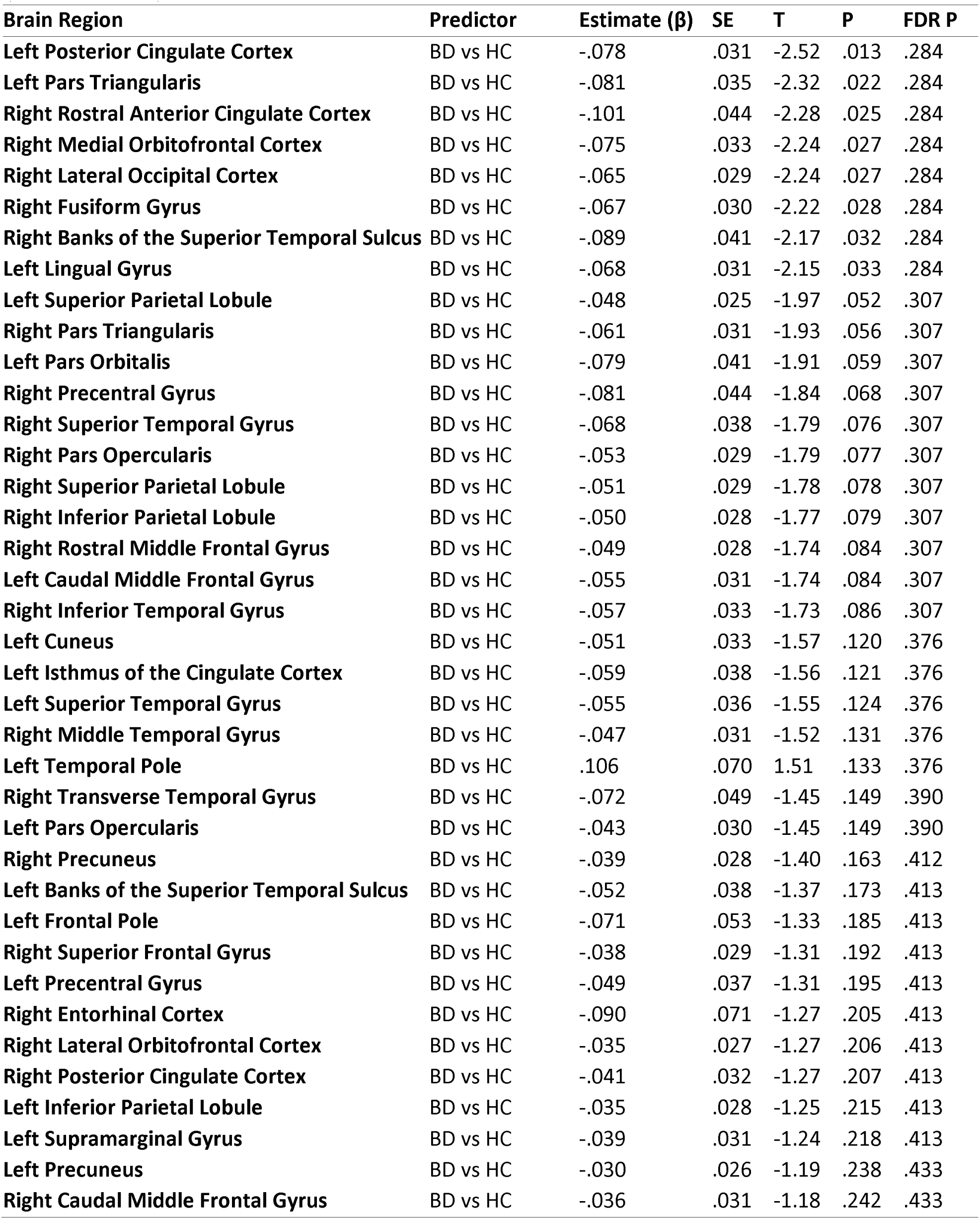
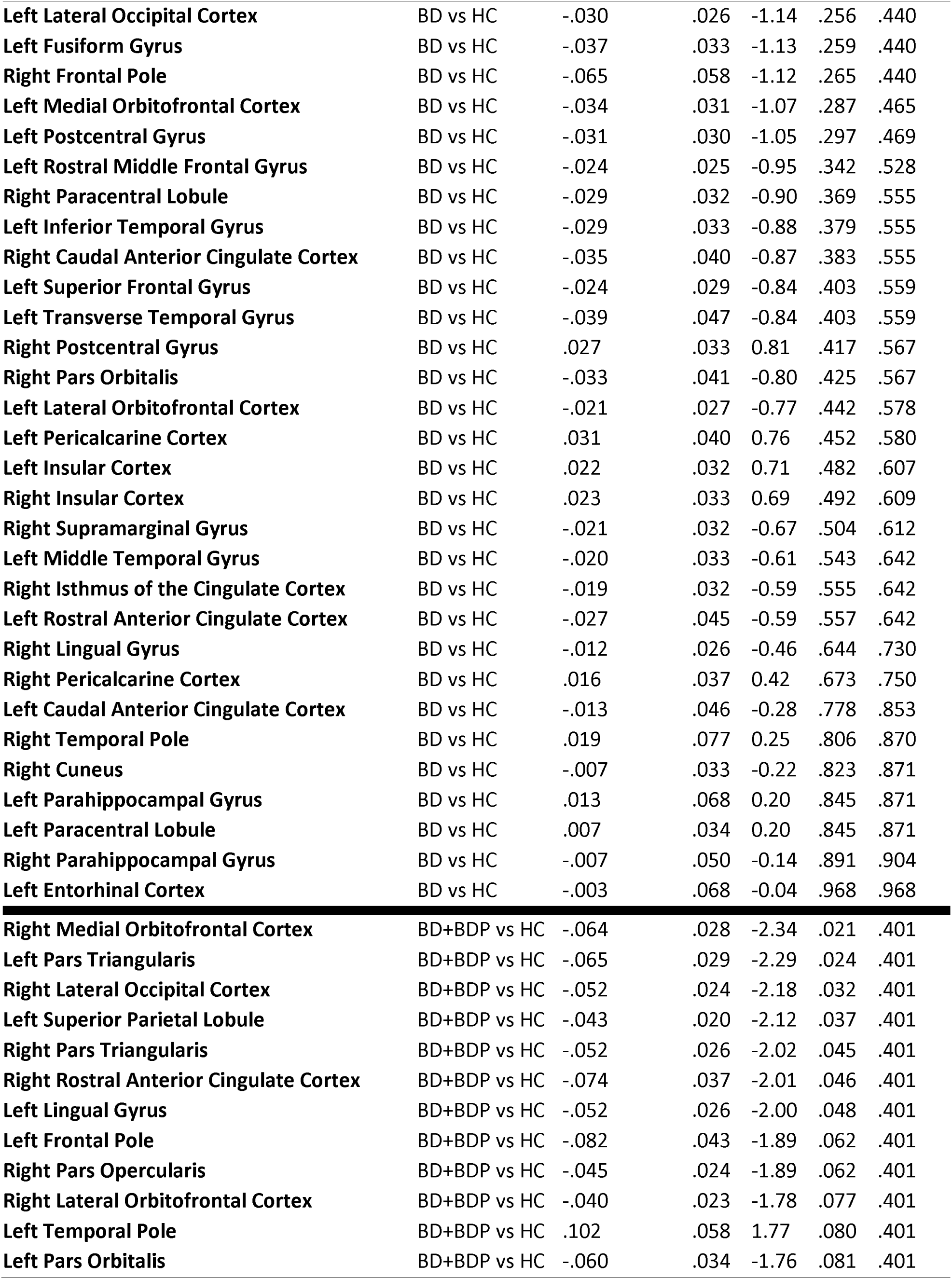
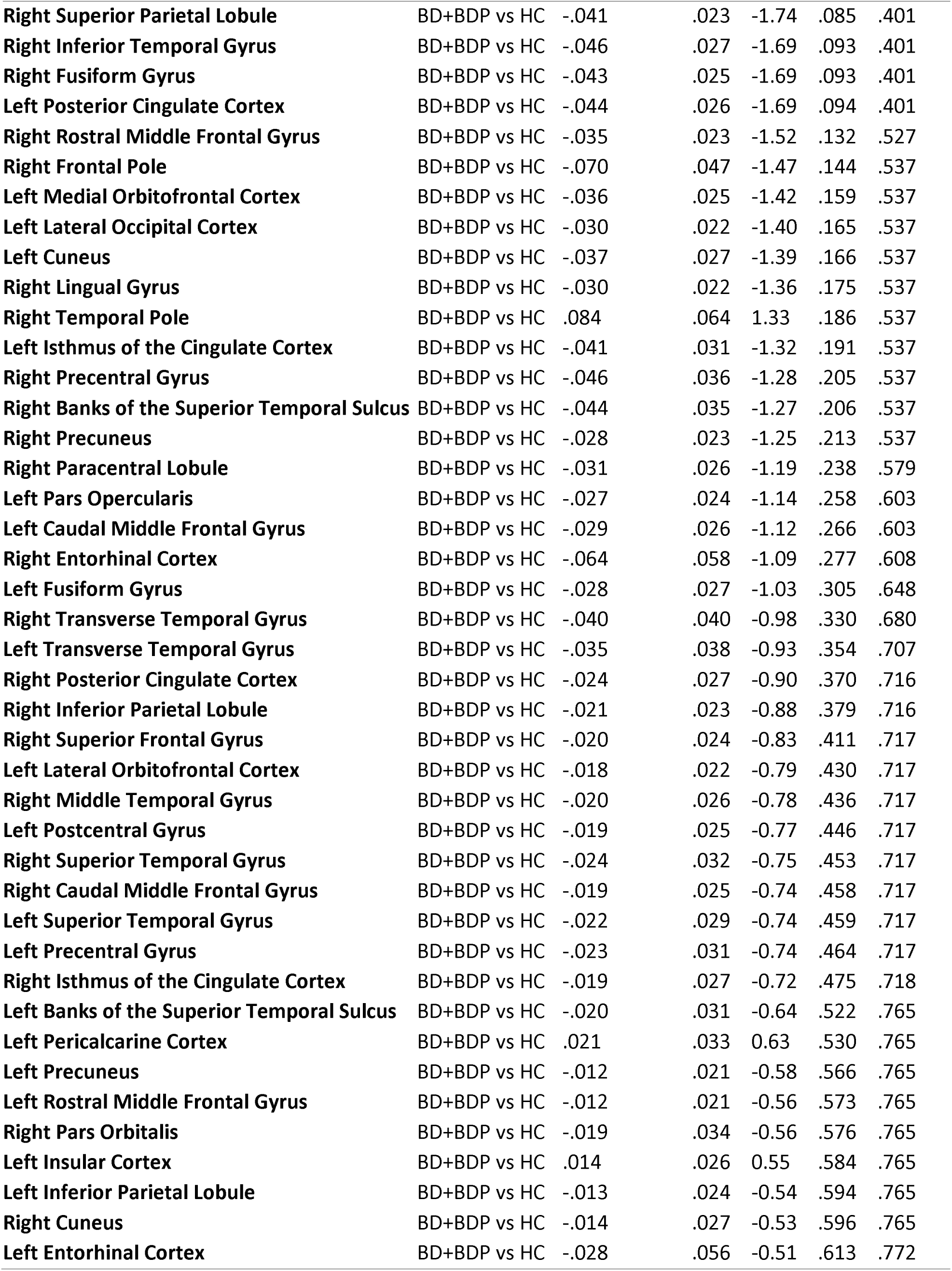
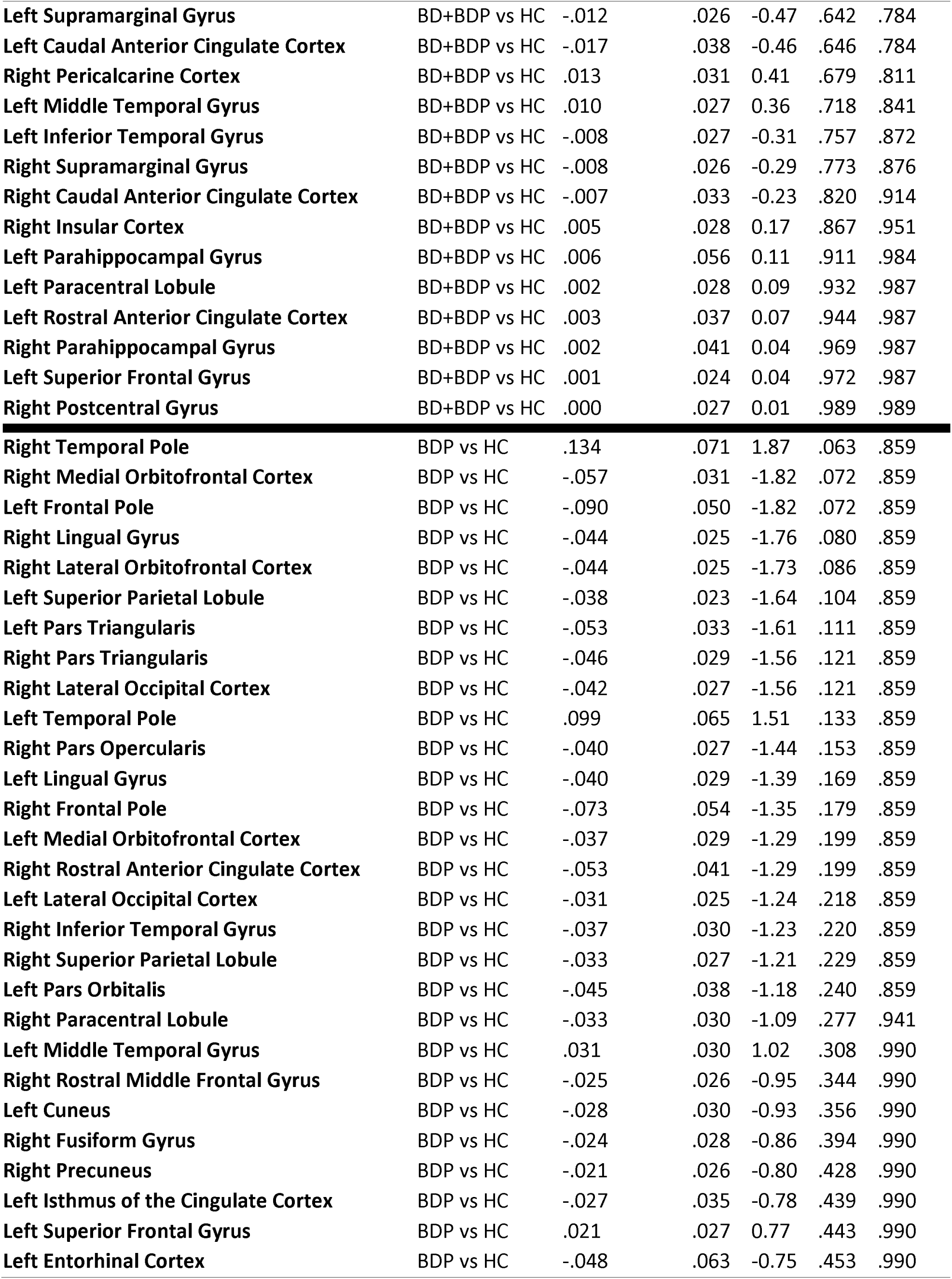
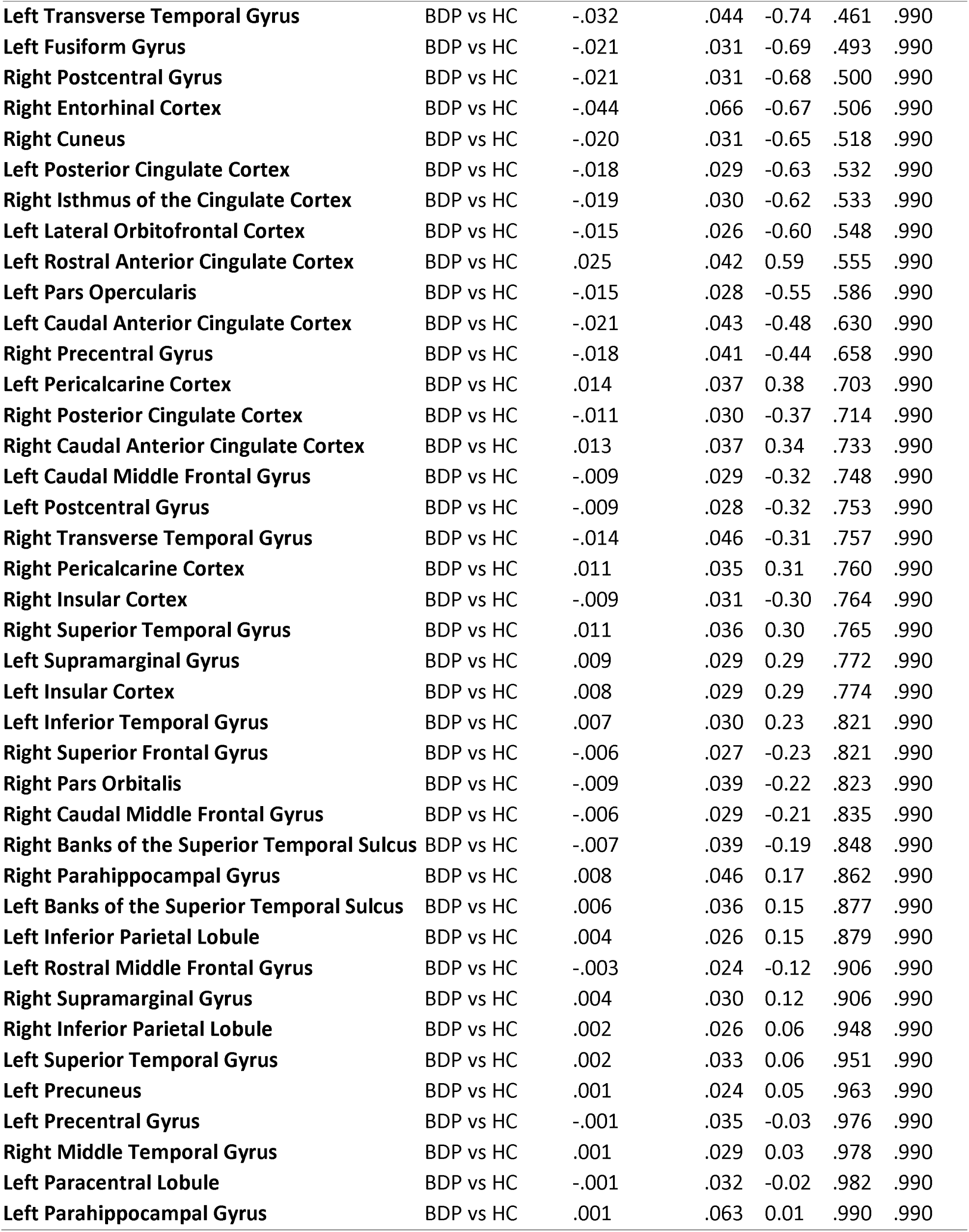
Cortical Thickness Across 68 Brain Regions by Diagnostic Comparisons (HC, BD, BDP).

| Brain Region | Predictor | Estimate ($\beta$) | SE | T | P | FDR P |
| --- | --- | --- | --- | --- | --- | --- |
| Left Posterior Cingulate Cortex | BD vs HC | -.078 | .031 | -2.52 | .013 | .284 |
| Left Pars Triangularis | BD vs HC | -.081 | .035 | -2.32 | .022 | .284 |
| Right Rostral Anterior Cingulate Cortex | BD vs HC | -.101 | .044 | -2.28 | .025 | .284 |
| Right Medial Orbitofrontal Cortex | BD vs HC | -.075 | .033 | -2.24 | .027 | .284 |
| Right Lateral Occipital Cortex | BD vs HC | -.065 | .029 | -2.24 | .027 | .284 |
| Right Fusiform Gyrus | BD vs HC | -.067 | .030 | -2.22 | .028 | .284 |
| Right Banks of the Superior Temporal Sulcus | BD vs HC | -.089 | .041 | -2.17 | .032 | .284 |
| Left Lingual Gyrus | BD vs HC | -.068 | .031 | -2.15 | .033 | .284 |
| Left Superior Parietal Lobule | BD vs HC | -.048 | .025 | -1.97 | .052 | .307 |
| Right Pars Triangularis | BD vs HC | -.061 | .031 | -1.93 | .056 | .307 |
| Left Pars Orbitalis | BD vs HC | -.079 | .041 | -1.91 | .059 | .307 |
| Right Precentral Gyrus | BD vs HC | -.081 | .044 | -1.84 | .068 | .307 |
| Right Superior Temporal Gyrus | BD vs HC | -.068 | .038 | -1.79 | .076 | .307 |
| Right Pars Opercularis | BD vs HC | -.053 | .029 | -1.79 | .077 | .307 |
| Right Superior Parietal Lobule | BD vs HC | -.051 | .029 | -1.78 | .078 | .307 |
| Right Inferior Parietal Lobule | BD vs HC | -.050 | .028 | -1.77 | .079 | .307 |
| Right Rostral Middle Frontal Gyrus | BD vs HC | -.049 | .028 | -1.74 | .084 | .307 |
| Left Caudal Middle Frontal Gyrus | BD vs HC | -.055 | .031 | -1.74 | .084 | .307 |
| Right Inferior Temporal Gyrus | BD vs HC | -.057 | .033 | -1.73 | .086 | .307 |
| Left Cuneus | BD vs HC | -.051 | .033 | -1.57 | .120 | .376 |
| Left Isthmus of the Cingulate Cortex | BD vs HC | -.059 | .038 | -1.56 | .121 | .376 |
| Left Superior Temporal Gyrus | BD vs HC | -.055 | .036 | -1.55 | .124 | .376 |
| Right Middle Temporal Gyrus | BD vs HC | -.047 | .031 | -1.52 | .131 | .376 |
| Left Temporal Pole | BD vs HC | .106 | .070 | 1.51 | .133 | .376 |
| Right Transverse Temporal Gyrus | BD vs HC | -.072 | .049 | -1.45 | .149 | .390 |
| Left Pars Opercularis | BD vs HC | -.043 | .030 | -1.45 | .149 | .390 |
| Right Precuneus | BD vs HC | -.039 | .028 | -1.40 | .163 | .412 |
| Left Banks of the Superior Temporal Sulcus | BD vs HC | -.052 | .038 | -1.37 | .173 | .413 |
| Left Frontal Pole | BD vs HC | -.071 | .053 | -1.33 | .185 | .413 |
| Right Superior Frontal Gyrus | BD vs HC | -.038 | .029 | -1.31 | .192 | .413 |
| Left Precentral Gyrus | BD vs HC | -.049 | .037 | -1.31 | .195 | .413 |
| Right Entorhinal Cortex | BD vs HC | -.090 | .071 | -1.27 | .205 | .413 |
| Right Lateral Orbitofrontal Cortex | BD vs HC | -.035 | .027 | -1.27 | .206 | .413 |
| Right Posterior Cingulate Cortex | BD vs HC | -.041 | .032 | -1.27 | .207 | .413 |
| Left Inferior Parietal Lobule | BD vs HC | -.035 | .028 | -1.25 | .215 | .413 |
| Left Supramarginal Gyrus | BD vs HC | -.039 | .031 | -1.24 | .218 | .413 |
| Left Precuneus | BD vs HC | -.030 | .026 | -1.19 | .238 | .433 |
| Right Caudal Middle Frontal Gyrus | BD vs HC | -.036 | .031 | -1.18 | .242 | .433 |
| **Left Lateral Occipital Cortex** | BD vs HC | -.030 | .026 | -1.14 | .256 | .440 |
| **Left Fusiform Gyrus** | BD vs HC | -.037 | .033 | -1.13 | .259 | .440 |
| **Right Frontal Pole** | BD vs HC | -.065 | .058 | -1.12 | .265 | .440 |
| **Left Medial Orbitofrontal Cortex** | BD vs HC | -.034 | .031 | -1.07 | .287 | .465 |
| **Left Postcentral Gyrus** | BD vs HC | -.031 | .030 | -1.05 | .297 | .469 |
| **Left Rostral Middle Frontal Gyrus** | BD vs HC | -.024 | .025 | -0.95 | .342 | .528 |
| **Right Paracentral Lobule** | BD vs HC | -.029 | .032 | -0.90 | .369 | .555 |
| **Left Inferior Temporal Gyrus** | BD vs HC | -.029 | .033 | -0.88 | .379 | .555 |
| **Right Caudal Anterior Cingulate Cortex** | BD vs HC | -.035 | .040 | -0.87 | .383 | .555 |
| **Left Superior Frontal Gyrus** | BD vs HC | -.024 | .029 | -0.84 | .403 | .559 |
| **Left Transverse Temporal Gyrus** | BD vs HC | -.039 | .047 | -0.84 | .403 | .559 |
| **Right Postcentral Gyrus** | BD vs HC | .027 | .033 | 0.81 | .417 | .567 |
| **Right Pars Orbitalis** | BD vs HC | -.033 | .041 | -0.80 | .425 | .567 |
| **Left Lateral Orbitofrontal Cortex** | BD vs HC | -.021 | .027 | -0.77 | .442 | .578 |
| **Left Pericalcarine Cortex** | BD vs HC | .031 | .040 | 0.76 | .452 | .580 |
| **Left Insular Cortex** | BD vs HC | .022 | .032 | 0.71 | .482 | .607 |
| **Right Insular Cortex** | BD vs HC | .023 | .033 | 0.69 | .492 | .609 |
| **Right Supramarginal Gyrus** | BD vs HC | -.021 | .032 | -0.67 | .504 | .612 |
| **Left Middle Temporal Gyrus** | BD vs HC | -.020 | .033 | -0.61 | .543 | .642 |
| **Right Isthmus of the Cingulate Cortex** | BD vs HC | -.019 | .032 | -0.59 | .555 | .642 |
| **Left Rostral Anterior Cingulate Cortex** | BD vs HC | -.027 | .045 | -0.59 | .557 | .642 |
| **Right Lingual Gyrus** | BD vs HC | -.012 | .026 | -0.46 | .644 | .730 |
| **Right Pericalcarine Cortex** | BD vs HC | .016 | .037 | 0.42 | .673 | .750 |
| **Left Caudal Anterior Cingulate Cortex** | BD vs HC | -.013 | .046 | -0.28 | .778 | .853 |
| **Right Temporal Pole** | BD vs HC | .019 | .077 | 0.25 | .806 | .870 |
| **Right Cuneus** | BD vs HC | -.007 | .033 | -0.22 | .823 | .871 |
| **Left Parahippocampal Gyrus** | BD vs HC | .013 | .068 | 0.20 | .845 | .871 |
| **Left Paracentral Lobule** | BD vs HC | .007 | .034 | 0.20 | .845 | .871 |
| **Right Parahippocampal Gyrus** | BD vs HC | -.007 | .050 | -0.14 | .891 | .904 |
| **Left Entorhinal Cortex** | BD vs HC | -.003 | .068 | -0.04 | .968 | .968 |
| **Right Medial Orbitofrontal Cortex** | BD+BDP vs HC | -.064 | .028 | -2.34 | .021 | .401 |
| **Left Pars Triangularis** | BD+BDP vs HC | -.065 | .029 | -2.29 | .024 | .401 |
| **Right Lateral Occipital Cortex** | BD+BDP vs HC | -.052 | .024 | -2.18 | .032 | .401 |
| **Left Superior Parietal Lobule** | BD+BDP vs HC | -.043 | .020 | -2.12 | .037 | .401 |
| **Right Pars Triangularis** | BD+BDP vs HC | -.052 | .026 | -2.02 | .045 | .401 |
| **Right Rostral Anterior Cingulate Cortex** | BD+BDP vs HC | -.074 | .037 | -2.01 | .046 | .401 |
| **Left Lingual Gyrus** | BD+BDP vs HC | -.052 | .026 | -2.00 | .048 | .401 |
| **Left Frontal Pole** | BD+BDP vs HC | -.082 | .043 | -1.89 | .062 | .401 |
| **Right Pars Opercularis** | BD+BDP vs HC | -.045 | .024 | -1.89 | .062 | .401 |
| **Right Lateral Orbitofrontal Cortex** | BD+BDP vs HC | -.040 | .023 | -1.78 | .077 | .401 |
| **Left Temporal Pole** | BD+BDP vs HC | .102 | .058 | 1.77 | .080 | .401 |
| **Left Pars Orbitalis** | BD+BDP vs HC | -.060 | .034 | -1.76 | .081 | .401 |
| **Right Superior Parietal Lobule** | BD+BDP vs HC | -.041 | .023 | -1.74 | .085 | .401 |
| **Right Inferior Temporal Gyrus** | BD+BDP vs HC | -.046 | .027 | -1.69 | .093 | .401 |
| **Right Fusiform Gyrus** | BD+BDP vs HC | -.043 | .025 | -1.69 | .093 | .401 |
| **Left Posterior Cingulate Cortex** | BD+BDP vs HC | -.044 | .026 | -1.69 | .094 | .401 |
| **Right Rostral Middle Frontal Gyrus** | BD+BDP vs HC | -.035 | .023 | -1.52 | .132 | .527 |
| **Right Frontal Pole** | BD+BDP vs HC | -.070 | .047 | -1.47 | .144 | .537 |
| **Left Medial Orbitofrontal Cortex** | BD+BDP vs HC | -.036 | .025 | -1.42 | .159 | .537 |
| **Left Lateral Occipital Cortex** | BD+BDP vs HC | -.030 | .022 | -1.40 | .165 | .537 |
| **Left Cuneus** | BD+BDP vs HC | -.037 | .027 | -1.39 | .166 | .537 |
| **Right Lingual Gyrus** | BD+BDP vs HC | -.030 | .022 | -1.36 | .175 | .537 |
| **Right Temporal Pole** | BD+BDP vs HC | .084 | .064 | 1.33 | .186 | .537 |
| **Left Isthmus of the Cingulate Cortex** | BD+BDP vs HC | -.041 | .031 | -1.32 | .191 | .537 |
| **Right Precentral Gyrus** | BD+BDP vs HC | -.046 | .036 | -1.28 | .205 | .537 |
| **Right Banks of the Superior Temporal Sulcus** | BD+BDP vs HC | -.044 | .035 | -1.27 | .206 | .537 |
| **Right Precuneus** | BD+BDP vs HC | -.028 | .023 | -1.25 | .213 | .537 |
| **Right Paracentral Lobule** | BD+BDP vs HC | -.031 | .026 | -1.19 | .238 | .579 |
| **Left Pars Opercularis** | BD+BDP vs HC | -.027 | .024 | -1.14 | .258 | .603 |
| **Left Caudal Middle Frontal Gyrus** | BD+BDP vs HC | -.029 | .026 | -1.12 | .266 | .603 |
| **Right Entorhinal Cortex** | BD+BDP vs HC | -.064 | .058 | -1.09 | .277 | .608 |
| **Left Fusiform Gyrus** | BD+BDP vs HC | -.028 | .027 | -1.03 | .305 | .648 |
| **Right Transverse Temporal Gyrus** | BD+BDP vs HC | -.040 | .040 | -0.98 | .330 | .680 |
| **Left Transverse Temporal Gyrus** | BD+BDP vs HC | -.035 | .038 | -0.93 | .354 | .707 |
| **Right Posterior Cingulate Cortex** | BD+BDP vs HC | -.024 | .027 | -0.90 | .370 | .716 |
| **Right Inferior Parietal Lobule** | BD+BDP vs HC | -.021 | .023 | -0.88 | .379 | .716 |
| **Right Superior Frontal Gyrus** | BD+BDP vs HC | -.020 | .024 | -0.83 | .411 | .717 |
| **Left Lateral Orbitofrontal Cortex** | BD+BDP vs HC | -.018 | .022 | -0.79 | .430 | .717 |
| **Right Middle Temporal Gyrus** | BD+BDP vs HC | -.020 | .026 | -0.78 | .436 | .717 |
| **Left Postcentral Gyrus** | BD+BDP vs HC | -.019 | .025 | -0.77 | .446 | .717 |
| **Right Superior Temporal Gyrus** | BD+BDP vs HC | -.024 | .032 | -0.75 | .453 | .717 |
| **Right Caudal Middle Frontal Gyrus** | BD+BDP vs HC | -.019 | .025 | -0.74 | .458 | .717 |
| **Left Superior Temporal Gyrus** | BD+BDP vs HC | -.022 | .029 | -0.74 | .459 | .717 |
| **Left Precentral Gyrus** | BD+BDP vs HC | -.023 | .031 | -0.74 | .464 | .717 |
| **Right Isthmus of the Cingulate Cortex** | BD+BDP vs HC | -.019 | .027 | -0.72 | .475 | .718 |
| **Left Banks of the Superior Temporal Sulcus** | BD+BDP vs HC | -.020 | .031 | -0.64 | .522 | .765 |
| **Left Pericalcarine Cortex** | BD+BDP vs HC | .021 | .033 | 0.63 | .530 | .765 |
| **Left Precuneus** | BD+BDP vs HC | -.012 | .021 | -0.58 | .566 | .765 |
| **Left Rostral Middle Frontal Gyrus** | BD+BDP vs HC | -.012 | .021 | -0.56 | .573 | .765 |
| **Right Pars Orbitalis** | BD+BDP vs HC | -.019 | .034 | -0.56 | .576 | .765 |
| **Left Insular Cortex** | BD+BDP vs HC | .014 | .026 | 0.55 | .584 | .765 |
| **Left Inferior Parietal Lobule** | BD+BDP vs HC | -.013 | .024 | -0.54 | .594 | .765 |
| **Right Cuneus** | BD+BDP vs HC | -.014 | .027 | -0.53 | .596 | .765 |
| **Left Entorhinal Cortex** | BD+BDP vs HC | -.028 | .056 | -0.51 | .613 | .772 |
| **Left Supramarginal Gyrus** | BD+BDP vs HC | -.012 | .026 | -0.47 | .642 | .784 |
| **Left Caudal Anterior Cingulate Cortex** | BD+BDP vs HC | -.017 | .038 | -0.46 | .646 | .784 |
| **Right Pericalcarine Cortex** | BD+BDP vs HC | .013 | .031 | 0.41 | .679 | .811 |
| **Left Middle Temporal Gyrus** | BD+BDP vs HC | .010 | .027 | 0.36 | .718 | .841 |
| **Left Inferior Temporal Gyrus** | BD+BDP vs HC | -.008 | .027 | -0.31 | .757 | .872 |
| **Right Supramarginal Gyrus** | BD+BDP vs HC | -.008 | .026 | -0.29 | .773 | .876 |
| **Right Caudal Anterior Cingulate Cortex** | BD+BDP vs HC | -.007 | .033 | -0.23 | .820 | .914 |
| **Right Insular Cortex** | BD+BDP vs HC | .005 | .028 | 0.17 | .867 | .951 |
| **Left Parahippocampal Gyrus** | BD+BDP vs HC | .006 | .056 | 0.11 | .911 | .984 |
| **Left Paracentral Lobule** | BD+BDP vs HC | .002 | .028 | 0.09 | .932 | .987 |
| **Left Rostral Anterior Cingulate Cortex** | BD+BDP vs HC | .003 | .037 | 0.07 | .944 | .987 |
| **Right Parahippocampal Gyrus** | BD+BDP vs HC | .002 | .041 | 0.04 | .969 | .987 |
| **Left Superior Frontal Gyrus** | BD+BDP vs HC | .001 | .024 | 0.04 | .972 | .987 |
| **Right Postcentral Gyrus** | BD+BDP vs HC | .000 | .027 | 0.01 | .989 | .989 |
| **Right Temporal Pole** | BDP vs HC | .134 | .071 | 1.87 | .063 | .859 |
| **Right Medial Orbitofrontal Cortex** | BDP vs HC | -.057 | .031 | -1.82 | .072 | .859 |
| **Left Frontal Pole** | BDP vs HC | -.090 | .050 | -1.82 | .072 | .859 |
| **Right Lingual Gyrus** | BDP vs HC | -.044 | .025 | -1.76 | .080 | .859 |
| **Right Lateral Orbitofrontal Cortex** | BDP vs HC | -.044 | .025 | -1.73 | .086 | .859 |
| **Left Superior Parietal Lobule** | BDP vs HC | -.038 | .023 | -1.64 | .104 | .859 |
| **Left Pars Triangularis** | BDP vs HC | -.053 | .033 | -1.61 | .111 | .859 |
| **Right Pars Triangularis** | BDP vs HC | -.046 | .029 | -1.56 | .121 | .859 |
| **Right Lateral Occipital Cortex** | BDP vs HC | -.042 | .027 | -1.56 | .121 | .859 |
| **Left Temporal Pole** | BDP vs HC | .099 | .065 | 1.51 | .133 | .859 |
| **Right Pars Opercularis** | BDP vs HC | -.040 | .027 | -1.44 | .153 | .859 |
| **Left Lingual Gyrus** | BDP vs HC | -.040 | .029 | -1.39 | .169 | .859 |
| **Right Frontal Pole** | BDP vs HC | -.073 | .054 | -1.35 | .179 | .859 |
| **Left Medial Orbitofrontal Cortex** | BDP vs HC | -.037 | .029 | -1.29 | .199 | .859 |
| **Right Rostral Anterior Cingulate Cortex** | BDP vs HC | -.053 | .041 | -1.29 | .199 | .859 |
| **Left Lateral Occipital Cortex** | BDP vs HC | -.031 | .025 | -1.24 | .218 | .859 |
| **Right Inferior Temporal Gyrus** | BDP vs HC | -.037 | .030 | -1.23 | .220 | .859 |
| **Right Superior Parietal Lobule** | BDP vs HC | -.033 | .027 | -1.21 | .229 | .859 |
| **Left Pars Orbitalis** | BDP vs HC | -.045 | .038 | -1.18 | .240 | .859 |
| **Right Paracentral Lobule** | BDP vs HC | -.033 | .030 | -1.09 | .277 | .941 |
| **Left Middle Temporal Gyrus** | BDP vs HC | .031 | .030 | 1.02 | .308 | .990 |
| **Right Rostral Middle Frontal Gyrus** | BDP vs HC | -.025 | .026 | -0.95 | .344 | .990 |
| **Left Cuneus** | BDP vs HC | -.028 | .030 | -0.93 | .356 | .990 |
| **Right Fusiform Gyrus** | BDP vs HC | -.024 | .028 | -0.86 | .394 | .990 |
| **Right Precuneus** | BDP vs HC | -.021 | .026 | -0.80 | .428 | .990 |
| **Left Isthmus of the Cingulate Cortex** | BDP vs HC | -.027 | .035 | -0.78 | .439 | .990 |
| **Left Superior Frontal Gyrus** | BDP vs HC | .021 | .027 | 0.77 | .443 | .990 |
| **Left Entorhinal Cortex** | BDP vs HC | -.048 | .063 | -0.75 | .453 | .990 |
| **Left Transverse Temporal Gyrus** | BDP vs HC | -.032 | .044 | -0.74 | .461 | .990 |
| **Left Fusiform Gyrus** | BDP vs HC | -.021 | .031 | -0.69 | .493 | .990 |
| **Right Postcentral Gyrus** | BDP vs HC | -.021 | .031 | -0.68 | .500 | .990 |
| **Right Entorhinal Cortex** | BDP vs HC | -.044 | .066 | -0.67 | .506 | .990 |
| **Right Cuneus** | BDP vs HC | -.020 | .031 | -0.65 | .518 | .990 |
| **Left Posterior Cingulate Cortex** | BDP vs HC | -.018 | .029 | -0.63 | .532 | .990 |
| **Right Isthmus of the Cingulate Cortex** | BDP vs HC | -.019 | .030 | -0.62 | .533 | .990 |
| **Left Lateral Orbitofrontal Cortex** | BDP vs HC | -.015 | .026 | -0.60 | .548 | .990 |
| **Left Rostral Anterior Cingulate Cortex** | BDP vs HC | .025 | .042 | 0.59 | .555 | .990 |
| **Left Pars Opercularis** | BDP vs HC | -.015 | .028 | -0.55 | .586 | .990 |
| **Left Caudal Anterior Cingulate Cortex** | BDP vs HC | -.021 | .043 | -0.48 | .630 | .990 |
| **Right Precentral Gyrus** | BDP vs HC | -.018 | .041 | -0.44 | .658 | .990 |
| **Left Pericalcarine Cortex** | BDP vs HC | .014 | .037 | 0.38 | .703 | .990 |
| **Right Posterior Cingulate Cortex** | BDP vs HC | -.011 | .030 | -0.37 | .714 | .990 |
| **Right Caudal Anterior Cingulate Cortex** | BDP vs HC | .013 | .037 | 0.34 | .733 | .990 |
| **Left Caudal Middle Frontal Gyrus** | BDP vs HC | -.009 | .029 | -0.32 | .748 | .990 |
| **Left Postcentral Gyrus** | BDP vs HC | -.009 | .028 | -0.32 | .753 | .990 |
| **Right Transverse Temporal Gyrus** | BDP vs HC | -.014 | .046 | -0.31 | .757 | .990 |
| **Right Pericalcarine Cortex** | BDP vs HC | .011 | .035 | 0.31 | .760 | .990 |
| **Right Insular Cortex** | BDP vs HC | -.009 | .031 | -0.30 | .764 | .990 |
| **Right Superior Temporal Gyrus** | BDP vs HC | .011 | .036 | 0.30 | .765 | .990 |
| **Left Supramarginal Gyrus** | BDP vs HC | .009 | .029 | 0.29 | .772 | .990 |
| **Left Insular Cortex** | BDP vs HC | .008 | .029 | 0.29 | .774 | .990 |
| **Left Inferior Temporal Gyrus** | BDP vs HC | .007 | .030 | 0.23 | .821 | .990 |
| **Right Superior Frontal Gyrus** | BDP vs HC | -.006 | .027 | -0.23 | .821 | .990 |
| **Right Pars Orbitalis** | BDP vs HC | -.009 | .039 | -0.22 | .823 | .990 |
| **Right Caudal Middle Frontal Gyrus** | BDP vs HC | -.006 | .029 | -0.21 | .835 | .990 |
| **Right Banks of the Superior Temporal Sulcus** | BDP vs HC | -.007 | .039 | -0.19 | .848 | .990 |
| **Right Parahippocampal Gyrus** | BDP vs HC | .008 | .046 | 0.17 | .862 | .990 |
| **Left Banks of the Superior Temporal Sulcus** | BDP vs HC | .006 | .036 | 0.15 | .877 | .990 |
| **Left Inferior Parietal Lobule** | BDP vs HC | .004 | .026 | 0.15 | .879 | .990 |
| **Left Rostral Middle Frontal Gyrus** | BDP vs HC | -.003 | .024 | -0.12 | .906 | .990 |
| **Right Supramarginal Gyrus** | BDP vs HC | .004 | .030 | 0.12 | .906 | .990 |
| **Right Inferior Parietal Lobule** | BDP vs HC | .002 | .026 | 0.06 | .948 | .990 |
| **Left Superior Temporal Gyrus** | BDP vs HC | .002 | .033 | 0.06 | .951 | .990 |
| **Left Precuneus** | BDP vs HC | .001 | .024 | 0.05 | .963 | .990 |
| **Left Precentral Gyrus** | BDP vs HC | -.001 | .035 | -0.03 | .976 | .990 |
| **Right Middle Temporal Gyrus** | BDP vs HC | .001 | .029 | 0.03 | .978 | .990 |
| **Left Paracentral Lobule** | BDP vs HC | -.001 | .032 | -0.02 | .982 | .990 |
| **Left Parahippocampal Gyrus** | BDP vs HC | .001 | .063 | 0.01 | .990 | .990 |

**Appendix Table A2.**
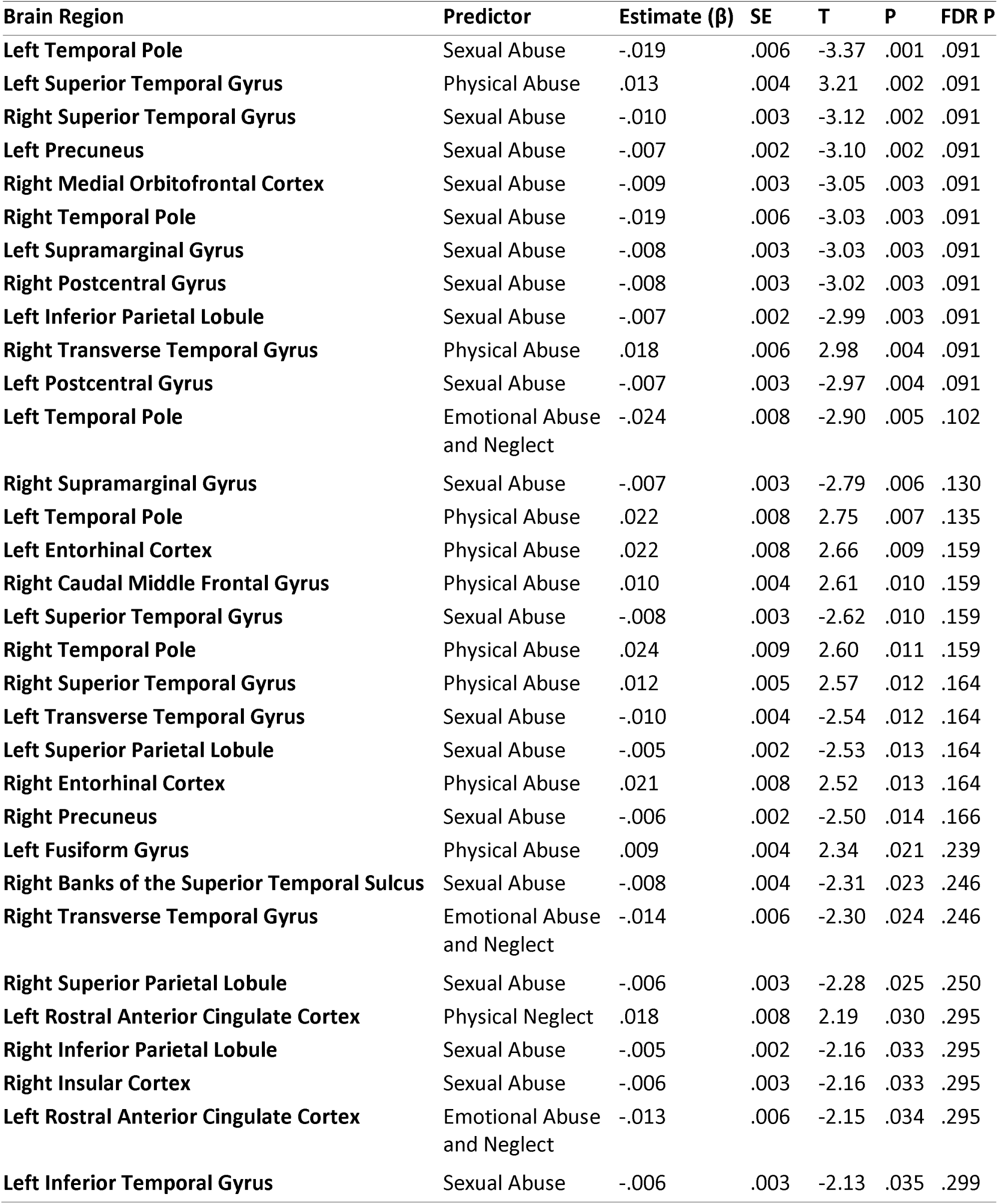
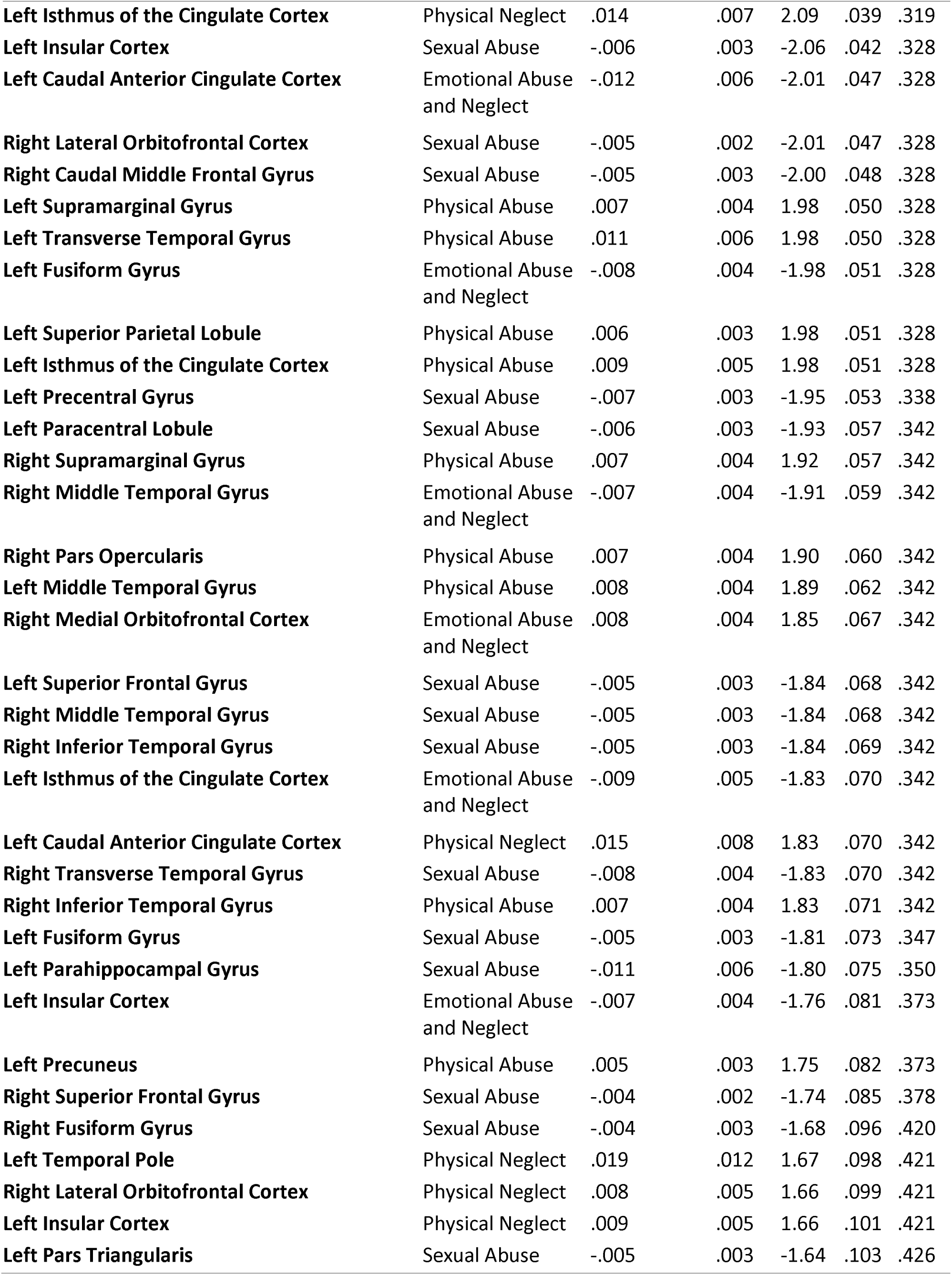
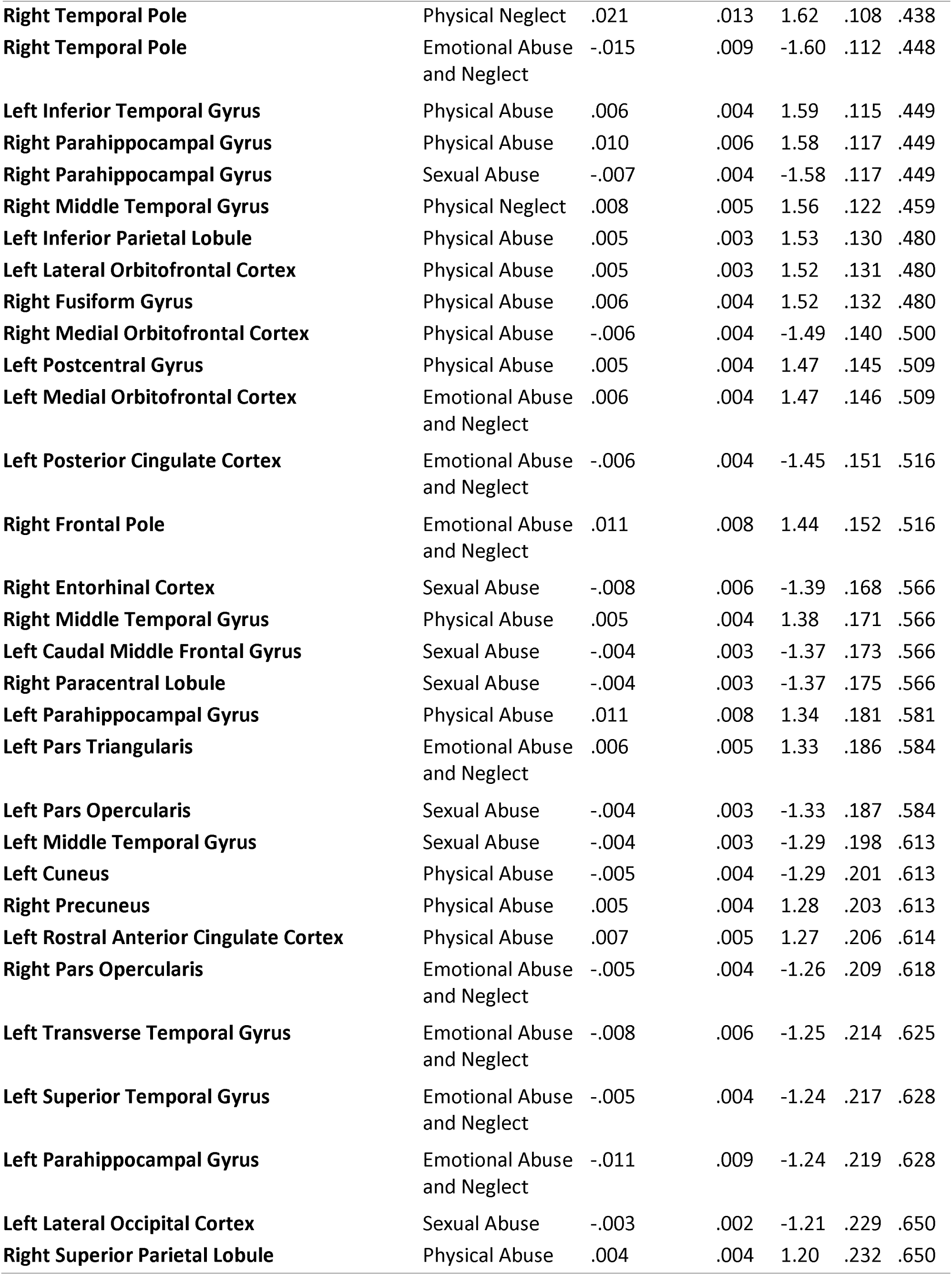
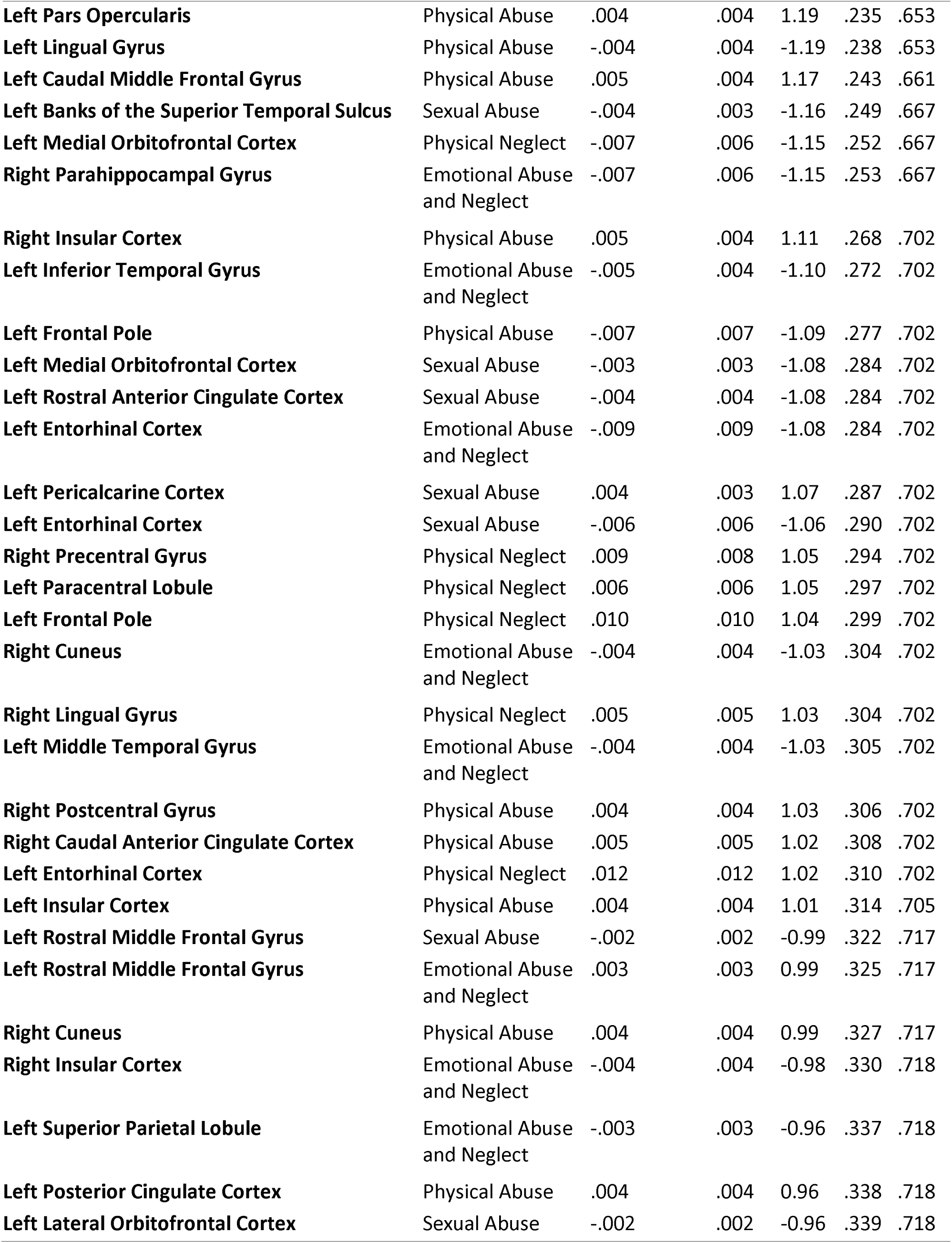
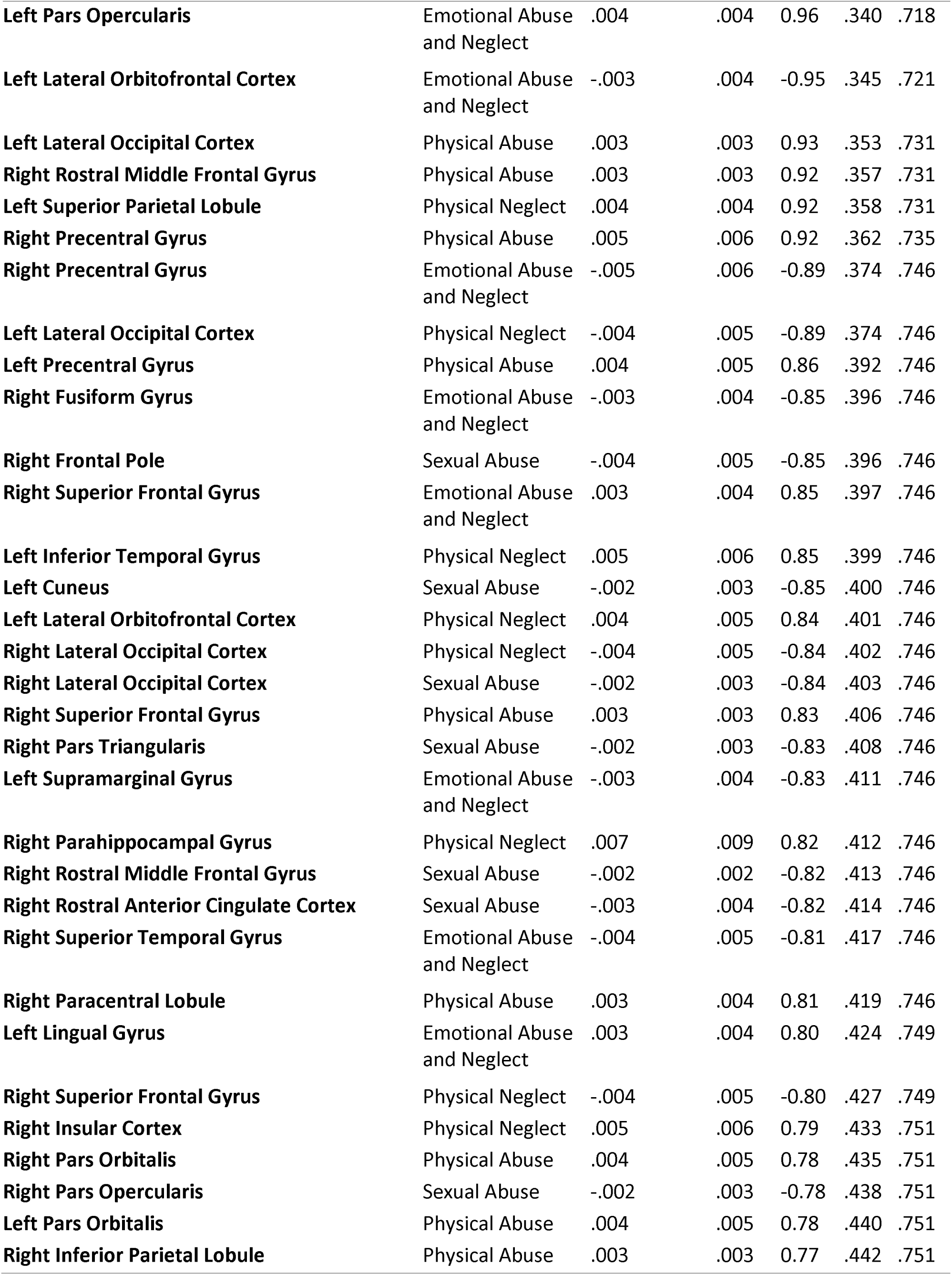
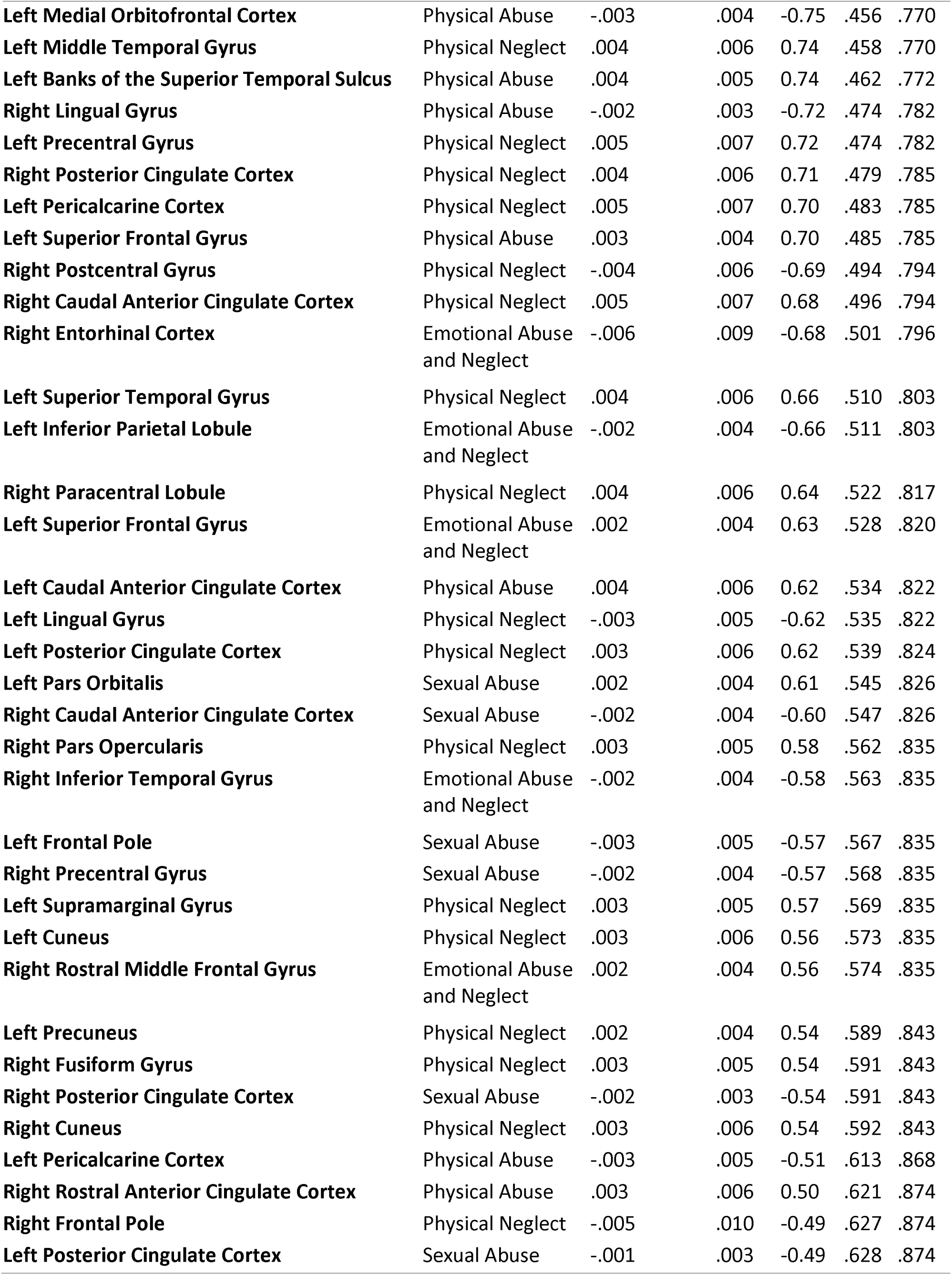
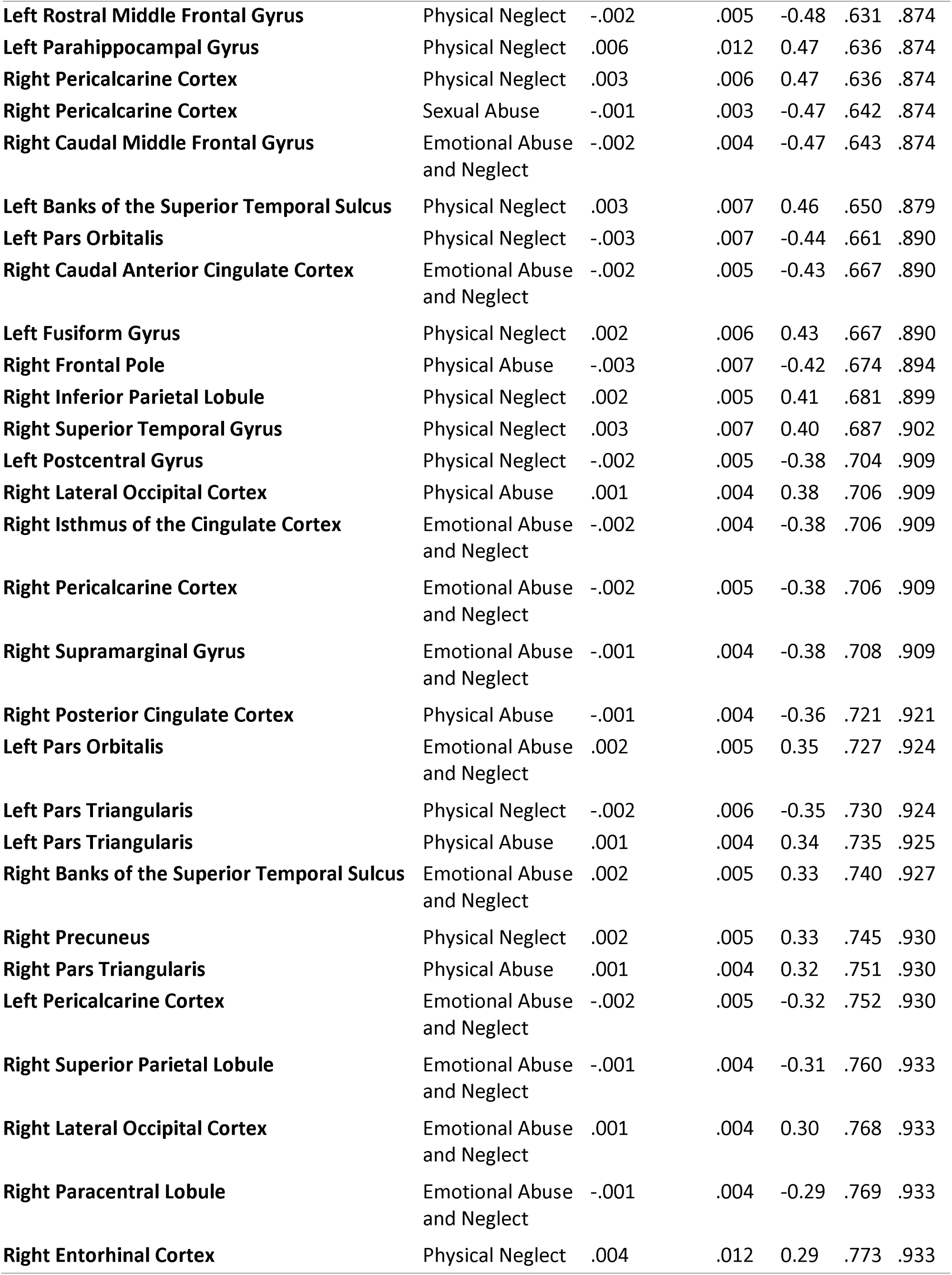
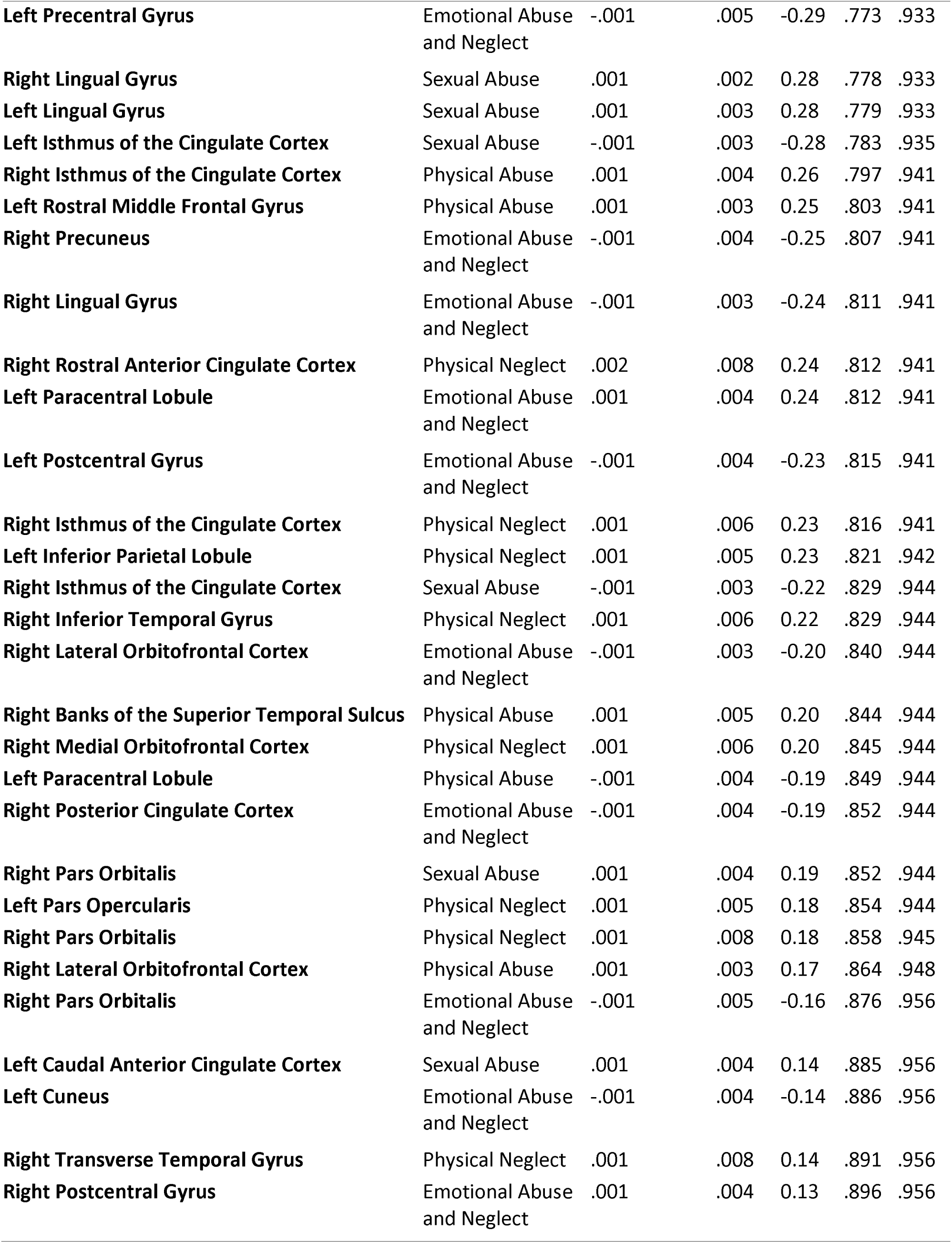
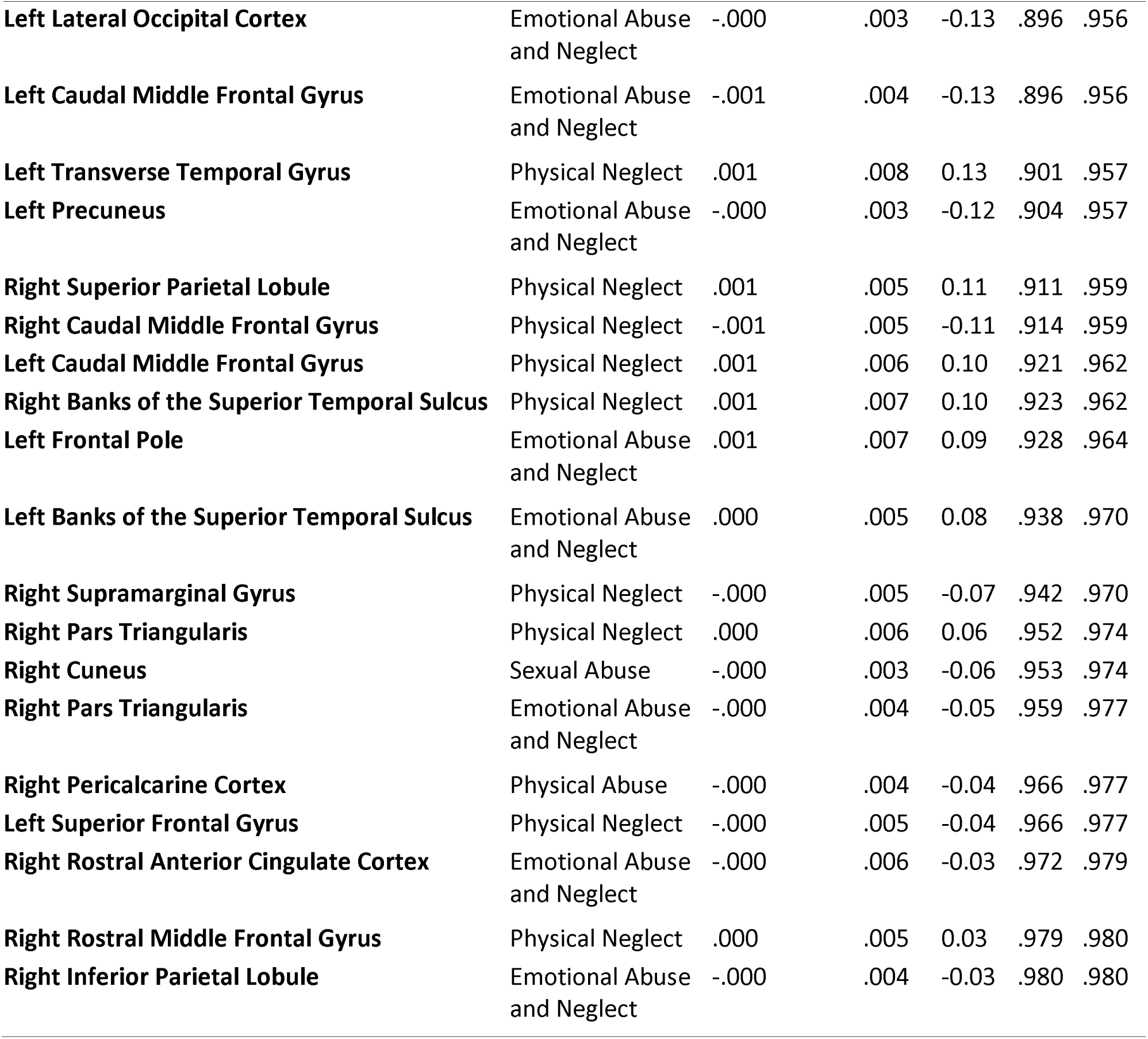
Childhood Trauma Subtype and Cortical Thickness Across 68 Brain.

| Brain Region | Predictor | Estimate ($\beta$) | SE | T | P | FDR P |
| --- | --- | --- | --- | --- | --- | --- |
| Left Temporal Pole | Sexual Abuse | -.019 | .006 | -3.37 | .001 | .091 |
| Left Superior Temporal Gyrus | Physical Abuse | .013 | .004 | 3.21 | .002 | .091 |
| Right Superior Temporal Gyrus | Sexual Abuse | -.010 | .003 | -3.12 | .002 | .091 |
| Left Precuneus | Sexual Abuse | -.007 | .002 | -3.10 | .002 | .091 |
| Right Medial Orbitofrontal Cortex | Sexual Abuse | -.009 | .003 | -3.05 | .003 | .091 |
| Right Temporal Pole | Sexual Abuse | -.019 | .006 | -3.03 | .003 | .091 |
| Left Supramarginal Gyrus | Sexual Abuse | -.008 | .003 | -3.03 | .003 | .091 |
| Right Postcentral Gyrus | Sexual Abuse | -.008 | .003 | -3.02 | .003 | .091 |
| Left Inferior Parietal Lobule | Sexual Abuse | -.007 | .002 | -2.99 | .003 | .091 |
| Right Transverse Temporal Gyrus | Physical Abuse | .018 | .006 | 2.98 | .004 | .091 |
| Left Postcentral Gyrus | Sexual Abuse | -.007 | .003 | -2.97 | .004 | .091 |
| Left Temporal Pole | Emotional Abuse and Neglect | -.024 | .008 | -2.90 | .005 | .102 |
| Right Supramarginal Gyrus | Sexual Abuse | -.007 | .003 | -2.79 | .006 | .130 |
| Left Temporal Pole | Physical Abuse | .022 | .008 | 2.75 | .007 | .135 |
| Left Entorhinal Cortex | Physical Abuse | .022 | .008 | 2.66 | .009 | .159 |
| Right Caudal Middle Frontal Gyrus | Physical Abuse | .010 | .004 | 2.61 | .010 | .159 |
| Left Superior Temporal Gyrus | Sexual Abuse | -.008 | .003 | -2.62 | .010 | .159 |
| Right Temporal Pole | Physical Abuse | .024 | .009 | 2.60 | .011 | .159 |
| Right Superior Temporal Gyrus | Physical Abuse | .012 | .005 | 2.57 | .012 | .164 |
| Left Transverse Temporal Gyrus | Sexual Abuse | -.010 | .004 | -2.54 | .012 | .164 |
| Left Superior Parietal Lobule | Sexual Abuse | -.005 | .002 | -2.53 | .013 | .164 |
| Right Entorhinal Cortex | Physical Abuse | .021 | .008 | 2.52 | .013 | .164 |
| Right Precuneus | Sexual Abuse | -.006 | .002 | -2.50 | .014 | .166 |
| Left Fusiform Gyrus | Physical Abuse | .009 | .004 | 2.34 | .021 | .239 |
| Right Banks of the Superior Temporal Sulcus | Sexual Abuse | -.008 | .004 | -2.31 | .023 | .246 |
| Right Transverse Temporal Gyrus | Emotional Abuse and Neglect | -.014 | .006 | -2.30 | .024 | .246 |
| Right Superior Parietal Lobule | Sexual Abuse | -.006 | .003 | -2.28 | .025 | .250 |
| Left Rostral Anterior Cingulate Cortex | Physical Neglect | .018 | .008 | 2.19 | .030 | .295 |
| Right Inferior Parietal Lobule | Sexual Abuse | -.005 | .002 | -2.16 | .033 | .295 |
| Right Insular Cortex | Sexual Abuse | -.006 | .003 | -2.16 | .033 | .295 |
| Left Rostral Anterior Cingulate Cortex | Emotional Abuse and Neglect | -.013 | .006 | -2.15 | .034 | .295 |
| Left Inferior Temporal Gyrus | Sexual Abuse | -.006 | .003 | -2.13 | .035 | .299 |
| **Left Isthmus of the Cingulate Cortex** | Physical Neglect | .014 | .007 | 2.09 | .039 | .319 |
| **Left Insular Cortex** | Sexual Abuse | -.006 | .003 | -2.06 | .042 | .328 |
| **Left Caudal Anterior Cingulate Cortex** | Emotional Abuse and Neglect | -.012 | .006 | -2.01 | .047 | .328 |
| **Right Lateral Orbitofrontal Cortex** | Sexual Abuse | -.005 | .002 | -2.01 | .047 | .328 |
| **Right Caudal Middle Frontal Gyrus** | Sexual Abuse | -.005 | .003 | -2.00 | .048 | .328 |
| **Left Supramarginal Gyrus** | Physical Abuse | .007 | .004 | 1.98 | .050 | .328 |
| **Left Transverse Temporal Gyrus** | Physical Abuse | .011 | .006 | 1.98 | .050 | .328 |
| **Left Fusiform Gyrus** | Emotional Abuse and Neglect | -.008 | .004 | -1.98 | .051 | .328 |
| **Left Superior Parietal Lobule** | Physical Abuse | .006 | .003 | 1.98 | .051 | .328 |
| **Left Isthmus of the Cingulate Cortex** | Physical Abuse | .009 | .005 | 1.98 | .051 | .328 |
| **Left Precentral Gyrus** | Sexual Abuse | -.007 | .003 | -1.95 | .053 | .338 |
| **Left Paracentral Lobule** | Sexual Abuse | -.006 | .003 | -1.93 | .057 | .342 |
| **Right Supramarginal Gyrus** | Physical Abuse | .007 | .004 | 1.92 | .057 | .342 |
| **Right Middle Temporal Gyrus** | Emotional Abuse and Neglect | -.007 | .004 | -1.91 | .059 | .342 |
| **Right Pars Opercularis** | Physical Abuse | .007 | .004 | 1.90 | .060 | .342 |
| **Left Middle Temporal Gyrus** | Physical Abuse | .008 | .004 | 1.89 | .062 | .342 |
| **Right Medial Orbitofrontal Cortex** | Emotional Abuse and Neglect | .008 | .004 | 1.85 | .067 | .342 |
| **Left Superior Frontal Gyrus** | Sexual Abuse | -.005 | .003 | -1.84 | .068 | .342 |
| **Right Middle Temporal Gyrus** | Sexual Abuse | -.005 | .003 | -1.84 | .068 | .342 |
| **Right Inferior Temporal Gyrus** | Sexual Abuse | -.005 | .003 | -1.84 | .069 | .342 |
| **Left Isthmus of the Cingulate Cortex** | Emotional Abuse and Neglect | -.009 | .005 | -1.83 | .070 | .342 |
| **Left Caudal Anterior Cingulate Cortex** | Physical Neglect | .015 | .008 | 1.83 | .070 | .342 |
| **Right Transverse Temporal Gyrus** | Sexual Abuse | -.008 | .004 | -1.83 | .070 | .342 |
| **Right Inferior Temporal Gyrus** | Physical Abuse | .007 | .004 | 1.83 | .071 | .342 |
| **Left Fusiform Gyrus** | Sexual Abuse | -.005 | .003 | -1.81 | .073 | .347 |
| **Left Parahippocampal Gyrus** | Sexual Abuse | -.011 | .006 | -1.80 | .075 | .350 |
| **Left Insular Cortex** | Emotional Abuse and Neglect | -.007 | .004 | -1.76 | .081 | .373 |
| **Left Precuneus** | Physical Abuse | .005 | .003 | 1.75 | .082 | .373 |
| **Right Superior Frontal Gyrus** | Sexual Abuse | -.004 | .002 | -1.74 | .085 | .378 |
| **Right Fusiform Gyrus** | Sexual Abuse | -.004 | .003 | -1.68 | .096 | .420 |
| **Left Temporal Pole** | Physical Neglect | .019 | .012 | 1.67 | .098 | .421 |
| **Right Lateral Orbitofrontal Cortex** | Physical Neglect | .008 | .005 | 1.66 | .099 | .421 |
| **Left Insular Cortex** | Physical Neglect | .009 | .005 | 1.66 | .101 | .421 |
| **Left Pars Triangularis** | Sexual Abuse | -.005 | .003 | -1.64 | .103 | .426 |
| **Right Temporal Pole** | Physical Neglect | .021 | .013 | 1.62 | .108 | .438 |
| **Right Temporal Pole** | Emotional Abuse and Neglect | -.015 | .009 | -1.60 | .112 | .448 |
| **Left Inferior Temporal Gyrus** | Physical Abuse | .006 | .004 | 1.59 | .115 | .449 |
| **Right Parahippocampal Gyrus** | Physical Abuse | .010 | .006 | 1.58 | .117 | .449 |
| **Right Parahippocampal Gyrus** | Sexual Abuse | -.007 | .004 | -1.58 | .117 | .449 |
| **Right Middle Temporal Gyrus** | Physical Neglect | .008 | .005 | 1.56 | .122 | .459 |
| **Left Inferior Parietal Lobule** | Physical Abuse | .005 | .003 | 1.53 | .130 | .480 |
| **Left Lateral Orbitofrontal Cortex** | Physical Abuse | .005 | .003 | 1.52 | .131 | .480 |
| **Right Fusiform Gyrus** | Physical Abuse | .006 | .004 | 1.52 | .132 | .480 |
| **Right Medial Orbitofrontal Cortex** | Physical Abuse | -.006 | .004 | -1.49 | .140 | .500 |
| **Left Postcentral Gyrus** | Physical Abuse | .005 | .004 | 1.47 | .145 | .509 |
| **Left Medial Orbitofrontal Cortex** | Emotional Abuse and Neglect | .006 | .004 | 1.47 | .146 | .509 |
| **Left Posterior Cingulate Cortex** | Emotional Abuse and Neglect | -.006 | .004 | -1.45 | .151 | .516 |
| **Right Frontal Pole** | Emotional Abuse and Neglect | .011 | .008 | 1.44 | .152 | .516 |
| **Right Entorhinal Cortex** | Sexual Abuse | -.008 | .006 | -1.39 | .168 | .566 |
| **Right Middle Temporal Gyrus** | Physical Abuse | .005 | .004 | 1.38 | .171 | .566 |
| **Left Caudal Middle Frontal Gyrus** | Sexual Abuse | -.004 | .003 | -1.37 | .173 | .566 |
| **Right Paracentral Lobule** | Sexual Abuse | -.004 | .003 | -1.37 | .175 | .566 |
| **Left Parahippocampal Gyrus** | Physical Abuse | .011 | .008 | 1.34 | .181 | .581 |
| **Left Pars Triangularis** | Emotional Abuse and Neglect | .006 | .005 | 1.33 | .186 | .584 |
| **Left Pars Opercularis** | Sexual Abuse | -.004 | .003 | -1.33 | .187 | .584 |
| **Left Middle Temporal Gyrus** | Sexual Abuse | -.004 | .003 | -1.29 | .198 | .613 |
| **Left Cuneus** | Physical Abuse | -.005 | .004 | -1.29 | .201 | .613 |
| **Right Precuneus** | Physical Abuse | .005 | .004 | 1.28 | .203 | .613 |
| **Left Rostral Anterior Cingulate Cortex** | Physical Abuse | .007 | .005 | 1.27 | .206 | .614 |
| **Right Pars Opercularis** | Emotional Abuse and Neglect | -.005 | .004 | -1.26 | .209 | .618 |
| **Left Transverse Temporal Gyrus** | Emotional Abuse and Neglect | -.008 | .006 | -1.25 | .214 | .625 |
| **Left Superior Temporal Gyrus** | Emotional Abuse and Neglect | -.005 | .004 | -1.24 | .217 | .628 |
| **Left Parahippocampal Gyrus** | Emotional Abuse and Neglect | -.011 | .009 | -1.24 | .219 | .628 |
| **Left Lateral Occipital Cortex** | Sexual Abuse | -.003 | .002 | -1.21 | .229 | .650 |
| **Right Superior Parietal Lobule** | Physical Abuse | .004 | .004 | 1.20 | .232 | .650 |
| **Left Pars Opercularis** | Physical Abuse | .004 | .004 | 1.19 | .235 | .653 |
| **Left Lingual Gyrus** | Physical Abuse | -.004 | .004 | -1.19 | .238 | .653 |
| **Left Caudal Middle Frontal Gyrus** | Physical Abuse | .005 | .004 | 1.17 | .243 | .661 |
| **Left Banks of the Superior Temporal Sulcus** | Sexual Abuse | -.004 | .003 | -1.16 | .249 | .667 |
| **Left Medial Orbitofrontal Cortex** | Physical Neglect | -.007 | .006 | -1.15 | .252 | .667 |
| **Right Parahippocampal Gyrus** | Emotional Abuse and Neglect | -.007 | .006 | -1.15 | .253 | .667 |
| **Right Insular Cortex** | Physical Abuse | .005 | .004 | 1.11 | .268 | .702 |
| **Left Inferior Temporal Gyrus** | Emotional Abuse and Neglect | -.005 | .004 | -1.10 | .272 | .702 |
| **Left Frontal Pole** | Physical Abuse | -.007 | .007 | -1.09 | .277 | .702 |
| **Left Medial Orbitofrontal Cortex** | Sexual Abuse | -.003 | .003 | -1.08 | .284 | .702 |
| **Left Rostral Anterior Cingulate Cortex** | Sexual Abuse | -.004 | .004 | -1.08 | .284 | .702 |
| **Left Entorhinal Cortex** | Emotional Abuse and Neglect | -.009 | .009 | -1.08 | .284 | .702 |
| **Left Pericalcarine Cortex** | Sexual Abuse | .004 | .003 | 1.07 | .287 | .702 |
| **Left Entorhinal Cortex** | Sexual Abuse | -.006 | .006 | -1.06 | .290 | .702 |
| **Right Precentral Gyrus** | Physical Neglect | .009 | .008 | 1.05 | .294 | .702 |
| **Left Paracentral Lobule** | Physical Neglect | .006 | .006 | 1.05 | .297 | .702 |
| **Left Frontal Pole** | Physical Neglect | .010 | .010 | 1.04 | .299 | .702 |
| **Right Cuneus** | Emotional Abuse and Neglect | -.004 | .004 | -1.03 | .304 | .702 |
| **Right Lingual Gyrus** | Physical Neglect | .005 | .005 | 1.03 | .304 | .702 |
| **Left Middle Temporal Gyrus** | Emotional Abuse and Neglect | -.004 | .004 | -1.03 | .305 | .702 |
| **Right Postcentral Gyrus** | Physical Abuse | .004 | .004 | 1.03 | .306 | .702 |
| **Right Caudal Anterior Cingulate Cortex** | Physical Abuse | .005 | .005 | 1.02 | .308 | .702 |
| **Left Entorhinal Cortex** | Physical Neglect | .012 | .012 | 1.02 | .310 | .702 |
| **Left Insular Cortex** | Physical Abuse | .004 | .004 | 1.01 | .314 | .705 |
| **Left Rostral Middle Frontal Gyrus** | Sexual Abuse | -.002 | .002 | -0.99 | .322 | .717 |
| **Left Rostral Middle Frontal Gyrus** | Emotional Abuse and Neglect | .003 | .003 | 0.99 | .325 | .717 |
| **Right Cuneus** | Physical Abuse | .004 | .004 | 0.99 | .327 | .717 |
| **Right Insular Cortex** | Emotional Abuse and Neglect | -.004 | .004 | -0.98 | .330 | .718 |
| **Left Superior Parietal Lobule** | Emotional Abuse and Neglect | -.003 | .003 | -0.96 | .337 | .718 |
| **Left Posterior Cingulate Cortex** | Physical Abuse | .004 | .004 | 0.96 | .338 | .718 |
| **Left Lateral Orbitofrontal Cortex** | Sexual Abuse | -.002 | .002 | -0.96 | .339 | .718 |
| **Left Pars Opercularis** | Emotional Abuse and Neglect | .004 | .004 | 0.96 | .340 | .718 |
| **Left Lateral Orbitofrontal Cortex** | Emotional Abuse and Neglect | -.003 | .004 | -0.95 | .345 | .721 |
| **Left Lateral Occipital Cortex** | Physical Abuse | .003 | .003 | 0.93 | .353 | .731 |
| **Right Rostral Middle Frontal Gyrus** | Physical Abuse | .003 | .003 | 0.92 | .357 | .731 |
| **Left Superior Parietal Lobule** | Physical Neglect | .004 | .004 | 0.92 | .358 | .731 |
| **Right Precentral Gyrus** | Physical Abuse | .005 | .006 | 0.92 | .362 | .735 |
| **Right Precentral Gyrus** | Emotional Abuse and Neglect | -.005 | .006 | -0.89 | .374 | .746 |
| **Left Lateral Occipital Cortex** | Physical Neglect | -.004 | .005 | -0.89 | .374 | .746 |
| **Left Precentral Gyrus** | Physical Abuse | .004 | .005 | 0.86 | .392 | .746 |
| **Right Fusiform Gyrus** | Emotional Abuse and Neglect | -.003 | .004 | -0.85 | .396 | .746 |
| **Right Frontal Pole** | Sexual Abuse | -.004 | .005 | -0.85 | .396 | .746 |
| **Right Superior Frontal Gyrus** | Emotional Abuse and Neglect | .003 | .004 | 0.85 | .397 | .746 |
| **Left Inferior Temporal Gyrus** | Physical Neglect | .005 | .006 | 0.85 | .399 | .746 |
| **Left Cuneus** | Sexual Abuse | -.002 | .003 | -0.85 | .400 | .746 |
| **Left Lateral Orbitofrontal Cortex** | Physical Neglect | .004 | .005 | 0.84 | .401 | .746 |
| **Right Lateral Occipital Cortex** | Physical Neglect | -.004 | .005 | -0.84 | .402 | .746 |
| **Right Lateral Occipital Cortex** | Sexual Abuse | -.002 | .003 | -0.84 | .403 | .746 |
| **Right Superior Frontal Gyrus** | Physical Abuse | .003 | .003 | 0.83 | .406 | .746 |
| **Right Pars Triangularis** | Sexual Abuse | -.002 | .003 | -0.83 | .408 | .746 |
| **Left Supramarginal Gyrus** | Emotional Abuse and Neglect | -.003 | .004 | -0.83 | .411 | .746 |
| **Right Parahippocampal Gyrus** | Physical Neglect | .007 | .009 | 0.82 | .412 | .746 |
| **Right Rostral Middle Frontal Gyrus** | Sexual Abuse | -.002 | .002 | -0.82 | .413 | .746 |
| **Right Rostral Anterior Cingulate Cortex** | Sexual Abuse | -.003 | .004 | -0.82 | .414 | .746 |
| **Right Superior Temporal Gyrus** | Emotional Abuse and Neglect | -.004 | .005 | -0.81 | .417 | .746 |
| **Right Paracentral Lobule** | Physical Abuse | .003 | .004 | 0.81 | .419 | .746 |
| **Left Lingual Gyrus** | Emotional Abuse and Neglect | .003 | .004 | 0.80 | .424 | .749 |
| **Right Superior Frontal Gyrus** | Physical Neglect | -.004 | .005 | -0.80 | .427 | .749 |
| **Right Insular Cortex** | Physical Neglect | .005 | .006 | 0.79 | .433 | .751 |
| **Right Pars Orbitalis** | Physical Abuse | .004 | .005 | 0.78 | .435 | .751 |
| **Right Pars Opercularis** | Sexual Abuse | -.002 | .003 | -0.78 | .438 | .751 |
| **Left Pars Orbitalis** | Physical Abuse | .004 | .005 | 0.78 | .440 | .751 |
| **Right Inferior Parietal Lobule** | Physical Abuse | .003 | .003 | 0.77 | .442 | .751 |
| **Left Medial Orbitofrontal Cortex** | Physical Abuse | -.003 | .004 | -0.75 | .456 | .770 |
| **Left Middle Temporal Gyrus** | Physical Neglect | .004 | .006 | 0.74 | .458 | .770 |
| **Left Banks of the Superior Temporal Sulcus** | Physical Abuse | .004 | .005 | 0.74 | .462 | .772 |
| **Right Lingual Gyrus** | Physical Abuse | -.002 | .003 | -0.72 | .474 | .782 |
| **Left Precentral Gyrus** | Physical Neglect | .005 | .007 | 0.72 | .474 | .782 |
| **Right Posterior Cingulate Cortex** | Physical Neglect | .004 | .006 | 0.71 | .479 | .785 |
| **Left Pericalcarine Cortex** | Physical Neglect | .005 | .007 | 0.70 | .483 | .785 |
| **Left Superior Frontal Gyrus** | Physical Abuse | .003 | .004 | 0.70 | .485 | .785 |
| **Right Postcentral Gyrus** | Physical Neglect | -.004 | .006 | -0.69 | .494 | .794 |
| **Right Caudal Anterior Cingulate Cortex** | Physical Neglect | .005 | .007 | 0.68 | .496 | .794 |
| **Right Entorhinal Cortex** | Emotional Abuse and Neglect | -.006 | .009 | -0.68 | .501 | .796 |
| **Left Superior Temporal Gyrus** | Physical Neglect | .004 | .006 | 0.66 | .510 | .803 |
| **Left Inferior Parietal Lobule** | Emotional Abuse and Neglect | -.002 | .004 | -0.66 | .511 | .803 |
| **Right Paracentral Lobule** | Physical Neglect | .004 | .006 | 0.64 | .522 | .817 |
| **Left Superior Frontal Gyrus** | Emotional Abuse and Neglect | .002 | .004 | 0.63 | .528 | .820 |
| **Left Caudal Anterior Cingulate Cortex** | Physical Abuse | .004 | .006 | 0.62 | .534 | .822 |
| **Left Lingual Gyrus** | Physical Neglect | -.003 | .005 | -0.62 | .535 | .822 |
| **Left Posterior Cingulate Cortex** | Physical Neglect | .003 | .006 | 0.62 | .539 | .824 |
| **Left Pars Orbitalis** | Sexual Abuse | .002 | .004 | 0.61 | .545 | .826 |
| **Right Caudal Anterior Cingulate Cortex** | Sexual Abuse | -.002 | .004 | -0.60 | .547 | .826 |
| **Right Pars Opercularis** | Physical Neglect | .003 | .005 | 0.58 | .562 | .835 |
| **Right Inferior Temporal Gyrus** | Emotional Abuse and Neglect | -.002 | .004 | -0.58 | .563 | .835 |
| **Left Frontal Pole** | Sexual Abuse | -.003 | .005 | -0.57 | .567 | .835 |
| **Right Precentral Gyrus** | Sexual Abuse | -.002 | .004 | -0.57 | .568 | .835 |
| **Left Supramarginal Gyrus** | Physical Neglect | .003 | .005 | 0.57 | .569 | .835 |
| **Left Cuneus** | Physical Neglect | .003 | .006 | 0.56 | .573 | .835 |
| **Right Rostral Middle Frontal Gyrus** | Emotional Abuse and Neglect | .002 | .004 | 0.56 | .574 | .835 |
| **Left Precuneus** | Physical Neglect | .002 | .004 | 0.54 | .589 | .843 |
| **Right Fusiform Gyrus** | Physical Neglect | .003 | .005 | 0.54 | .591 | .843 |
| **Right Posterior Cingulate Cortex** | Sexual Abuse | -.002 | .003 | -0.54 | .591 | .843 |
| **Right Cuneus** | Physical Neglect | .003 | .006 | 0.54 | .592 | .843 |
| **Left Pericalcarine Cortex** | Physical Abuse | -.003 | .005 | -0.51 | .613 | .868 |
| **Right Rostral Anterior Cingulate Cortex** | Physical Abuse | .003 | .006 | 0.50 | .621 | .874 |
| **Right Frontal Pole** | Physical Neglect | -.005 | .010 | -0.49 | .627 | .874 |
| **Left Posterior Cingulate Cortex** | Sexual Abuse | -.001 | .003 | -0.49 | .628 | .874 |
| **Left Rostral Middle Frontal Gyrus** | Physical Neglect | -.002 | .005 | -0.48 | .631 | .874 |
| **Left Parahippocampal Gyrus** | Physical Neglect | .006 | .012 | 0.47 | .636 | .874 |
| **Right Pericalcarine Cortex** | Physical Neglect | .003 | .006 | 0.47 | .636 | .874 |
| **Right Pericalcarine Cortex** | Sexual Abuse | -.001 | .003 | -0.47 | .642 | .874 |
| **Right Caudal Middle Frontal Gyrus** | Emotional Abuse and Neglect | -.002 | .004 | -0.47 | .643 | .874 |
| **Left Banks of the Superior Temporal Sulcus** | Physical Neglect | .003 | .007 | 0.46 | .650 | .879 |
| **Left Pars Orbitalis** | Physical Neglect | -.003 | .007 | -0.44 | .661 | .890 |
| **Right Caudal Anterior Cingulate Cortex** | Emotional Abuse and Neglect | -.002 | .005 | -0.43 | .667 | .890 |
| **Left Fusiform Gyrus** | Physical Neglect | .002 | .006 | 0.43 | .667 | .890 |
| **Right Frontal Pole** | Physical Abuse | -.003 | .007 | -0.42 | .674 | .894 |
| **Right Inferior Parietal Lobule** | Physical Neglect | .002 | .005 | 0.41 | .681 | .899 |
| **Right Superior Temporal Gyrus** | Physical Neglect | .003 | .007 | 0.40 | .687 | .902 |
| **Left Postcentral Gyrus** | Physical Neglect | -.002 | .005 | -0.38 | .704 | .909 |
| **Right Lateral Occipital Cortex** | Physical Abuse | .001 | .004 | 0.38 | .706 | .909 |
| **Right Isthmus of the Cingulate Cortex** | Emotional Abuse and Neglect | -.002 | .004 | -0.38 | .706 | .909 |
| **Right Pericalcarine Cortex** | Emotional Abuse and Neglect | -.002 | .005 | -0.38 | .706 | .909 |
| **Right Supramarginal Gyrus** | Emotional Abuse and Neglect | -.001 | .004 | -0.38 | .708 | .909 |
| **Right Posterior Cingulate Cortex** | Physical Abuse | -.001 | .004 | -0.36 | .721 | .921 |
| **Left Pars Orbitalis** | Emotional Abuse and Neglect | .002 | .005 | 0.35 | .727 | .924 |
| **Left Pars Triangularis** | Physical Neglect | -.002 | .006 | -0.35 | .730 | .924 |
| **Left Pars Triangularis** | Physical Abuse | .001 | .004 | 0.34 | .735 | .925 |
| **Right Banks of the Superior Temporal Sulcus** | Emotional Abuse and Neglect | .002 | .005 | 0.33 | .740 | .927 |
| **Right Precuneus** | Physical Neglect | .002 | .005 | 0.33 | .745 | .930 |
| **Right Pars Triangularis** | Physical Abuse | .001 | .004 | 0.32 | .751 | .930 |
| **Left Pericalcarine Cortex** | Emotional Abuse and Neglect | -.002 | .005 | -0.32 | .752 | .930 |
| **Right Superior Parietal Lobule** | Emotional Abuse and Neglect | -.001 | .004 | -0.31 | .760 | .933 |
| **Right Lateral Occipital Cortex** | Emotional Abuse and Neglect | .001 | .004 | 0.30 | .768 | .933 |
| **Right Paracentral Lobule** | Emotional Abuse and Neglect | -.001 | .004 | -0.29 | .769 | .933 |
| **Right Entorhinal Cortex** | Physical Neglect | .004 | .012 | 0.29 | .773 | .933 |
| **Left Precentral Gyrus** | Emotional Abuse and Neglect | -.001 | .005 | -0.29 | .773 | .933 |
| **Right Lingual Gyrus** | Sexual Abuse | .001 | .002 | 0.28 | .778 | .933 |
| **Left Lingual Gyrus** | Sexual Abuse | .001 | .003 | 0.28 | .779 | .933 |
| **Left Isthmus of the Cingulate Cortex** | Sexual Abuse | -.001 | .003 | -0.28 | .783 | .935 |
| **Right Isthmus of the Cingulate Cortex** | Physical Abuse | .001 | .004 | 0.26 | .797 | .941 |
| **Left Rostral Middle Frontal Gyrus** | Physical Abuse | .001 | .003 | 0.25 | .803 | .941 |
| **Right Precuneus** | Emotional Abuse and Neglect | -.001 | .004 | -0.25 | .807 | .941 |
| **Right Lingual Gyrus** | Emotional Abuse and Neglect | -.001 | .003 | -0.24 | .811 | .941 |
| **Right Rostral Anterior Cingulate Cortex** | Physical Neglect | .002 | .008 | 0.24 | .812 | .941 |
| **Left Paracentral Lobule** | Emotional Abuse and Neglect | .001 | .004 | 0.24 | .812 | .941 |
| **Left Postcentral Gyrus** | Emotional Abuse and Neglect | -.001 | .004 | -0.23 | .815 | .941 |
| **Right Isthmus of the Cingulate Cortex** | Physical Neglect | .001 | .006 | 0.23 | .816 | .941 |
| **Left Inferior Parietal Lobule** | Physical Neglect | .001 | .005 | 0.23 | .821 | .942 |
| **Right Isthmus of the Cingulate Cortex** | Sexual Abuse | -.001 | .003 | -0.22 | .829 | .944 |
| **Right Inferior Temporal Gyrus** | Physical Neglect | .001 | .006 | 0.22 | .829 | .944 |
| **Right Lateral Orbitofrontal Cortex** | Emotional Abuse and Neglect | -.001 | .003 | -0.20 | .840 | .944 |
| **Right Banks of the Superior Temporal Sulcus** | Physical Abuse | .001 | .005 | 0.20 | .844 | .944 |
| **Right Medial Orbitofrontal Cortex** | Physical Neglect | .001 | .006 | 0.20 | .845 | .944 |
| **Left Paracentral Lobule** | Physical Abuse | -.001 | .004 | -0.19 | .849 | .944 |
| **Right Posterior Cingulate Cortex** | Emotional Abuse and Neglect | -.001 | .004 | -0.19 | .852 | .944 |
| **Right Pars Orbitalis** | Sexual Abuse | .001 | .004 | 0.19 | .852 | .944 |
| **Left Pars Opercularis** | Physical Neglect | .001 | .005 | 0.18 | .854 | .944 |
| **Right Pars Orbitalis** | Physical Neglect | .001 | .008 | 0.18 | .858 | .945 |
| **Right Lateral Orbitofrontal Cortex** | Physical Abuse | .001 | .003 | 0.17 | .864 | .948 |
| **Right Pars Orbitalis** | Emotional Abuse and Neglect | -.001 | .005 | -0.16 | .876 | .956 |
| **Left Caudal Anterior Cingulate Cortex** | Sexual Abuse | .001 | .004 | 0.14 | .885 | .956 |
| **Left Cuneus** | Emotional Abuse and Neglect | -.001 | .004 | -0.14 | .886 | .956 |
| **Right Transverse Temporal Gyrus** | Physical Neglect | .001 | .008 | 0.14 | .891 | .956 |
| **Right Postcentral Gyrus** | Emotional Abuse and Neglect | .001 | .004 | 0.13 | .896 | .956 |
| **Left Lateral Occipital Cortex** | Emotional Abuse and Neglect | -.000 | .003 | -0.13 | .896 | .956 |
| **Left Caudal Middle Frontal Gyrus** | Emotional Abuse and Neglect | -.001 | .004 | -0.13 | .896 | .956 |
| **Left Transverse Temporal Gyrus** | Physical Neglect | .001 | .008 | 0.13 | .901 | .957 |
| **Left Precuneus** | Emotional Abuse and Neglect | -.000 | .003 | -0.12 | .904 | .957 |
| **Right Superior Parietal Lobule** | Physical Neglect | .001 | .005 | 0.11 | .911 | .959 |
| **Right Caudal Middle Frontal Gyrus** | Physical Neglect | -.001 | .005 | -0.11 | .914 | .959 |
| **Left Caudal Middle Frontal Gyrus** | Physical Neglect | .001 | .006 | 0.10 | .921 | .962 |
| **Right Banks of the Superior Temporal Sulcus** | Physical Neglect | .001 | .007 | 0.10 | .923 | .962 |
| **Left Frontal Pole** | Emotional Abuse and Neglect | .001 | .007 | 0.09 | .928 | .964 |
| **Left Banks of the Superior Temporal Sulcus** | Emotional Abuse and Neglect | .000 | .005 | 0.08 | .938 | .970 |
| **Right Supramarginal Gyrus** | Physical Neglect | -.000 | .005 | -0.07 | .942 | .970 |
| **Right Pars Triangularis** | Physical Neglect | .000 | .006 | 0.06 | .952 | .974 |
| **Right Cuneus** | Sexual Abuse | -.000 | .003 | -0.06 | .953 | .974 |
| **Right Pars Triangularis** | Emotional Abuse and Neglect | -.000 | .004 | -0.05 | .959 | .977 |
| **Right Pericalcarine Cortex** | Physical Abuse | -.000 | .004 | -0.04 | .966 | .977 |
| **Left Superior Frontal Gyrus** | Physical Neglect | -.000 | .005 | -0.04 | .966 | .977 |
| **Right Rostral Anterior Cingulate Cortex** | Emotional Abuse and Neglect | -.000 | .006 | -0.03 | .972 | .979 |
| **Right Rostral Middle Frontal Gyrus** | Physical Neglect | .000 | .005 | 0.03 | .979 | .980 |
| **Right Inferior Parietal Lobule** | Emotional Abuse and Neglect | -.000 | .004 | -0.03 | .980 | .980 |

**Appendix Table A3.**
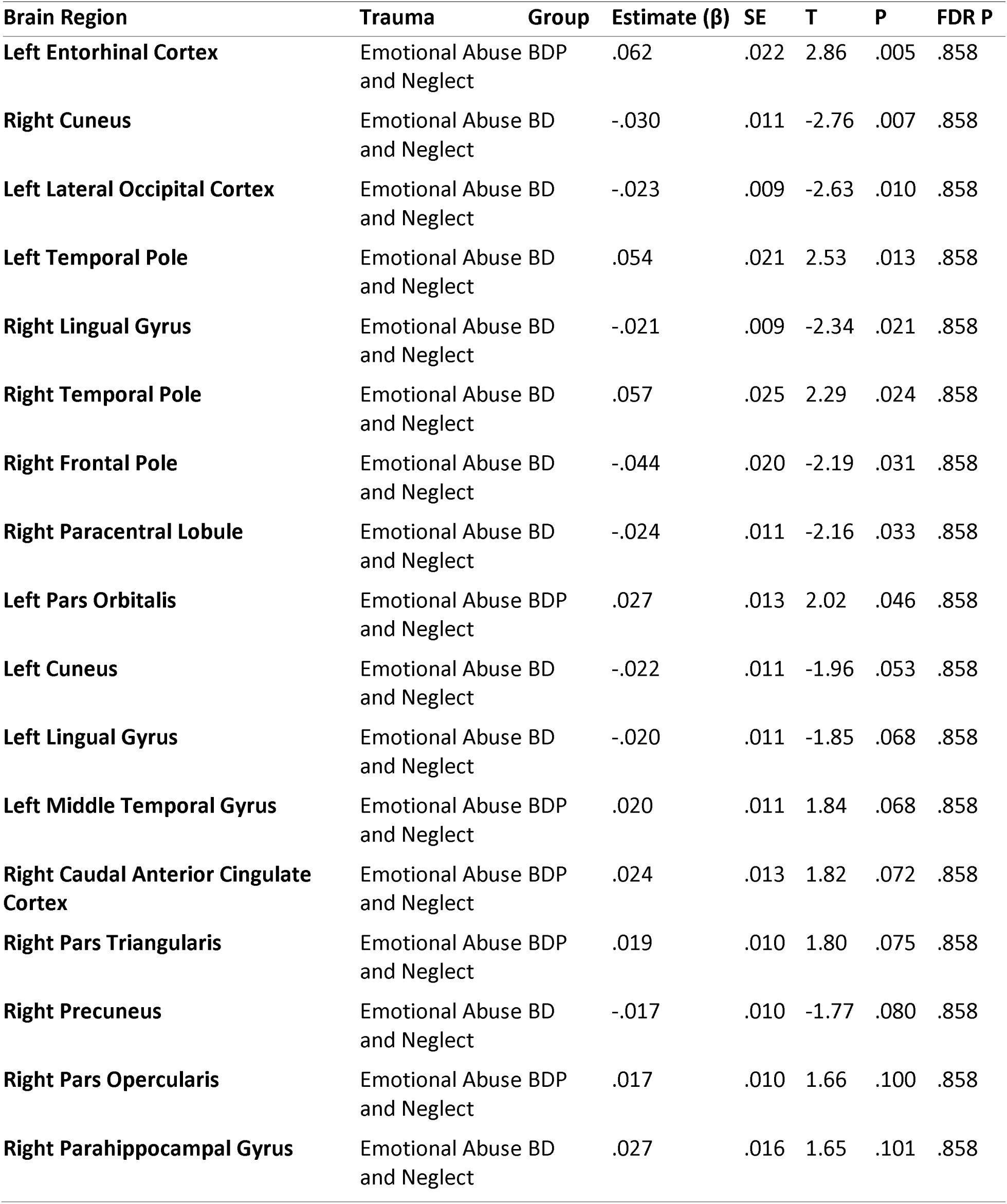
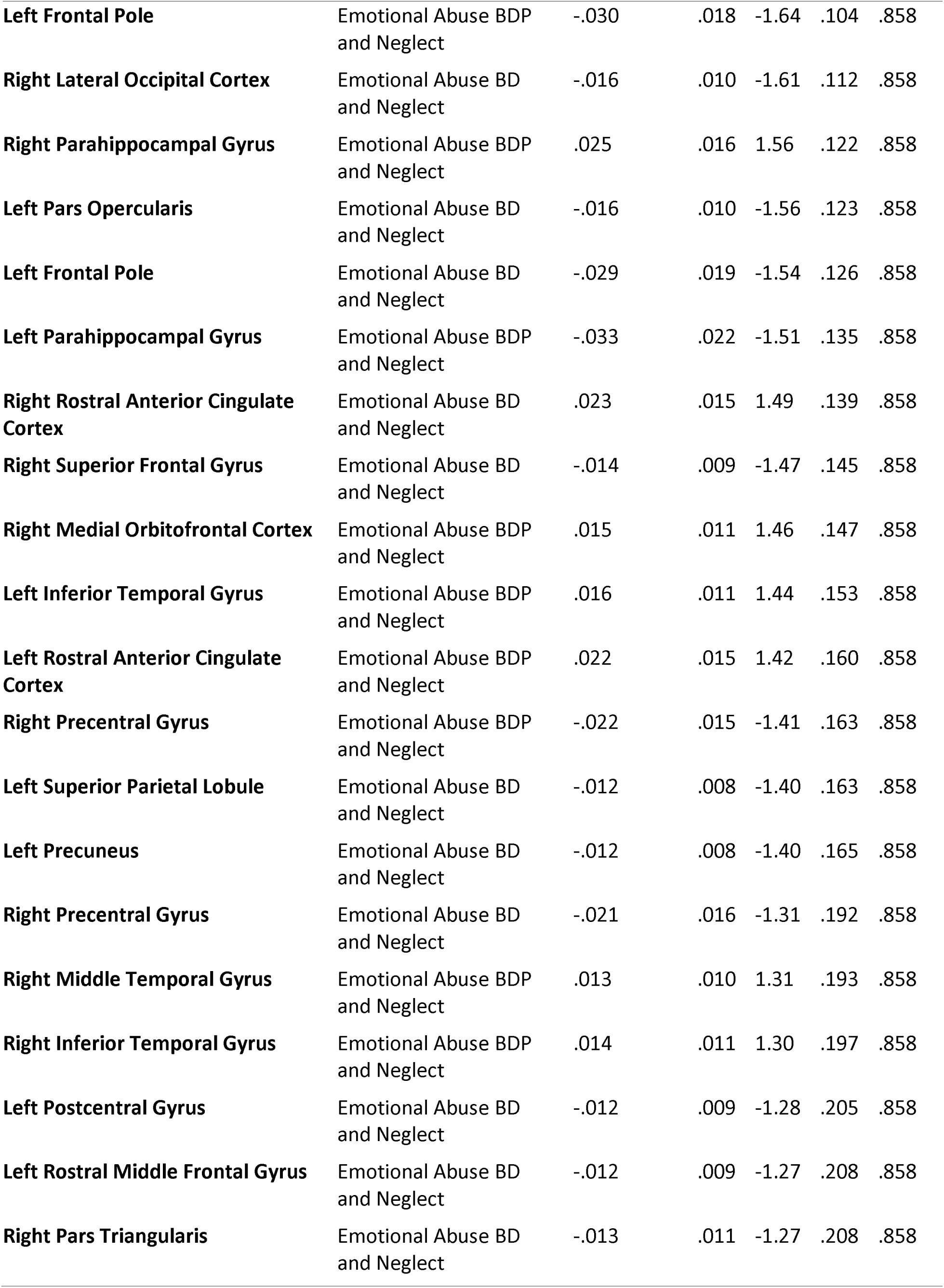
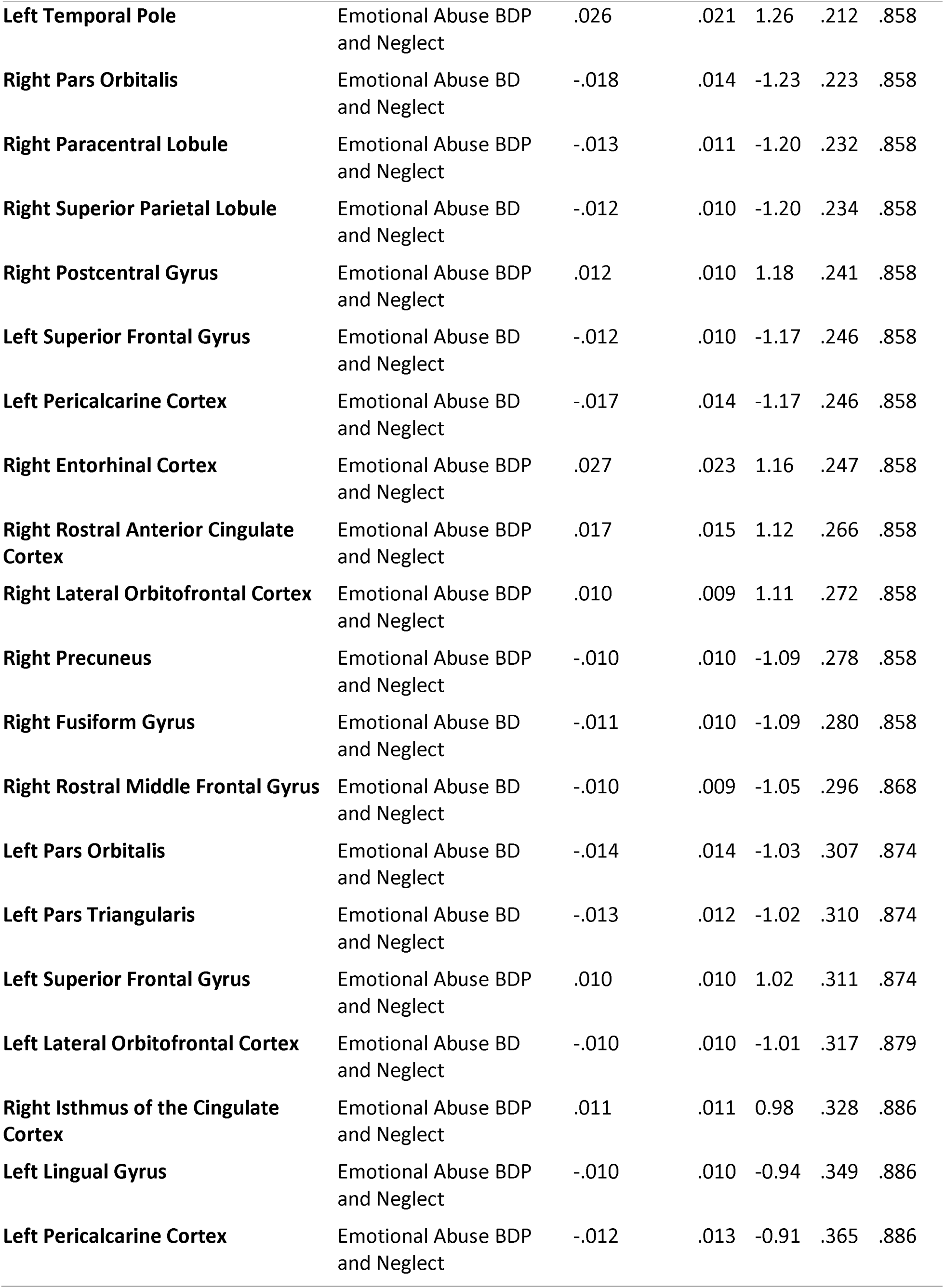
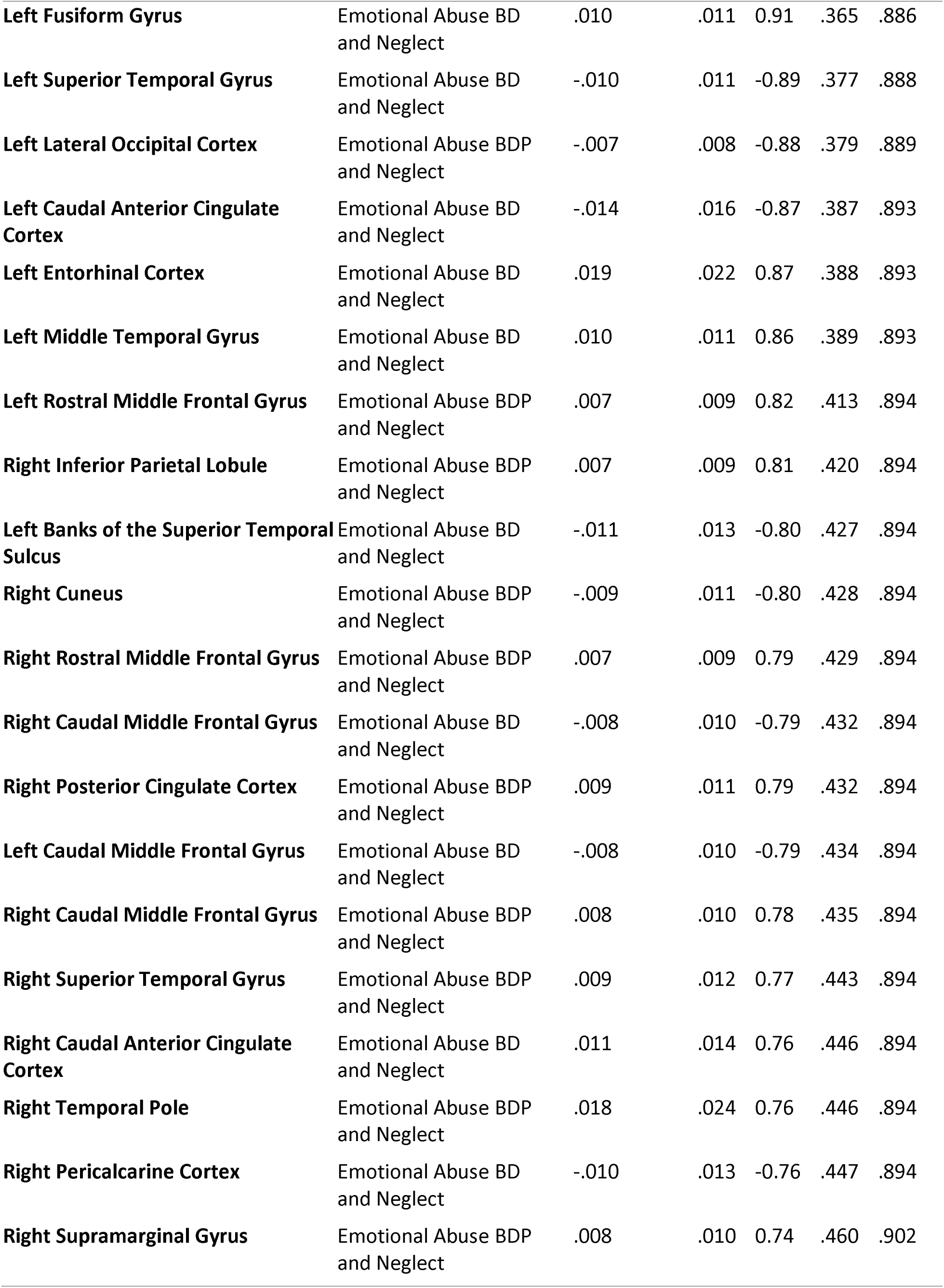
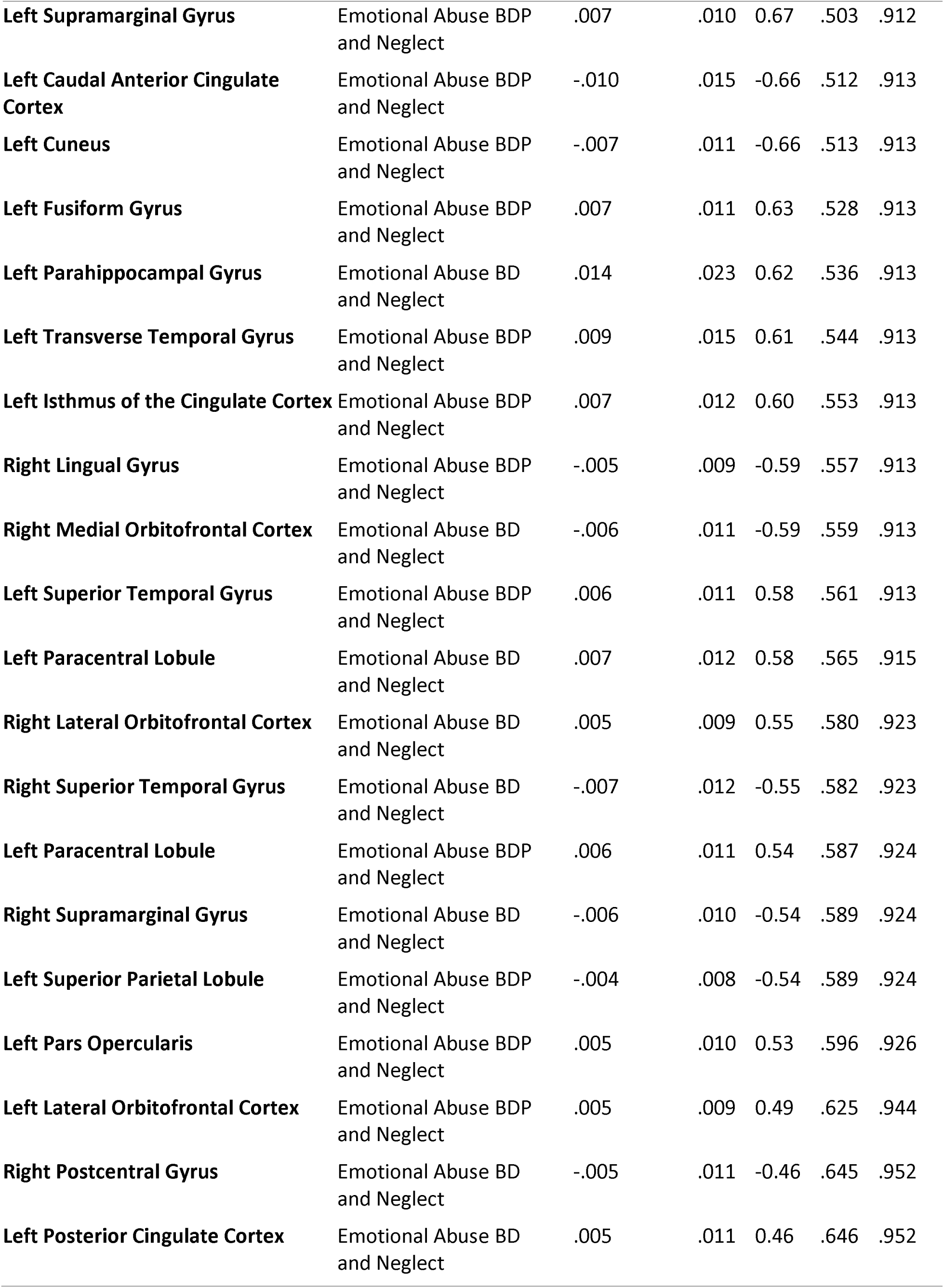
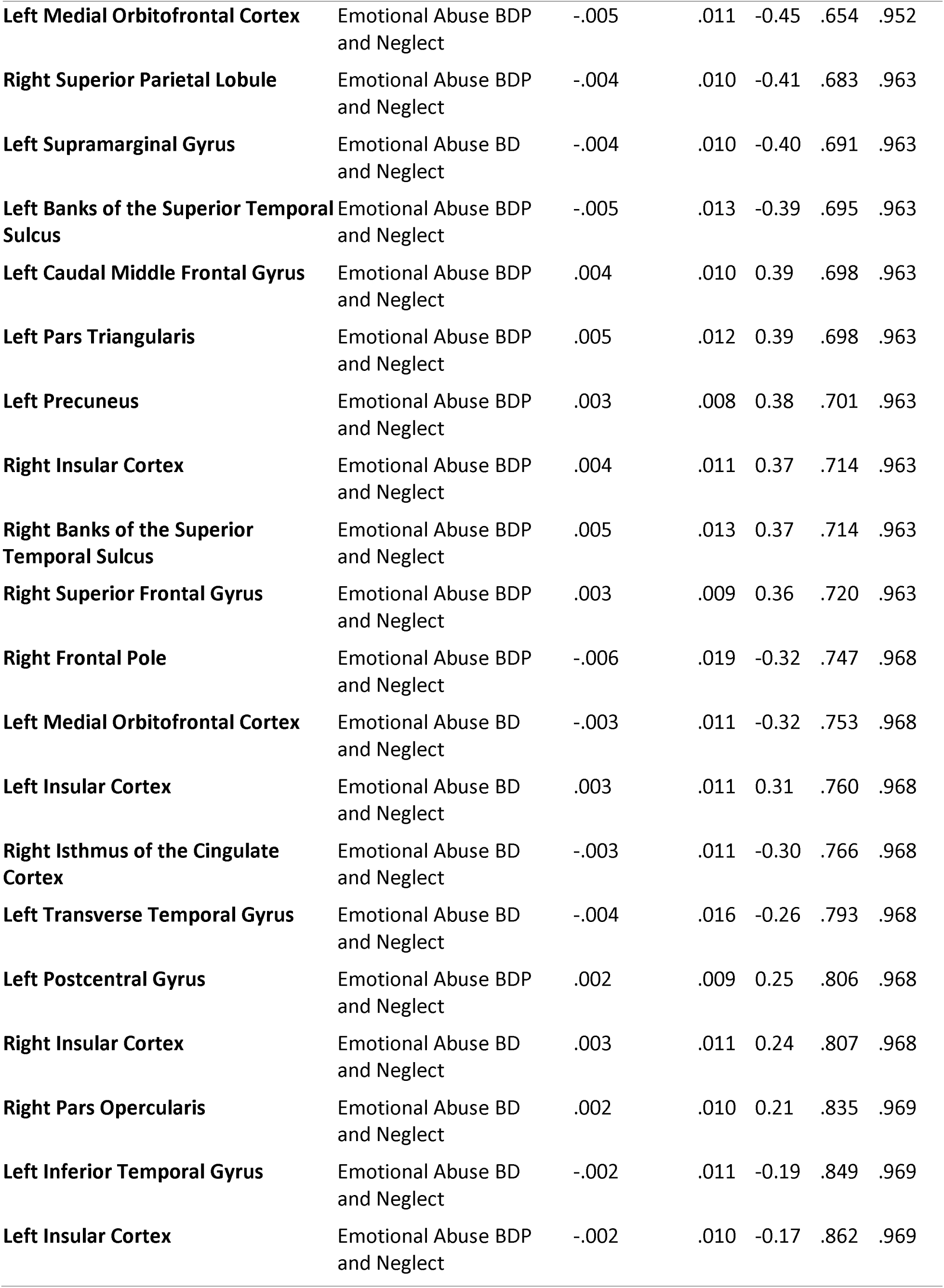
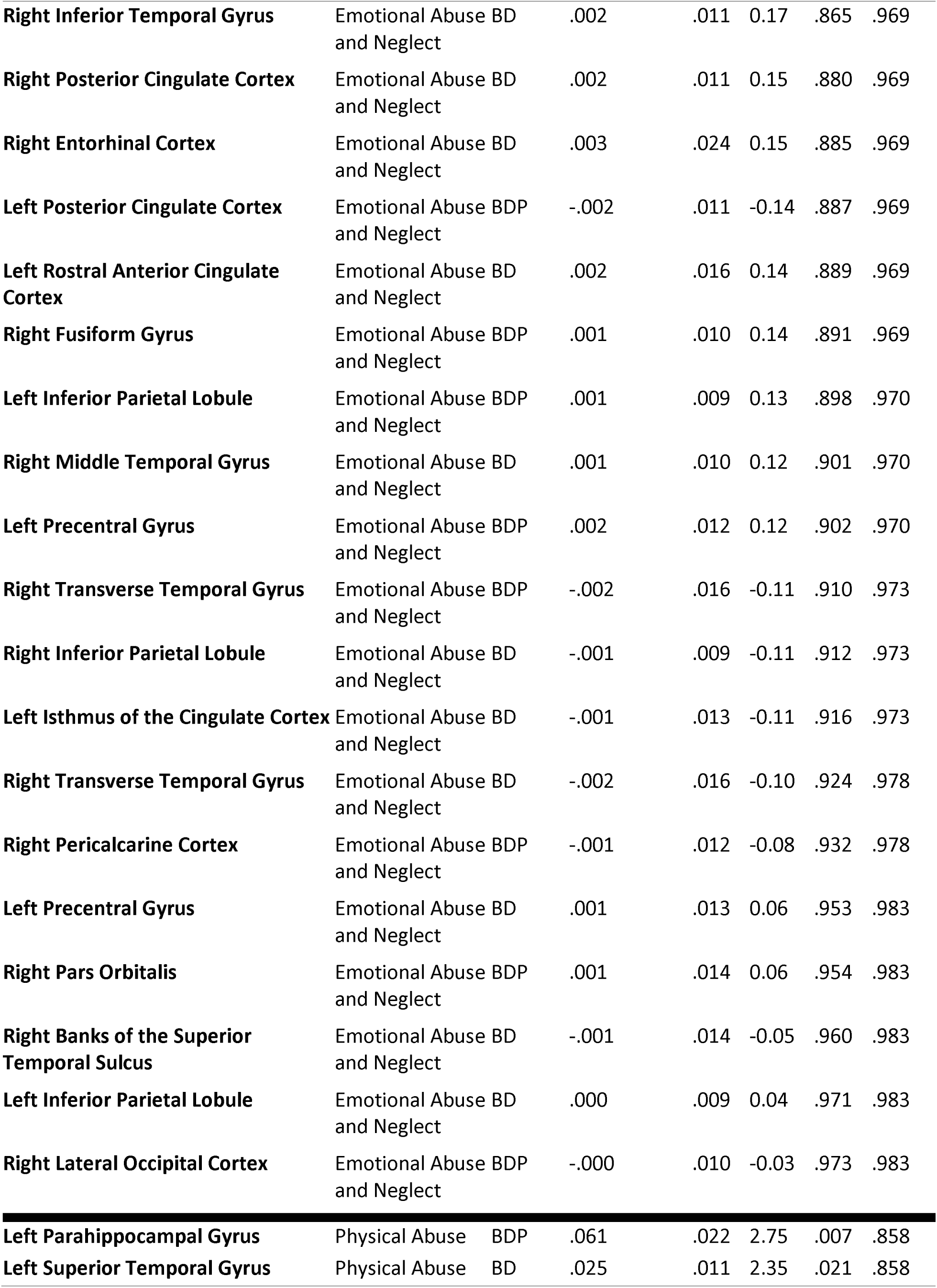
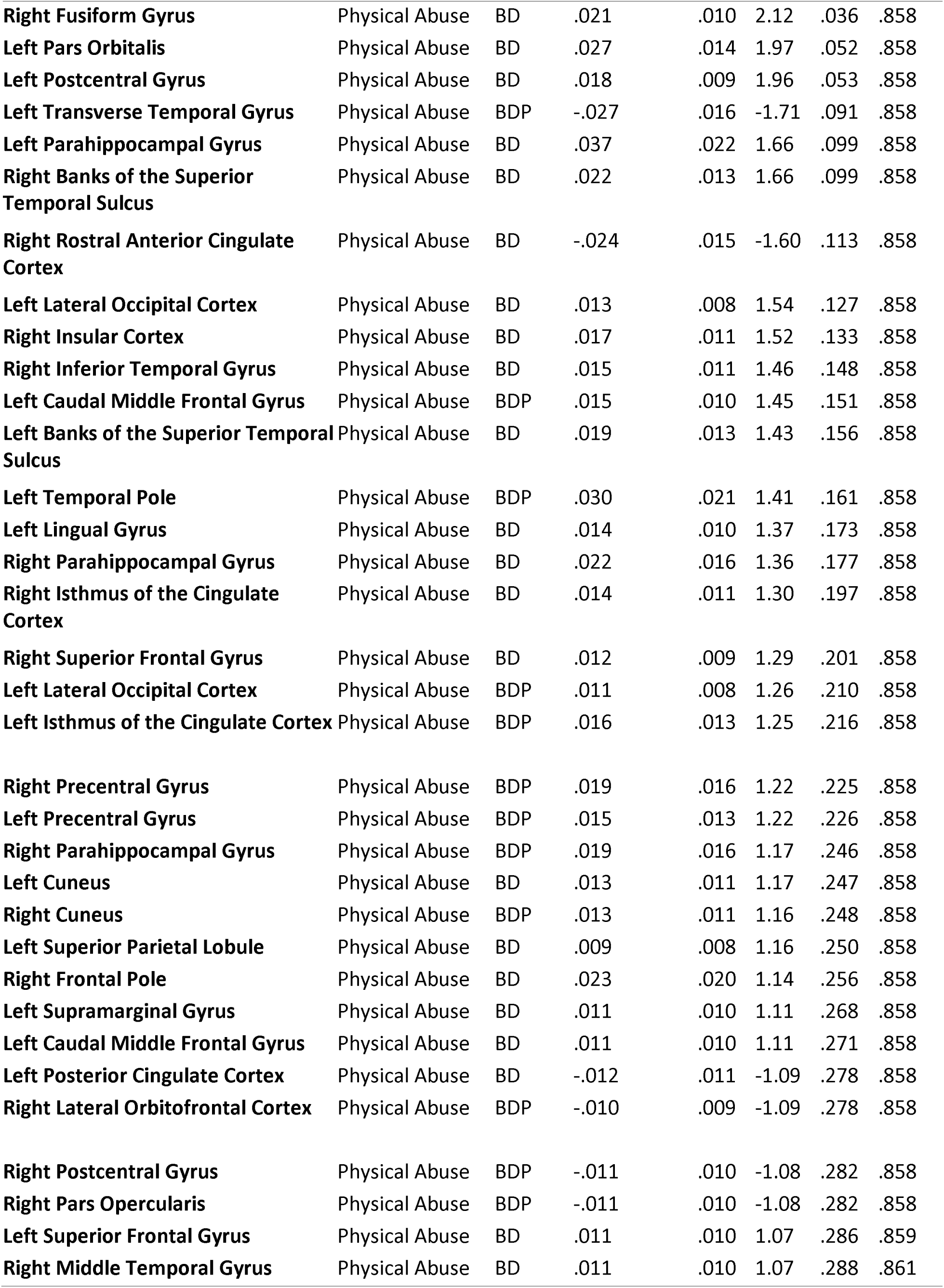
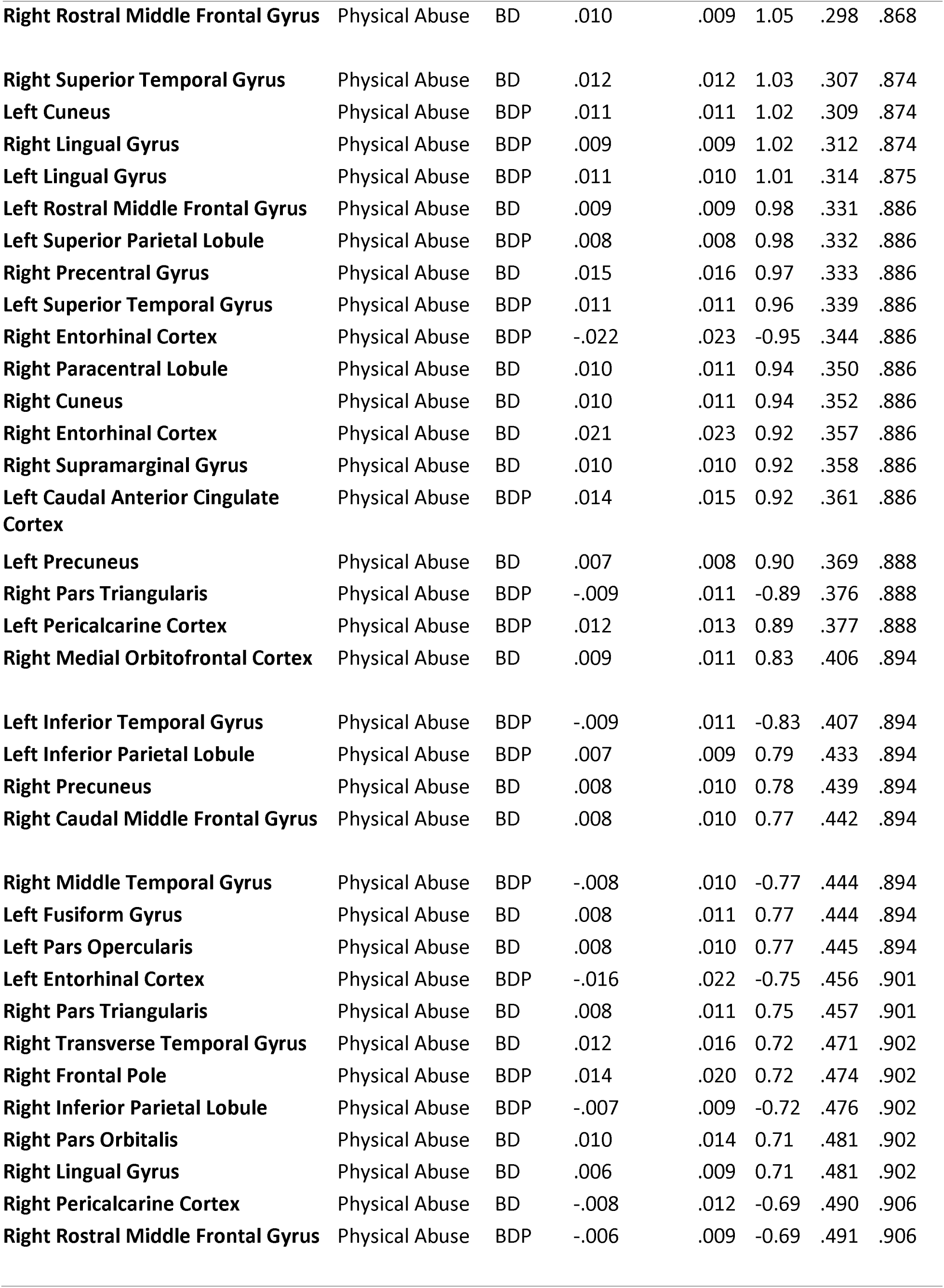
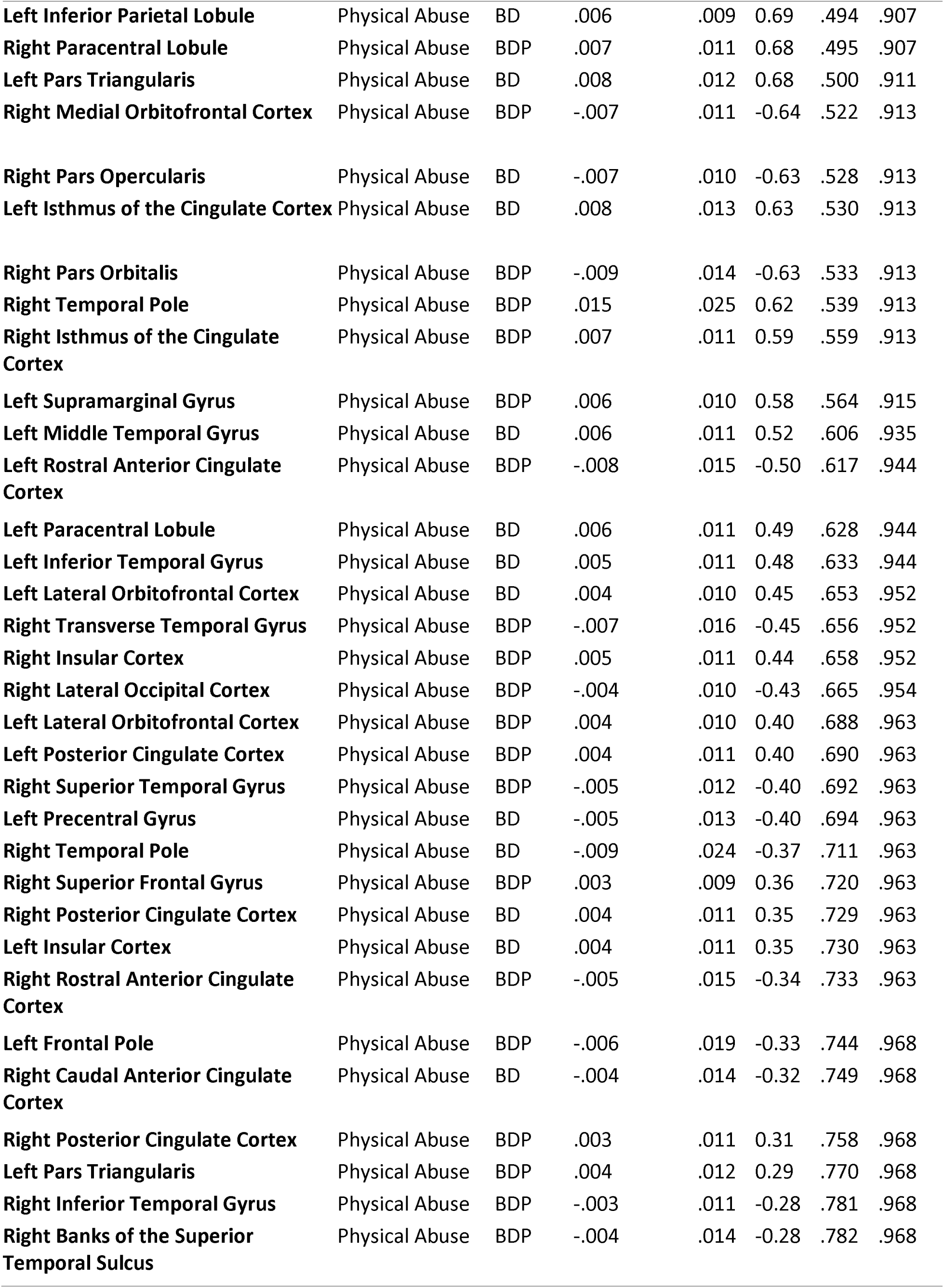
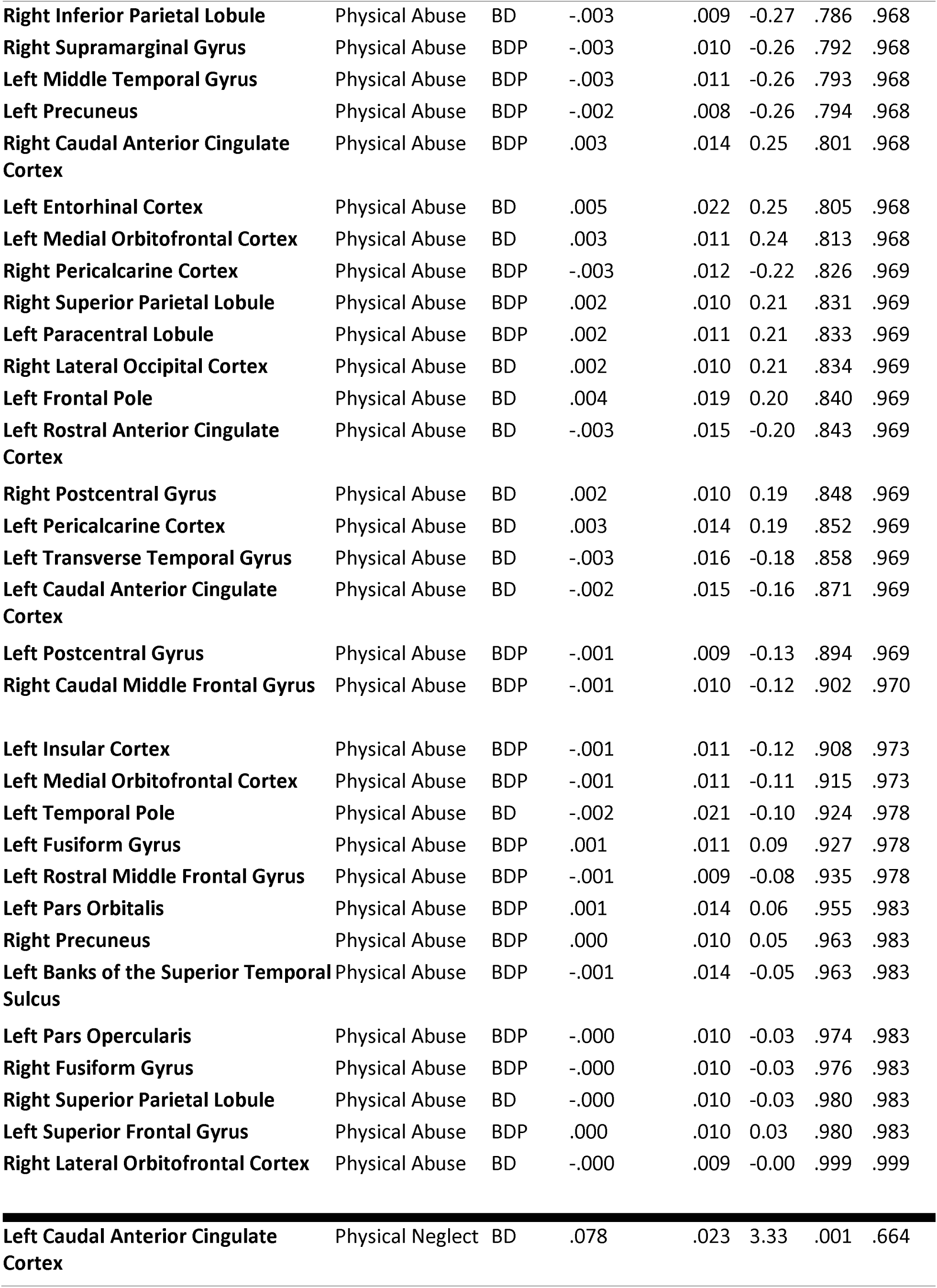
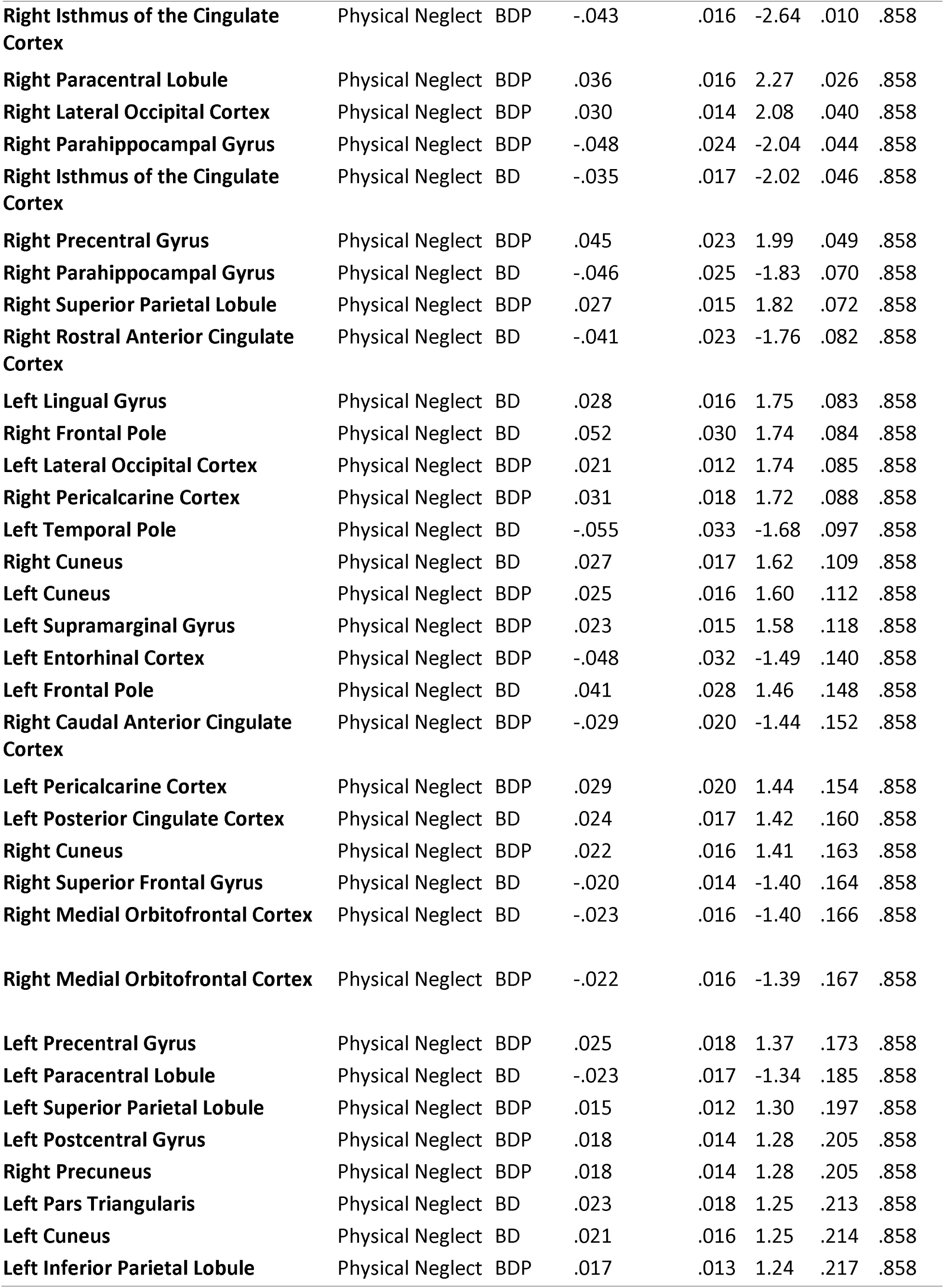
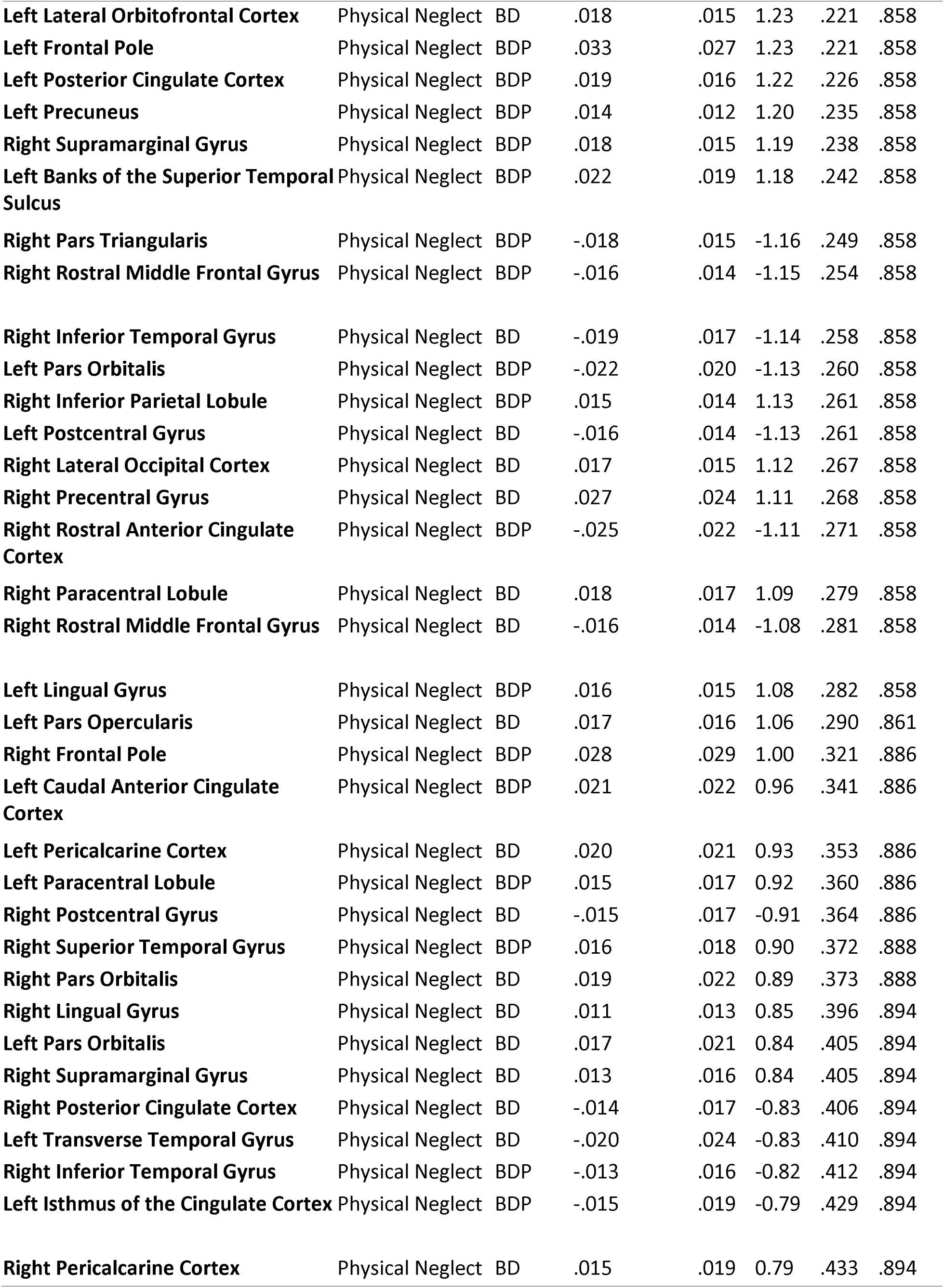
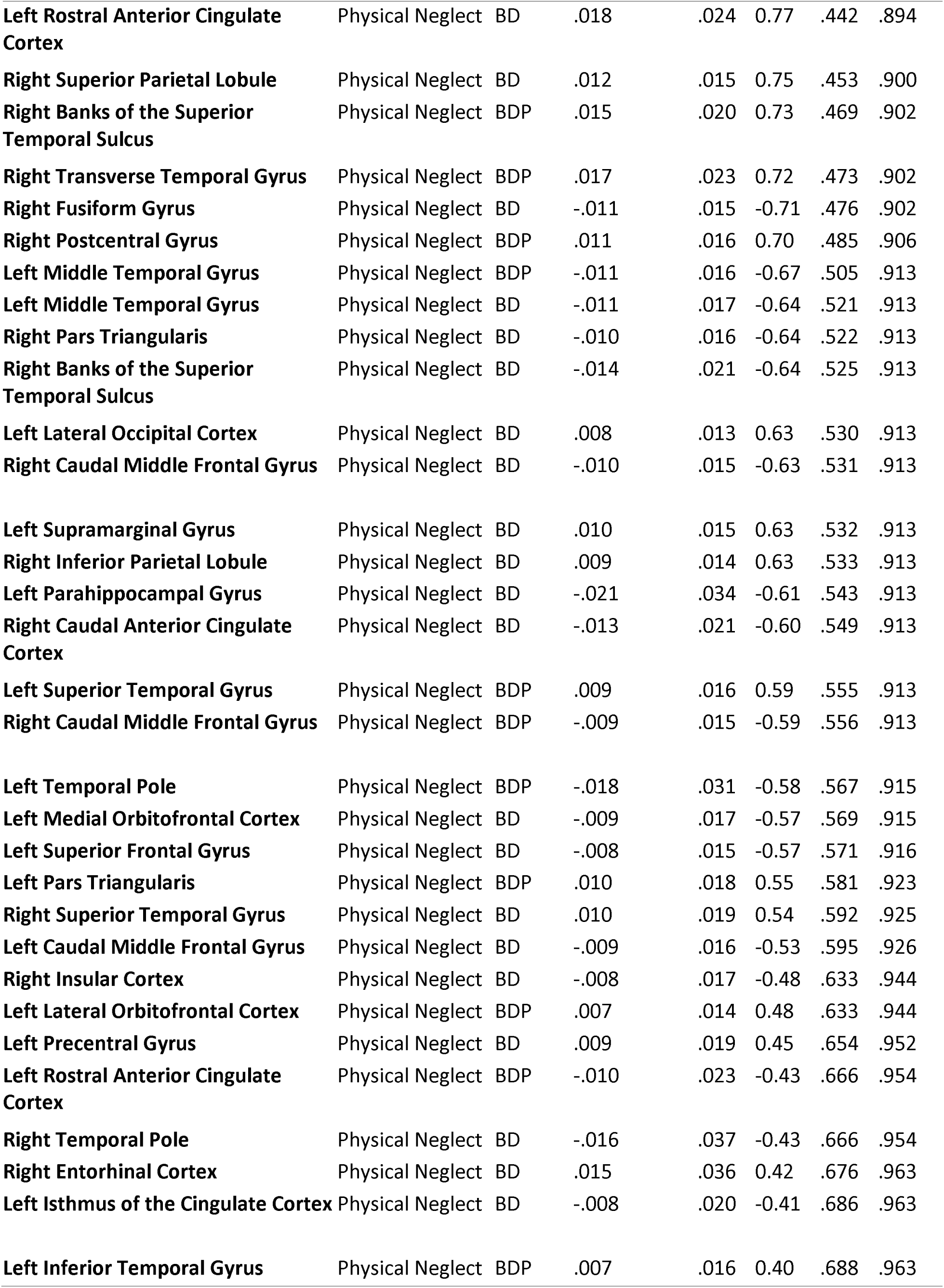
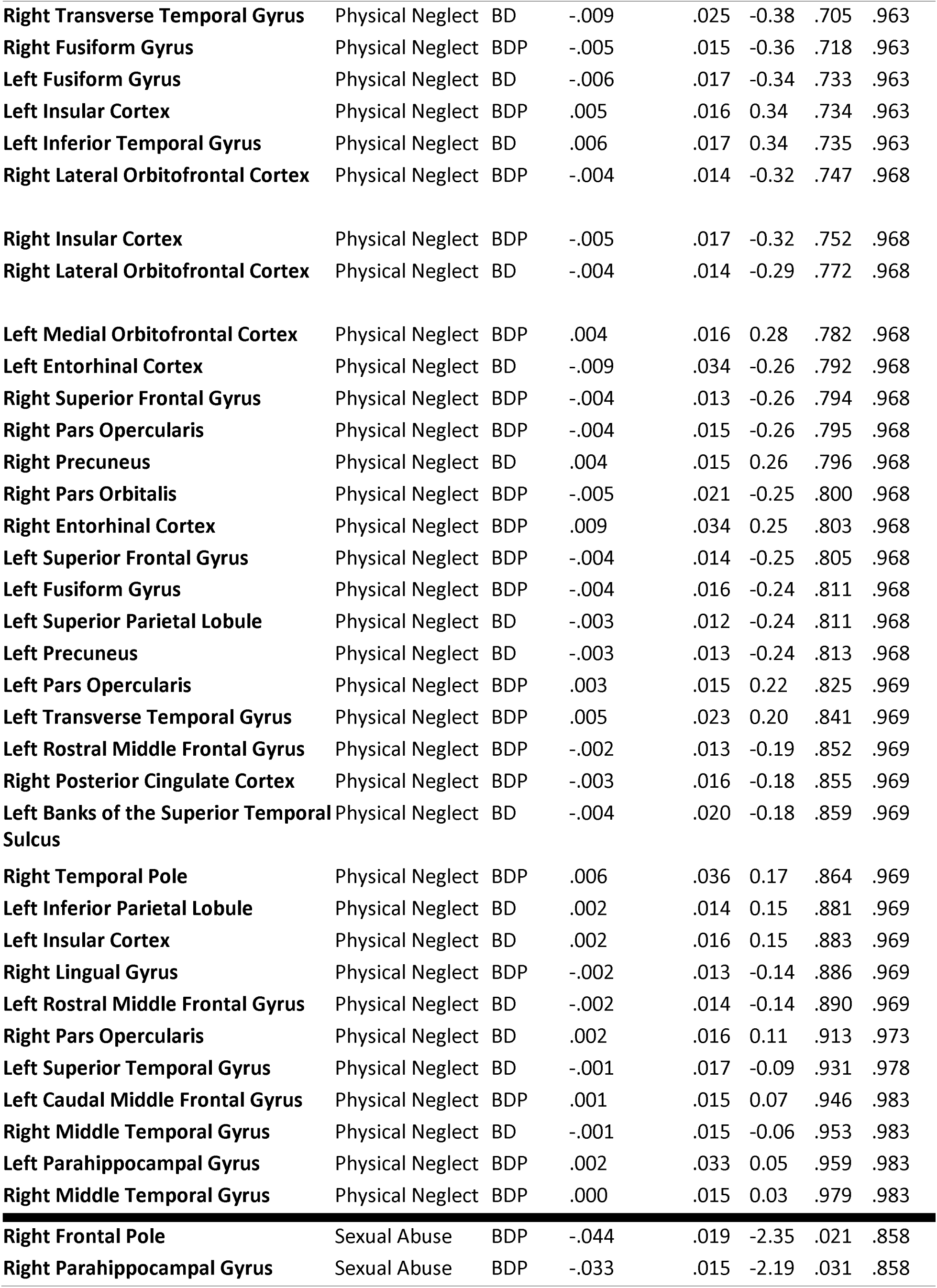
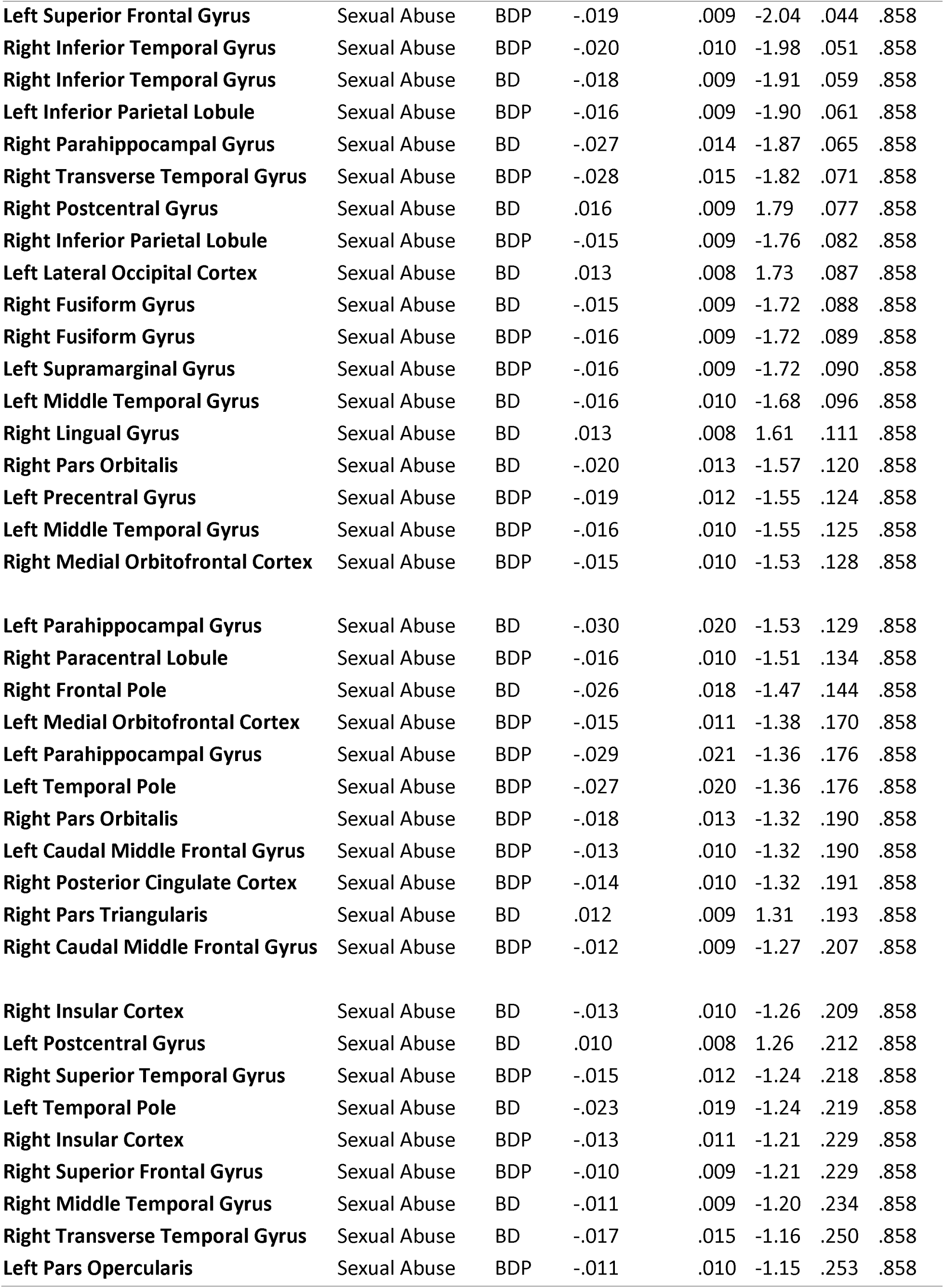
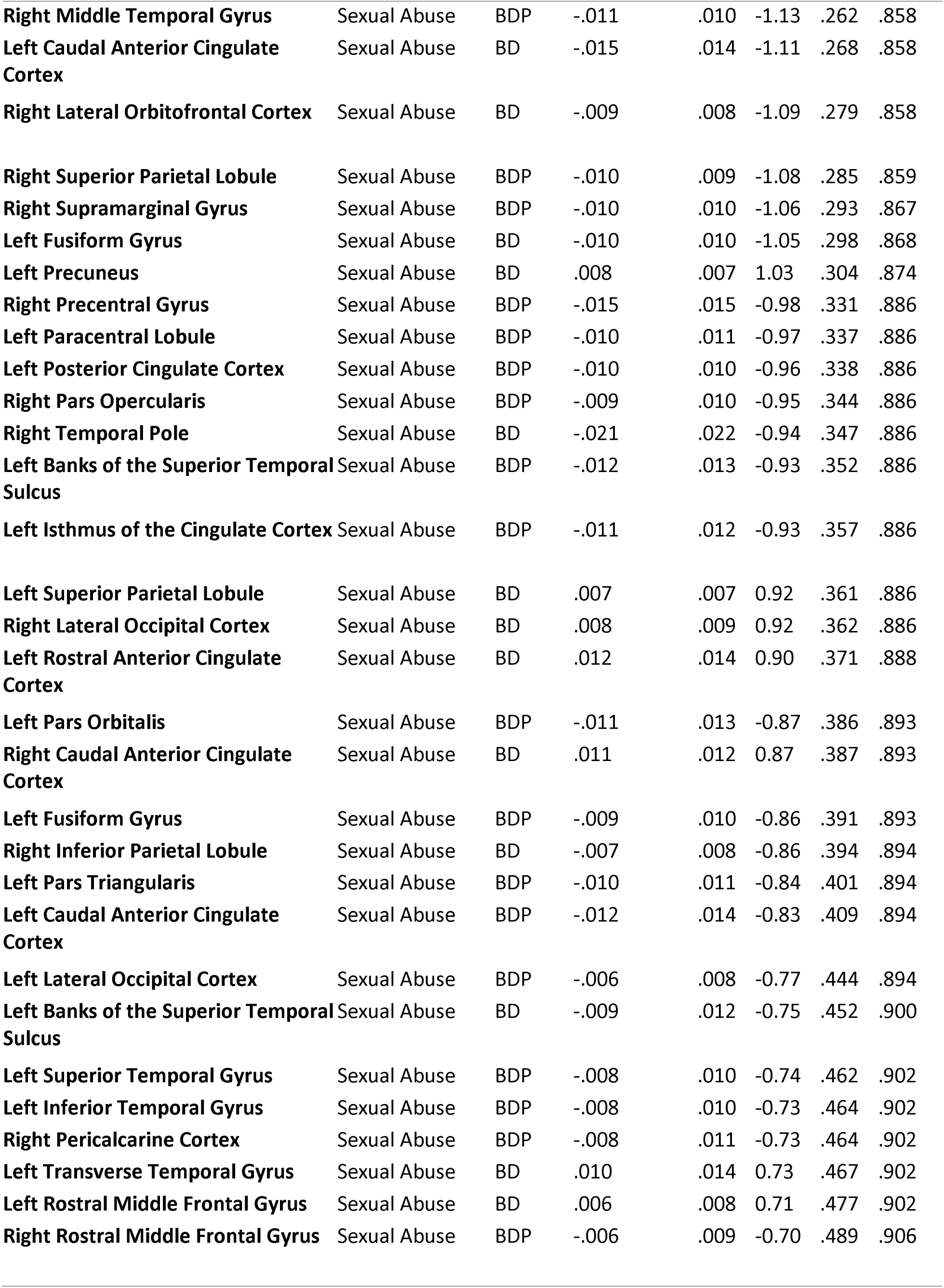
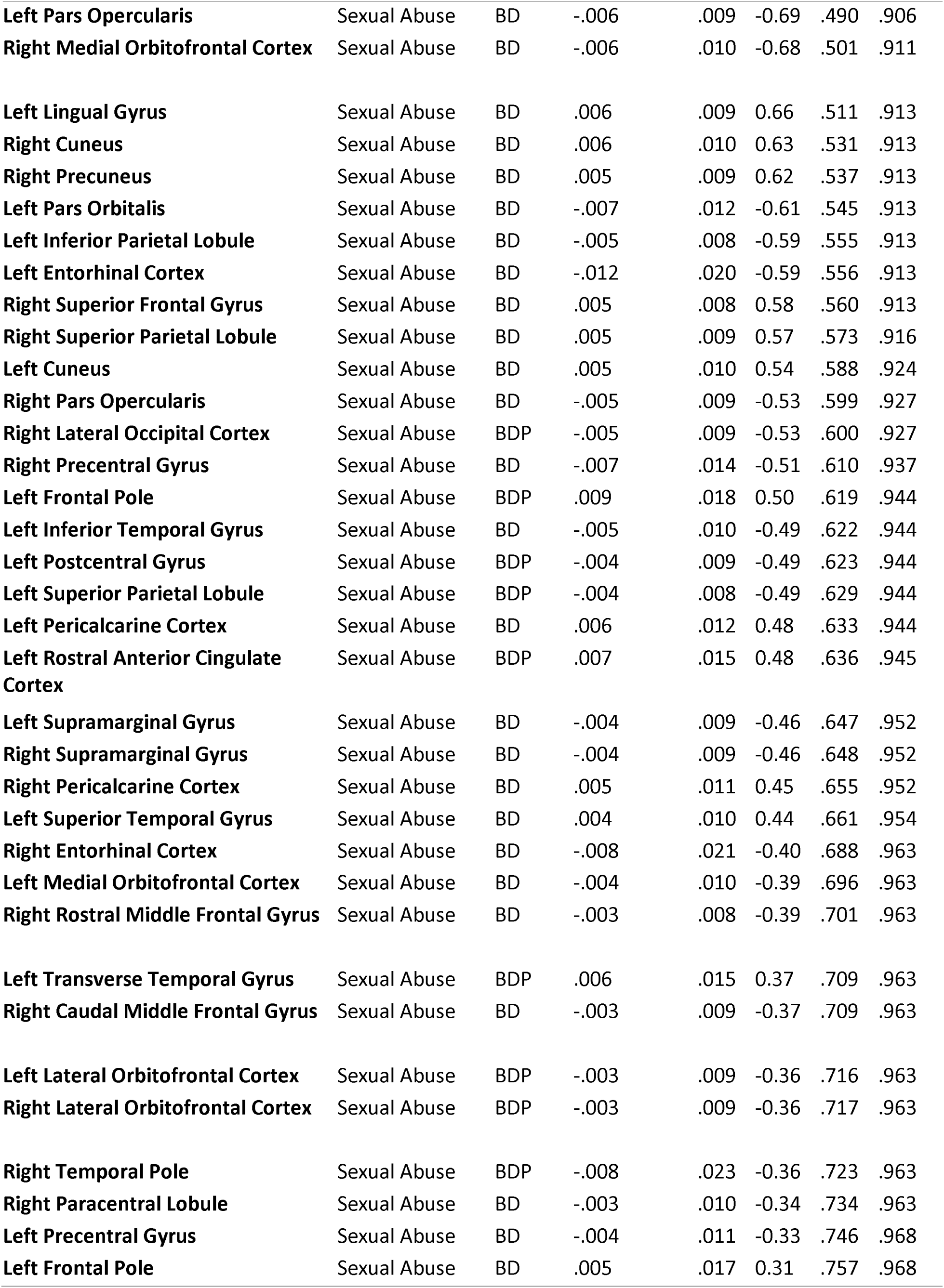
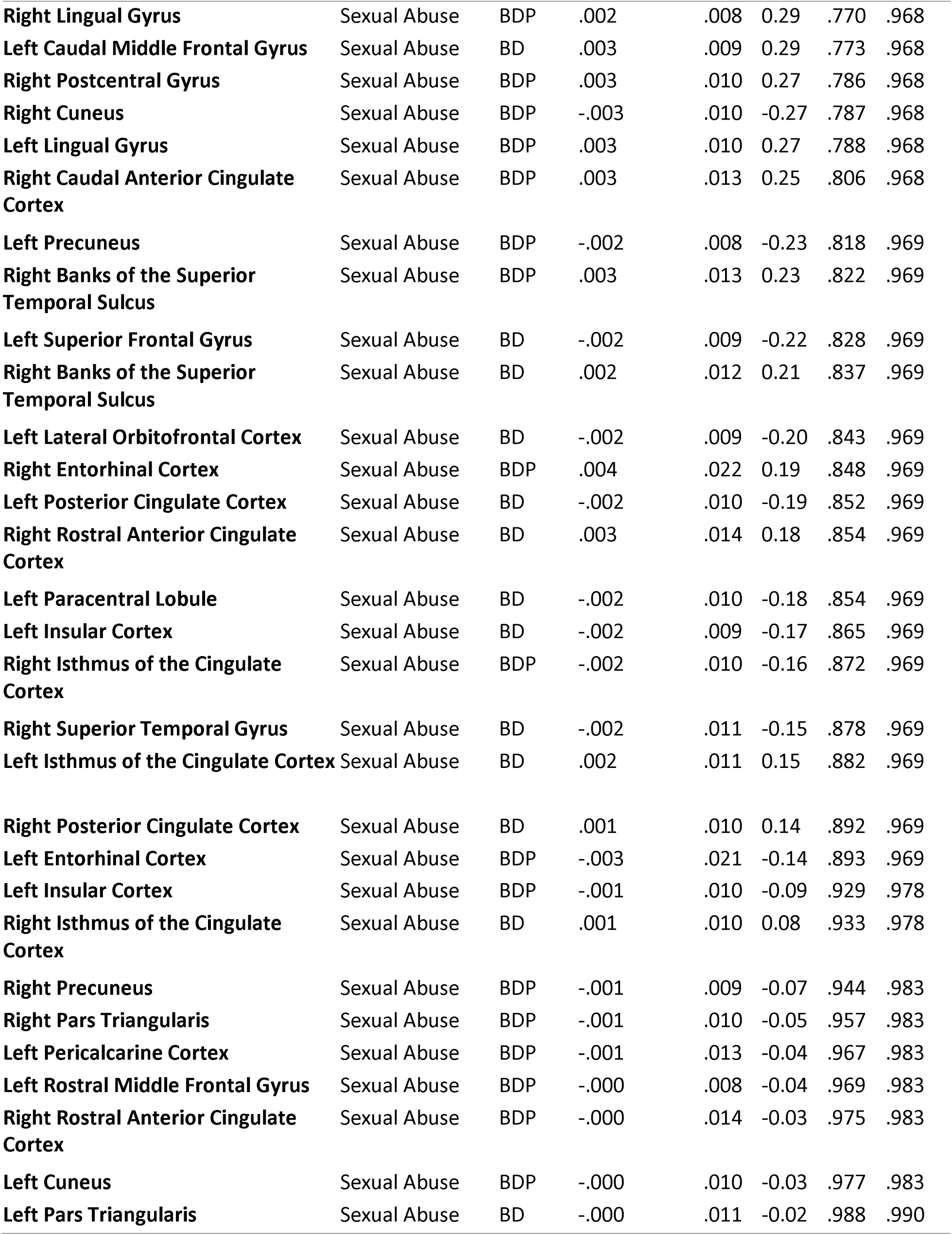
Trauma x Diagnostic Group Interaction Effects on Cortical Thickness.

